# RNA promotes synapsin phase separation providing a platform for local translation

**DOI:** 10.1101/2025.01.29.635107

**Authors:** Branislava Rankovic, Zachary M Geisterfer, Akshita Chhabra, Vladimir M Jovanovic, Ian Seim, Christian Hoffmann, Antonela Condric, Gerard Aguilar Pérez, Rinse de Boer, Lars T. Hofstede, Kiara Freitag, Katja Nowick, Marina Jendrach, Frank Heppner, Vladimir Despic, Michaela Müller-McNicoll, Geert van den Bogaart, Sofiia Reshetniak, Amy S. Gladfelter, Dragomir Milovanovic

**Author notes:** These authors contributed equally (Branislava Rankovic, Zachary M Geisterfer). These authors contributed equally (Akshita Chhabra, Vladimir M Jovanovic, Ian Seim, Christian Hoffmann).

## Abstract

Condensates at synapses organize synaptic vesicle (SV) clusters and are essential for efficient neurotransmitter release. While it is established that RNA granules traffic along the axons, the function of RNA at the presynapse remains unclear. Here, we uncover a direct structural role of coding RNAs in organizing presynaptic condensates by focusing on SV clusters, condensates between synapsin-1 and lipid vesicles, a defining feature of nerve terminals. Using in vitro reconstitution systems, we show that RNA drives synapsin-1 coacervation, with bias toward structured RNAs being more effective at promoting phase transitions. The importance of RNA was confirmed in living synapses, where acute disruption of native RNA induces a dispersion of SVs and synapsin. Conversely, ectopically expressed SV-like condensates have the ability to recruit the translational machinery. The microscopy-based in vitro translation assay demonstrates increased translation efficiency within synapsin-1/RNA condensates. Together, our work indicates a novel structural role of RNAs in modulating SV condensates.

## 1. Introduction

Brain functioning critically relies on efficient neuronal communication. This is achieved via defined compartments—synapses, where chemical neurotransmitters are released from presynaptic axonal boutons to postsynaptic terminals of a dendrite ^[1]^. As highly polarized, long-living and non-dividing cells, neurons require constant remodeling of the local proteome, which is tightly controlled in a spatial and temporal manner ^[2][3]^. In response to environmental cues, protein synthesis occurs in soma ^[4][5][6]^, dendrites ^[7][8]^ and axons ^[9][10][11][12]^ including pre- and post- synapse ^[13][14][15]^, axonal growth cones ^[16][17]^ and on moving membranous organelles such as late endosomes and lysosomes ^[18][19]^.

For effective local translation, mRNA molecules and translational regulators are packaged into RNA granules, membrane-less compartments that form by the biomolecular condensation of proteins and nucleic acids ^[20][21]^. This form of organization allows for the coordinated transport to and translation of mRNAs at designated locations in the cell ^[22][23][24]^. The failure in assembly and trafficking of RNA condensates is implicated in neurodegenerative diseases ^[25][26][27][28][29].^

Despite the clear relevance of targeted mRNA trafficking and the controlled local translation in axons, several crucial questions about spatial control of translation remain open. How are presynaptic RNAs, ribosomes, and translational components sequestered and maintained within the presynaptic boutons? What is the impact of RNAs and translation machinery on the organization of presynaptic boutons, in particular the cluster of synaptic vesicles (SVs), a defining feature of presynapse and an example of biomolecular condensate composed of hundreds of SVs ^[30]^? Finally, it is unclear whether the crowded environment of presynapse is permissive for an augmented local translation.

Here, we discover that synapsin-1 phase-separates with RNAs and that synapsin-1/RNA condensates sequester SVs. Interestingly, the secondary structure and length of both non-coding ^[31][32]^ and messenger (m)RNAs ^[33][34][35]^ modulate assemblies of RNA granules. By establishing opto-chemical manipulations in living neurons, we further show that the acute disruption of RNAs in situ disperses SVs from the boutons. Moreover, by adapting an ex situ translation assay, we show that synapsin-1 condensates are permissive for the translation of a client mRNA, further supporting biomolecular condensates not only as attenuators but as activators of translation ^[36][37][38]^. Together, our work provides evidence of a role for RNAs in modulating SV condensates at the synapse.

## 2. Results

### 2.1 RNA promotes the biomolecular condensation of synapsin-1

Biomolecular condensation can be driven through interactions between oppositely charged molecules, a process known as complex coacervation ^[39]^. As synapsin-1 is one of the most abundant, highly charged proteins (pI 12.4) in the presynapse ^[40]^, we were interested to see if it could undergo biomolecular condensation with RNA (**Fig. 1A**). We incubated recombinant EGFP-tagged synapsin-1 (synapsin-1) and total neuronal RNA across a range of concentrations. Importantly, these mixtures did not include any additional molecular crowder. Indeed, we observed that synapsin-1 was able to undergo phase separation in vitro in the presence of RNA (**Fig. 1B, C and S1**). Similar to previously reported synapsin-1 homotypic protein condensates ^[41]^, these synapsin-1/RNA condensates are dynamic, and show a 65 % recovery of mobile fraction within minutes after photobleaching (**Fig. 1C**). In addition, these RNA-driven synapsin-1 condensates retained their property to recruit SVs into their phases (**S2**). To assess if the synapsin-1 condensates require neuronal RNA to be maintained (**Fig. 1B, C**), we added RNase A upon formation of synapsin-1/RNA condensates and followed its effect on the condensates over the course of 10 min (see Methods) (**Fig. 1D, S3A, B**). Upon RNase A treatment, we observed a decrease of synapsin-1 signal within the condensates (**Fig. 1D)**, and in some cases, the complete dissolution of condensates (**Fig. 1D**, RNase A 1 µg/µl treatment). The significant reduction in synapsin-1 fluorescence enrichment was observed, both with concentration of 1 µg/µl and 0.1 µg/µl of RNase A treatment. (**Fig. 1E**). Moreover, synapsin-1 fluorescence intensity signal increased in a dilute phase after the RNase A treatment (**S3C**). Similarly, RNA pre-digested with RNase A was incapable of supporting the condensation of synapsin-1 (**S3D**) indicating the polymeric form of RNA is required. These results suggest that RNA, in the absence of molecular crowders can promote the condensation of synapsin-1 in vitro, as synapsin-1 alone would not form condensates without RNA, lipid vesicles, or chemical crowders ^[42][43]^.

**Figure 1.**
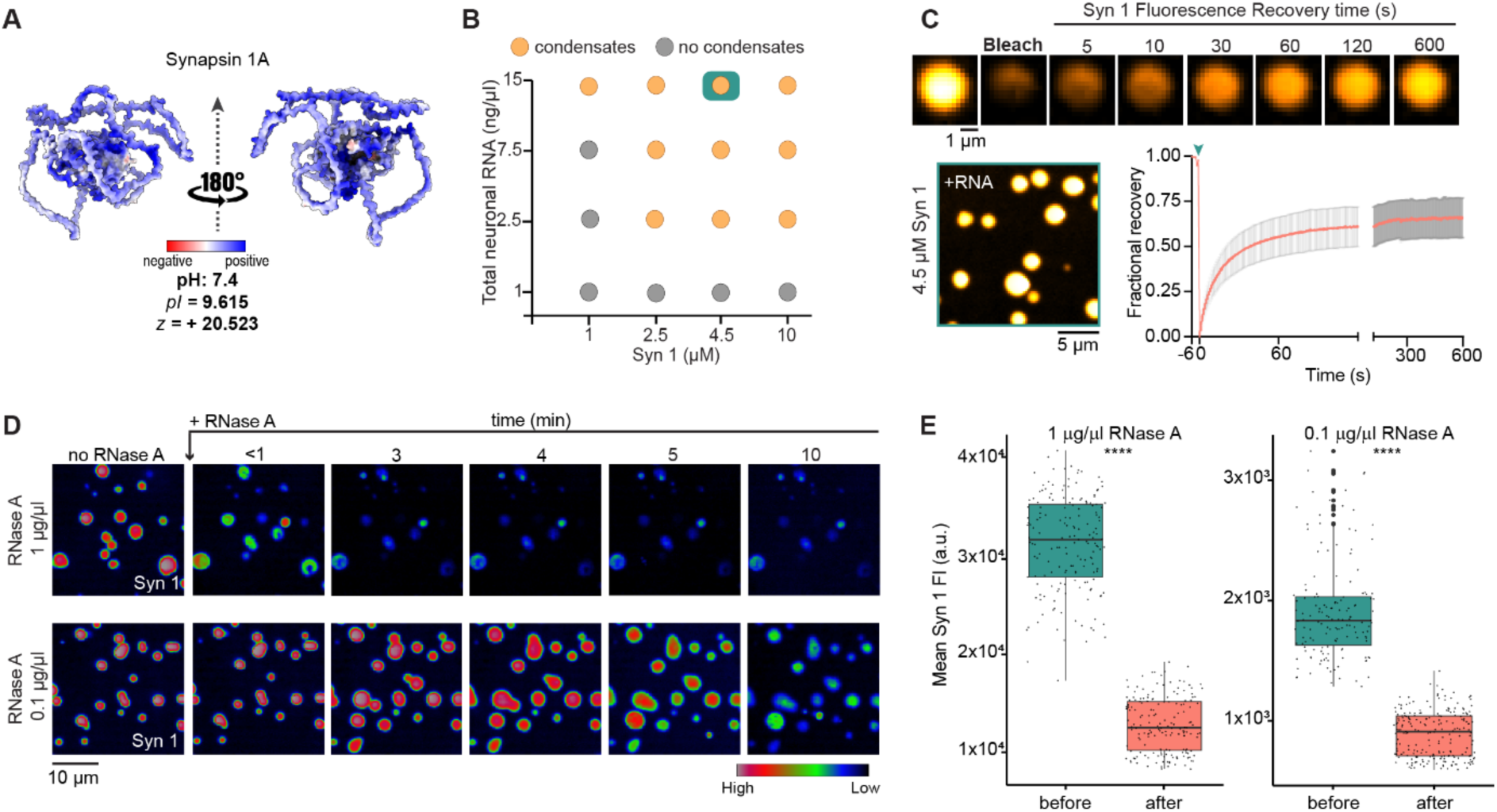
RNA promotes condensation of synapsin-1. **(A)** Charge-based surface model of synapsin-1 (*Rattus norvegicus*; UniProt ID P09951) showing charge distribution. EGFP-tagged rat synapsin-1 (Syn 1) was purified for in vitro reconstitutions. **(B)** Summary of phase-separation behavior of synapsin-1 in the presence of total RNA from primary mouse neurons, with increasing concentrations of both protein and RNA without any molecular crowder in 10 µl-reaction buffer (150 mM NaCl, 25 mM Tris-HCl at pH 7.4, 0.5 mM TCEP). Corresponding images are shown in the S1. **(C)** FRAP of reconstituted synapsin-1 condensates (4.5 µM) with 150 ng of total neuronal RNA. Data was collected from two independent in vitro reconstitutions. Error bars in FRAP curve plot represent standard deviation (± SD). The mobile fraction of synapsin-1 was 65 %. Teal arrow indicates the moment of the bleach. Representative images show pre-bleach, bleaching and recovery of synapsin-1 fluorescence in the course of 600 seconds. **(D)** Representative images of selected RNA-driven synapsin-1 condensates treated with 1 µg/µl or 0.1 µg/µl final concentration of RNase A for 10 min. Fluorescence intensity signal in gray values (see histogram legend) indicates fluorescence intensity signal of Syn 1 before and at different time points upon the addition of RNase A to pre-formed synapsin/RNA condensates with 10 µM synapsin-1 (to a final concentration) and 300 ng of total neuronal RNA. **(E)** Quantification of synapsin-1 fluorescence enrichment in not treated RNA-driven synapsin-1 condensates (before) and 10 min after the treatment (after) with 1 or 0.1 µg/µl of RNase A. The jitter box plot encompasses 25^th^ and 75^th^ quartiles with median marked by central line. Minimum and maximum of quantified synapsin-1 mean fluorescence intensities (FI) are denoted by whiskers. Black circles (above the whiskers) represent mean fluorescence intensities outside the range of adjacent values. Each point represents a value from one condensate, which was slightly shifted for better visualization of overlapping data points. **** indicates P ≤ 0.0001 by Wilcoxon rank sum test. Data was collected from three independent in vitro reconstitutions.

### 2.2 RNA structure modulates the stoichiometry of RNA and synapsin-1 within condensates

Recent evidence suggests RNA structure, in addition to charge and primary sequence, can control the formation and specificity of protein-RNA condensates ^[44][45]^. To understand the contribution of RNA structure in the formation of synapsin condensates, we utilized an orthogonal mRNA, *CLN3*, designed in silico to either sample many different conformations akin to an intrinsically disordered region (IDR) in proteins (high ensemble diversity (ED), called *CLN3* high-ED) or sample relatively few different conformations (low ED, called *CLN3* low-ED), while preserving the total nucleic acid content and length ^[46]^. Two sequences were chosen representing these different classes of conformational heterogeneity based on their minimum free energy and ED (**Fig. 2A**) ^[47][48]^. These RNAs were added to reactions containing synapsin-1 in a 1:1 polyelectrolyte charge ratio and allowed to form condensates. For both *CLN3* structure mutant systems, we saw the immediate formation of synapsin-1/RNA condensates (**Fig. 2A**). Notably, the partitioning of synapsin-1to the condensates varied based on RNA structure (**Fig. 2B**). Condensates formed with RNAs that have low conformational heterogeneity (*CLN3* low-ED) tended to be more protein-rich, as opposed to condensates formed in the presence of RNAs with more diversity in possible structures (*CLN3* high-ED). (**Fig. 2B**). While full-length synapsin-1 was able to form condensates with both kinds of RNAs, the synapsin-1 IDR (421-705) was only able to form condensates with more conformationally heterogeneous RNAs (**S4**). These results suggest that RNA structure heterogeneity, in addition to charge, is an important feature in regulating the formation of synapsin-1/RNA condensates.

**Figure 2.**
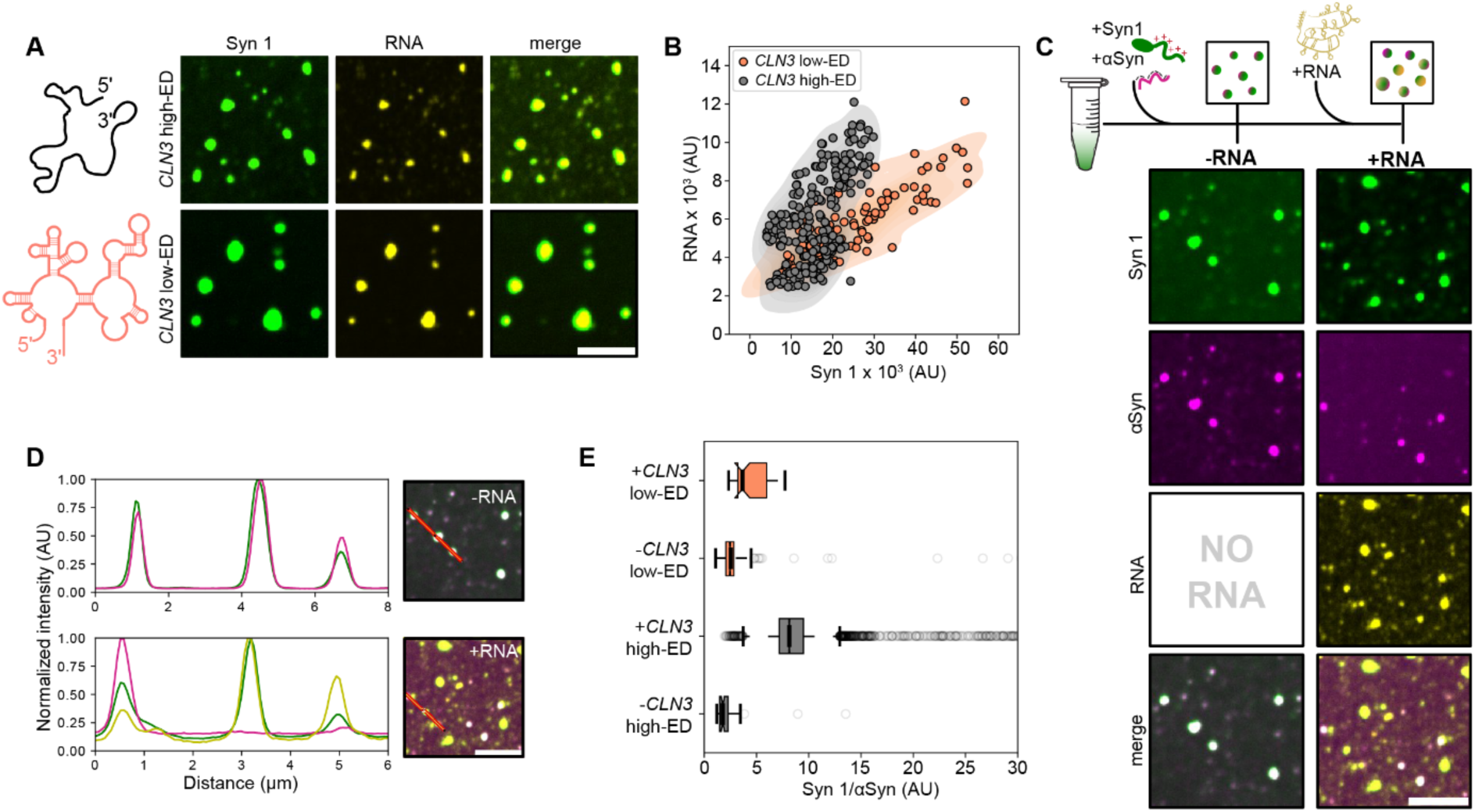
RNA structure modulates the stoichiometry of synapsin-1 within condensates. **(A)** Representative images of in vitro reconstituted EGFP-synapsin-1 (Syn 1) and *CLN3* condensates with high-ED and low-ED Cy5-labelled *CLN3* mutant in the absence of molecular crowder in reaction buffer (150 mM, NaCl, 25 mM Tris-HCl (pH 7.4), 0.5 mM TCEP). Scale bar, 10 µm. **(B)** Scatter plot overlayed with Kernel Density Estimate (KDE) shows fluorescence intensities of *CLN3* RNA (high-ED in grey and low-ED in magenta) as a function of Syn 1 intensities. Each circle represents a single Syn 1/*CLN3* condensate. **(C)** Top: schematic representation of three-component in vitro reconstitution system -type I (see Methods), with purified EGFP-synapsin-1, α-synuclein-AF647 (αSyn) and Cy3 labelled *CLN3* RNAs. Bottom: Representative micrographs of in vitro reconstituted Syn 1/α-Syn condensates without or with *CLN3* RNA. Scale bar, 10 µm. **(D)** Fluorescence intensity line-profiles accompanied with corresponding representative images with selected Syn 1/αSyn ± *CLN3* RNA condensates used to plot the line profiles, where Syn 1 is represented in green, α-Syn in magenta and *CLN3* RNA in yellow. **(E)** Quantifications of Syn 1 and α-Syn fluorescence intensities expressed as a ratio (Syn 1/αSyn) in reconstituted protein-protein droplets with or without low- or high-ED *CLN3* RNAs. The box plot encompasses the 25^th^ and 75^th^ quartiles with the median marked by central line. Whiskers extend to 1.5 times the interquartile range (IQR). Circles indicate outliers. A paired two-tailed t-Test comparing values before and after the addition of RNA indicate a P-value of <0.00001 and 0.00579 for high-ED and *CLN3* low-ED respectively. All data were collected from three independent in vitro reconstitutions.

Each presynapse contains over 300,000 protein molecules ^[40]^, among which the most abundant is synapsin-1. α-Synuclein, another abundant protein in the presynapse that regulates the formation and stoichiometry of synapsin-1 condensates, is a highly acidic and negatively charged protein (pI = -9.73 at pH 7.4) ^[49]^ ^[50]^. We hypothesized that RNA and α-synuclein may compete in their recruitment to synapsin-1 condensates based on their negative charge. To understand how the presence of multiple different anionic polymeric molecules might affect synapsin-1 condensation in vitro, we first pre-formed α-synuclein/synapsin-1 condensates at their physiological stoichiometry ^[40]^. Once the condensates were formed, *CLN3* high-ED or *CLN3* low-ED were added to the reactions and the subsequent shifts in the partitioning of synapsin-1 and α-synuclein were measured. Upon the addition of either *CLN3* high-ED or low-ED, the RNA was recruited into pre-existing α-synuclein/synapsin-1 condensates, in addition to forming synapsin-1/RNA condensates devoid of α-synuclein (**Fig. 2C, D and S5**), which suggested both negatively charged molecules, RNA and α-synuclein, can regulate formation and stoichiometries of synapsin condensates in vitro.

To further understand the impact of RNA on the association of synapsin-1 and α-synuclein, we analyzed the stoichiometry of synapsin-1 and α-synuclein within condensates, calculated as a ratio of intensities of synapsin-1 to α-synuclein before and after the addition of RNA to the reactions (**Fig. 2E**). After the addition of either *CLN3* high- or low- ED, we observed a disruption in the ratio of synapsin/α-synuclein concentrations within synapsin condensates. Such differential enrichments of synapsin and α-synuclein after the addition of RNA suggests that the availability and regulation of RNA in the presynapse may be an important determinant of synapsin condensate stoichiometry in both healthy and disease states.

### 2.3 Synaptic mRNAs modulate α-synuclein recruitment into synapsin-1 condensates

Given the clear impact of RNA conformational heterogeneity in driving the stoichiometry of RNA and protein within synapsin-1 condensates with orthogonal RNAs (**Fig. 2**), we were interested to see whether native synaptic mRNAs of differing structure produce the same effect. To investigate this, we selected two synaptic mRNAs, *Scg3* and *Prnp* from the recently determined transcriptome of mouse primary neurons ^[51]^ with identical lengths but different structural features (*Scg3_ED_*=435, *Prnp_ED_*=635) (indicated in **S6**). We pre-formed protein only condensates containing α-synuclein and synapsin-1 to a final protein concentration ratio, mimicking near-physiological conditions (**Fig. 3A**). After formation of protein-only condensates, *Scg3* or *Prnp* were added to the reactions in a 1:1 polyelectrolyte charge ratio (based on the charge and abundance of synapsin-1) and partitioning of each molecule was analyzed (**Fig. 3B**). RNA recruitment was observed within a few minutes after their addition to the reaction (**S 7A-D**), followed by the recruitment of synapsin-1 but no compensatory recruitment of α-synuclein (**Fig. 3C**), in agreement with what was observed for the reconstitutions with model RNAs (**Fig. 2C-E**).

**Figure 3.**
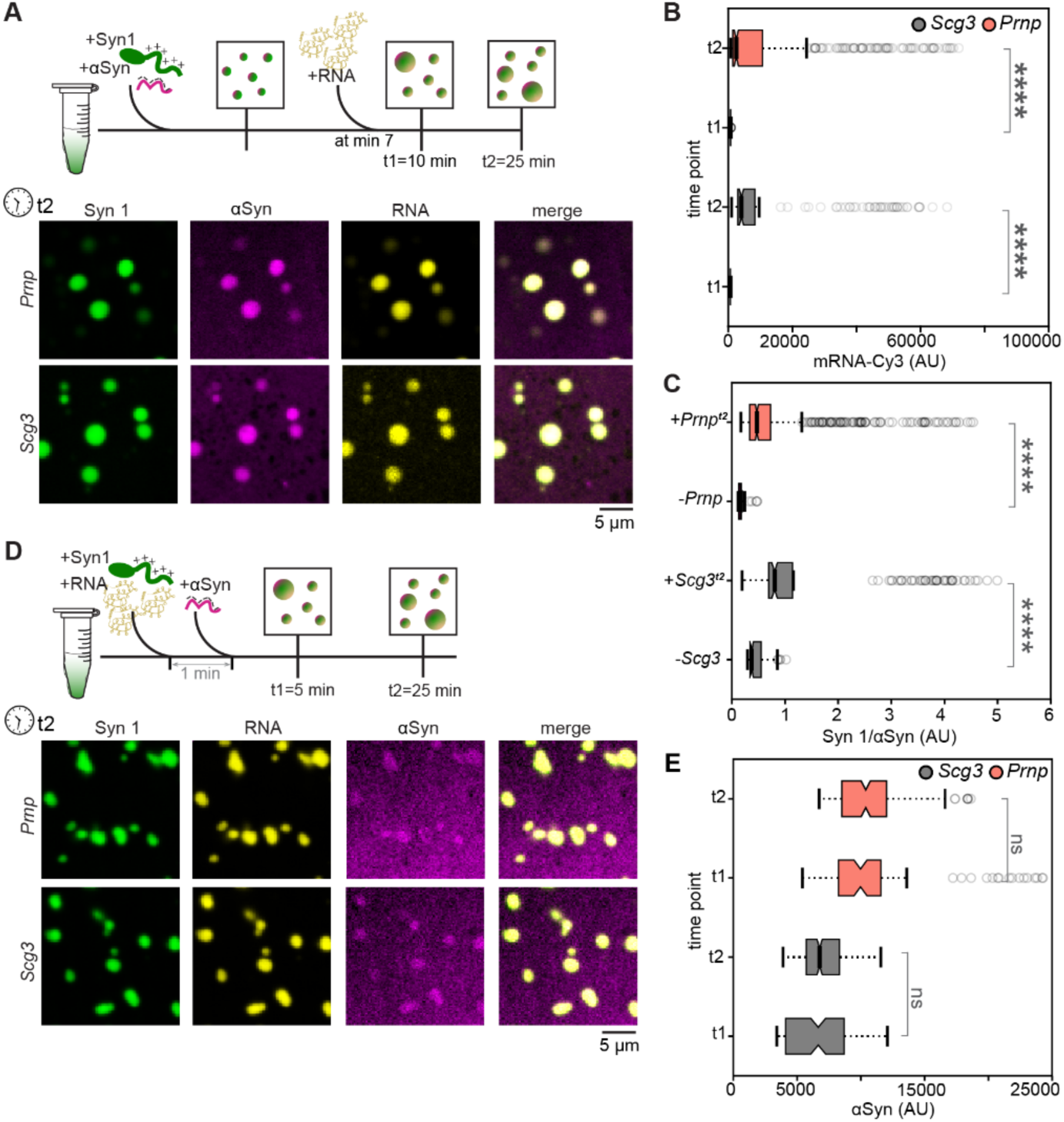
Synaptic mRNAs compete out α-synuclein while maintaining the synapsin-1 stoichiometry within condensates. **(A)** Top: schematic representation of three-component in vitro reconstitution system -type I (see Methods), with purified EGFP-synapsin-1 (Syn 1), α-synuclein-AF647 (αSyn) and Cy3-labelled *Scg3* or *Prnp* RNAs. Bottom: Representative micrographs of type I in vitro reconstituted Syn 1/αSyn condensates with *Scg3* or *Prnp* RNA at 25 min of incubation upon protein-protein condensate formation corresponding to 15 min post mRNA addition (t2). **(B)** Quantifications of RNA fluorescence intensities (*Scg3* and *Prnp*) in in vitro reconstituted protein-protein droplets after RNA addition (t1) and at the time point t2 (see above). A two-tailed Mann-Whitney U test comparing RNA intensities at t1 to those at t2 indicates a P-value of <0.00001 (****) for both RNAs tested. **(C)** Quantifications of Syn 1 and αSyn fluorescence intensities expressed as a ratio (Syn 1/αSyn) in in vitro reconstituted protein-protein droplets without or in the presence of *Scg3* or *Prnp* RNAs 25 min after reaction set up (t2). A two-tailed Mann-Whitney U test comparing the ratio of Syn1/αSyn intensities before and after the addition of RNA indicates a P-value of <0.00001 (****) for both RNAs tested. **(D) ,** Top: schematic representation of three-component in vitro reconstitution system type II (see Methods), with purified EGFP-Syn 1, αSyn-AF647 and Cy3-labelled *Scg3* or *Prnp* RNAs. Bottom: Representative micrographs of type II in vitro reconstituted Syn 1/αSyn condensates with *Scg3* or *Prnp* RNA 25 min of incubation addition of all three components in the reaction mix. **(E)** Quantifications of αSyn fluorescence intensities in in vitro reconstituted Syn 1/*Scg3*/αSyn and Syn 1/*Prnp*/αSyn condensates at two different time points (t1=5 and t2=25 min upon type II reaction set-up). Paired two-tailed t-test comparing the intensities of αSyn within condensates at t1 and t2 result in a P-value of 0.1652 and 0.1633 for conditions containing *Scg3* and *Prnp*, respectively. All the box plots (b, c, e) encompass the 25^th^ and 75^th^ quartiles with the median marked by central line. Whiskers extend to 1.5 times the interquartile range (IQR). Circles indicate outliers. All the data were collected from three independent in vitro reconstitutions.

The rapid and continued recruitment of RNA to preformed synapsin/α-synuclein condensates (**Fig 3B**) prompted us to investigate how readily α-synuclein could be recruited to preformed synapsin-RNA condensates. After preforming synapsin-RNA condensates and supplementing reactions with α-synuclein (**Fig. 3D**), we measured α-synuclein recruitment into synapsin-1/RNA condensates at two different time points (**Fig. 3D, E**). Despite seeing an initial, moderate recruitment of α-synuclein to synapsin-1/RNA condensates, no additional enrichment was observed in the course of incubation time. This is in contrast to synapsin-1 and RNA, which continued recruitment to preformed synapsin-1/RNA condensates over the two time points measured (**S7E, F**). Interestingly, the difference in structural heterogeneity between tested synaptic RNAs was insufficient to drive significant differences in the amount of synapsin recruited to condensates (**S8**), as it was observed with the *CLN3* RNAs (**Fig. 2B**). It is important to note that in addition to differences in the magnitude of structural heterogeneity between the two RNA systems (*CLN3* ED range = 114 – 627; Prnp/Scg3 ED range = 435 – 635), there are notable length differences (*CLN3* = 1588 nt; Prnp/Scg3 = 2193 nt) that could contribute to different outcome of these reactions. Furthermore, these reactions were only carried out in a 1:1 charge ratio, which results in the same ratio of synapsin to nucleotide content between the two systems, but results in different numbers of RNA molecules per reaction, with *Scg3* and *Prnp* reactions containing fewer RNAs. Nonetheless, altogether, these results suggest that RNA can be a potent factor in modulating the condensation of synapsin-1 in a minimal system.

### 2.4 The acute RNA disruption causes a dispersion of synaptic vesicles in living neurons

After observing synapsin-1/RNA condensation in vitro, we wanted to know whether the global degradation of RNA could lead to the depletion of synapsin-1 condensates at the presynapse of primary hippocampal neurons. To promote the uptake of RNase A into neurites we employed an approach based on photochemical internalization (PCI) system ^[52]^ ^[53]^, modified in a way to prevent damage to recycling organelles (**Fig. 4A, S9)**. After PCI-mediated membrane permeabilization, 150 µg/ml of RNase A, or BSA as a control, was supplemented to WT rat neuronal cultures co-expressing EGFP-synaptophysin (Syp) and mScarlet-synapsin-1. We then measured the fluorescence intensity and distribution of synaptophysin and synapsin-1 at synaptic boutons (**Fig. 4B, S10**). We observed a significant decrease of fluorescent signal of both proteins, as well as their dispersion from the synaptic boutons (**Fig. 4C, D**). As opposed to RNase A treatment, no effect was observed when neurons were incubated with BSA (**Fig. 4E-G, S11**), indicating the importance of intact RNA molecules for synaptophysin and synapsin-1 enrichment at the presynapse. Together, these results in live neurons clearly demonstrate that synapsin-1 enrichment at the presynapse is dependent on the presence of undegraded RNA pools. Our results so far suggest RNA as a potential scaffold, for the promotion and or stabilization of synapsin-1 condensates both in vitro and primary neurons.

**Figure 4.**
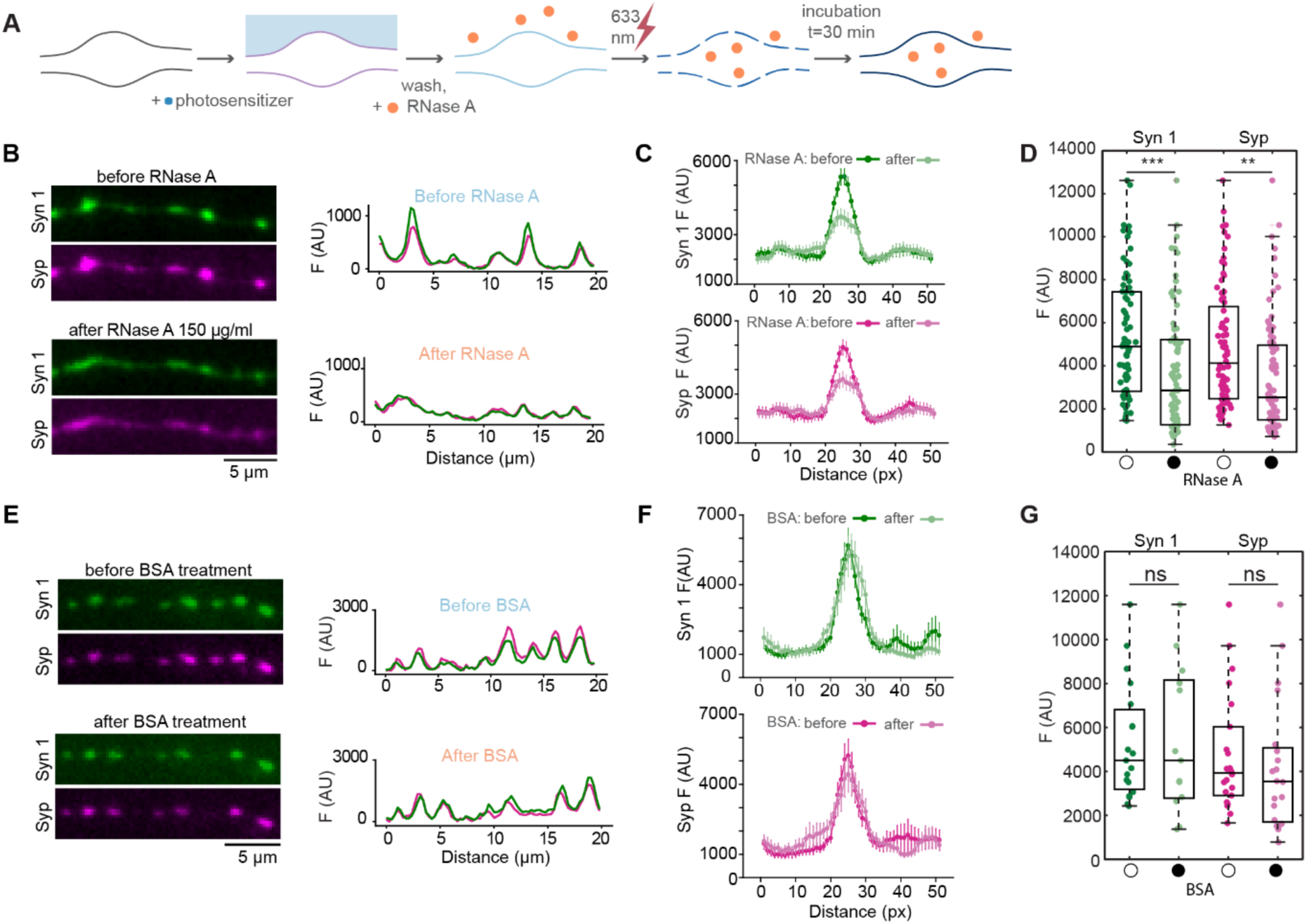
Acute RNA disruption leads to partial dispersion of synaptic vesicle cluster. **(A)** Scheme representing a modified photochemical internalization (PCI) method applied for RNase A treatment of rat primary hippocampal neuron culture. **(B)** Representative micrographs of axon before and after RNase A treatment showing mScarlet-synapsin-1 (Syn 1) and EGFP-synaptophysin (Syp) signal distribution at the synaptic boutons accompanied with the corresponding Syn 1 (green) and Syp (magenta) fluorescence intensity line profiles of representative axon segment. **(C)** Averaged intensity line scans of individual boutons representing Syn 1 (above) and Syp (below) fluorescence intensity signal before and after RNase A treatment. Error bars represent standard error of the mean (± SEM). Data was collected from 3 independent primary cultures. **(D)** Quantifications of Syn 1 and Syp peak fluorescence signal at the synaptic boutons before and after RNase A treatment. The box plot of quantified peak fluorescence signal encompasses 25th and 75th quartiles with median marked by central line. Minimum and maximum are denoted by whiskers. Individual circles represent peak fluorescence signal of a single bouton for each condition. ** indicates P ≤ 0.01 and *** indicates P ≤ 0.001 by paired t-test. Data was collected from three independent biological replicates. **(E)** Representative micrographs of axon before and after BSA treatment showing Syn 1 and Syp signal distribution at the synaptic boutons in the presence of the photosensitizer used for PCI. **(F)** Averaged intensity line scans of individual boutons representing Syn 1 (left) and Syp (right) fluorescence intensity signal before and after BSA treatment. Error bars represent standard error of the mean (± SEM). Data are from three independent biological replicates. **(G)** Quantifications of Syn 1 and Syp peak fluorescence signal at the synaptic boutons before and after BSA treatment. The box plot of quantified peak fluorescence signal encompasses 25^th^ and 75^th^ quartiles with median marked by central line. Minimum and maximum are denoted by whiskers. Individual circles represent peak fluorescence signal of a single bouton for each condition. ns indicates non-significant by paired Wilcoxon test. Data were collected from three independent biological replicates.

### 2.5 The interface of synapsin-1 condensates harbors translational machinery

To visualize mRNA at the presynapse, we used fluorescent in situ hybridization (FISH) directed towards the poly(A) tail of mRNA at DIV14. We observed an obvious enrichment of mRNA at vGLUT1 positive foci, a presynaptic marker (**Fig. 5A, B and S12A**), suggesting a local enrichment of mRNA at the presynapse. In addition, employing single molecule (sm)FISH in primary neurons we observed that highly abundant mRNA, *Atp5b*, coding for β subunit of ATP synthase ^[54]^ ^[55]^ was also colocalizing with our presynaptic marker, vGLUT1 (**Fig. 5C, D and S12B-D**).

**Figure 5.**
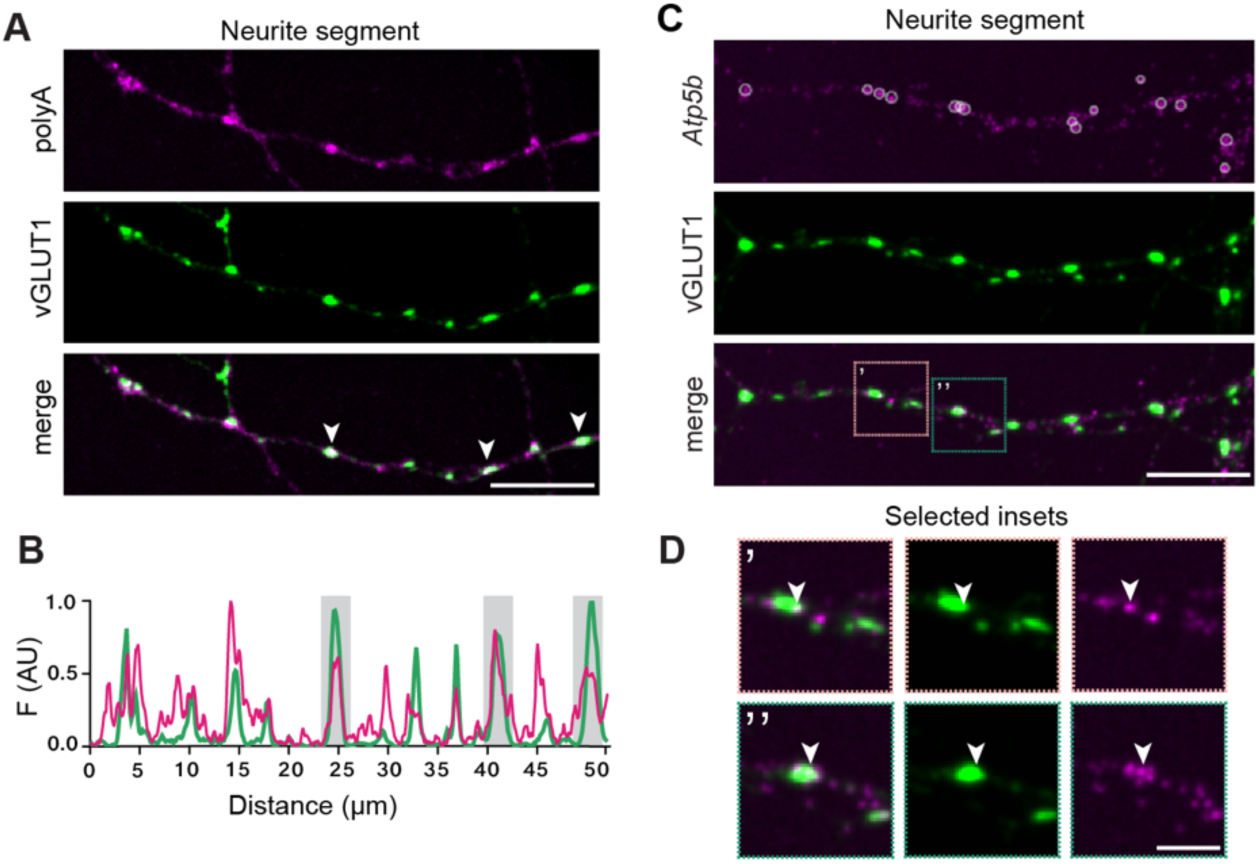
Synaptic vesicle clusters contain coding RNA. **(A)** Representative images of neurite segment showing poly(A) FISH signal (magenta) and immunostaining against presynaptic marker vGLUT1 (green) in mouse primary hippocampal neurons. Arrows in merge indicate overlap between poly(A) and vGLUT1 in the presynaptic compartment. **(B)** Line profiles represent quantified normalized fluorescence intensity signal (F) of poly(A) and vGLUT1 along 50 µm long neurite segment presented in images above. Scale bar, 10 µm. **(C)** Representative images of neurite segment showing neurite segment with detected *Atp5b* smFISH (magenta) with white circles indicating segmented *Atp5b* foci using RS-FISH analysis software and vGLUT1 signal (green). In merge panels light orange (’) and green (’’) insets highlight presynaptic region shown in panel **(D)**: Scale bar, 10 µm. Arrows indicate *Atp5b* signal co-localizing with vGLUT1. For insets scale bar, 3 µm.

Given that RNA can be recruited to synapsin-1 condensates, we hypothesized that biomolecular condensation of synapsin-1 may function in modulating translation capacity of the recruited transcripts in addition to organizing SVs. This would provide an additional mechanism by which neurons regulate local protein synthesis, a critical feature of these large and highly polarized cells. To test if there is a role of synapsin-1 in regulating mRNA translation, we first wanted to test whether synapsin-1 condensates could associate with the components of translational machinery in addition to their association with RNA molecules.

To this end, we ectopically generated SV-like condensates in HEK293 cells. Our immunocytochemistry analysis in HEK293 cells showed that eukaryotic initiation factor 4E (EIF4E), one of the components of the translational machinery, colocalizes with or at the interface of mCherry-synapsin-1 condensates (**Fig. 6A-C, S12E**). Moreover, ultrastructural analyses using correlative light-electron microscopy indicates ribosomal structures at the interface of vesicular condensates in cells (**Fig. 6, S13**).

**Figure 6.**
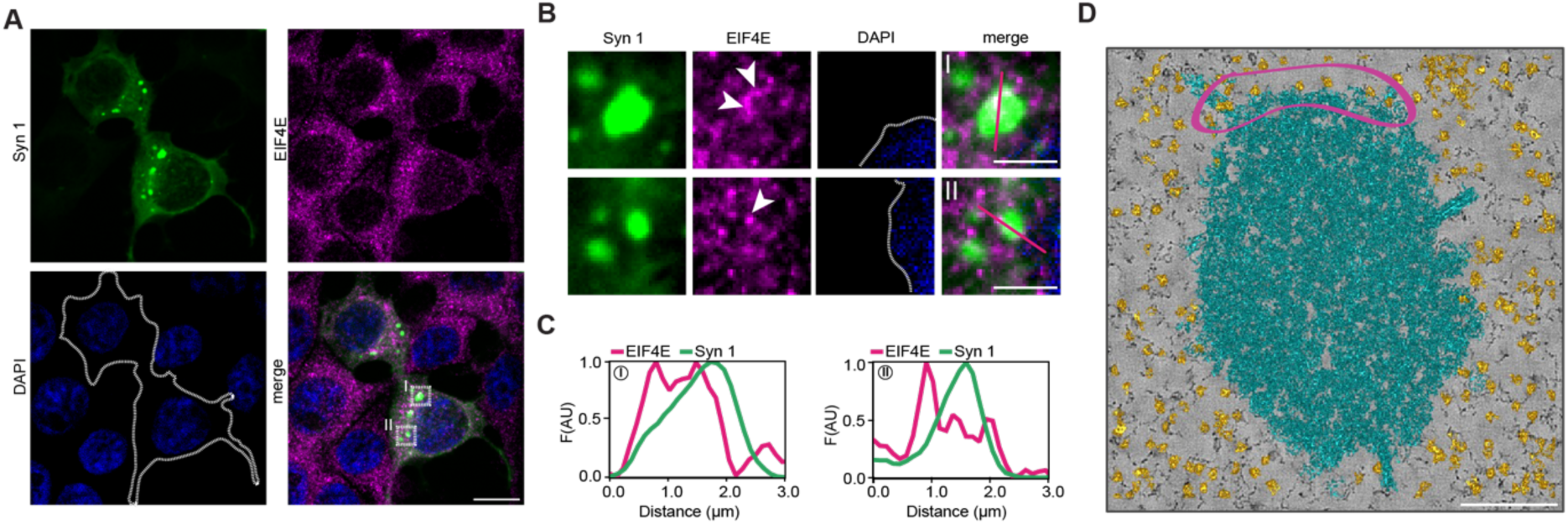
Synapsin-1 condensates harbors the translational machinery components. **(A)** HEK293 cells expressing synapsin-1 condensates (mCherry -synapsin-1; Syn 1 in green) fixed and immunostained for EIF4E translational regulator (magenta), DAPI: nucleus. Scale bar, 10 μm. **(B)** Magnified regions of condensates from panel A (white dashed quadrats). Arrows indicate few EIF4E foci. In merge note the accumulation of EIF4E at the interface and within condensates. Gaussian filter (0.4) was applied to magnified insets for better visualization. Dark pink lines in merge (I and II) indicate trajectory of respective fluorescence intensity (F) line profiles in panel C. Scale bars, 3 µm. **(C)** Line profiles across exemplary condensates shown in **B**. **(D)** Exemplary tomogram of a correlative light-EM image of cells ectopically expressing Syn 1/vesicular condensates. Region highlighted in magenta indicates the accumulation of ribosomes (orange) at the interface of vesicular condensates (cyan). Scale bar, 200 nm.

### 2.6 Synapsin-1 condensates are permissive for mRNA translation in vitro

To assess if the association of mRNAs with synapsin influences their translation, we first generated a luciferase reporter for one of the synaptic mRNAs analyzed above, *Atp5b*, which is highly abundant at the presynapse. In our reconstitution assays, *Atp5b* readily co-condenses with synapsin-1 when mixed in a 1:1 polyelectrolyte charge ratio (**Fig. 7A**). To measure translation in the presence of synapsin-1 condensates, we preformed synapsin-1/RNA condensates with *Atp5b* mRNA containing a 3’ fusion to NanoLuciferase (Nluc). After the formation of condensates, rabbit reticulocyte extract (RRE) was added to the reaction and left to incubate at 30℃. After one hour, vivazine was added to the reactions and luminescence generated by the successful translation of *Atp5b-Nluc* was read out on the luminometer. Strikingly, mRNAs in reactions containing pre-formed synapsin-1/RNA condensates showed enhanced translation when compared to RNA only reactions, and translation could be further increased by the addition of α-synuclein to reactions (**Fig. 7B, C**). Importantly, this increase in translation is not a byproduct of simply crowding or the mere presence of synapsin-1, as the addition to reactions of 4.5 µM BSA or the inclusion of synapsin-1 IDR mutant that is unable to phase separate ^[43]^ did not enhance translation above RNA alone. These data strongly indicate that the presence of synapsin-1 condensates in the reaction mixture augments the translational capacity of mRNAs.

**Figure 7.**
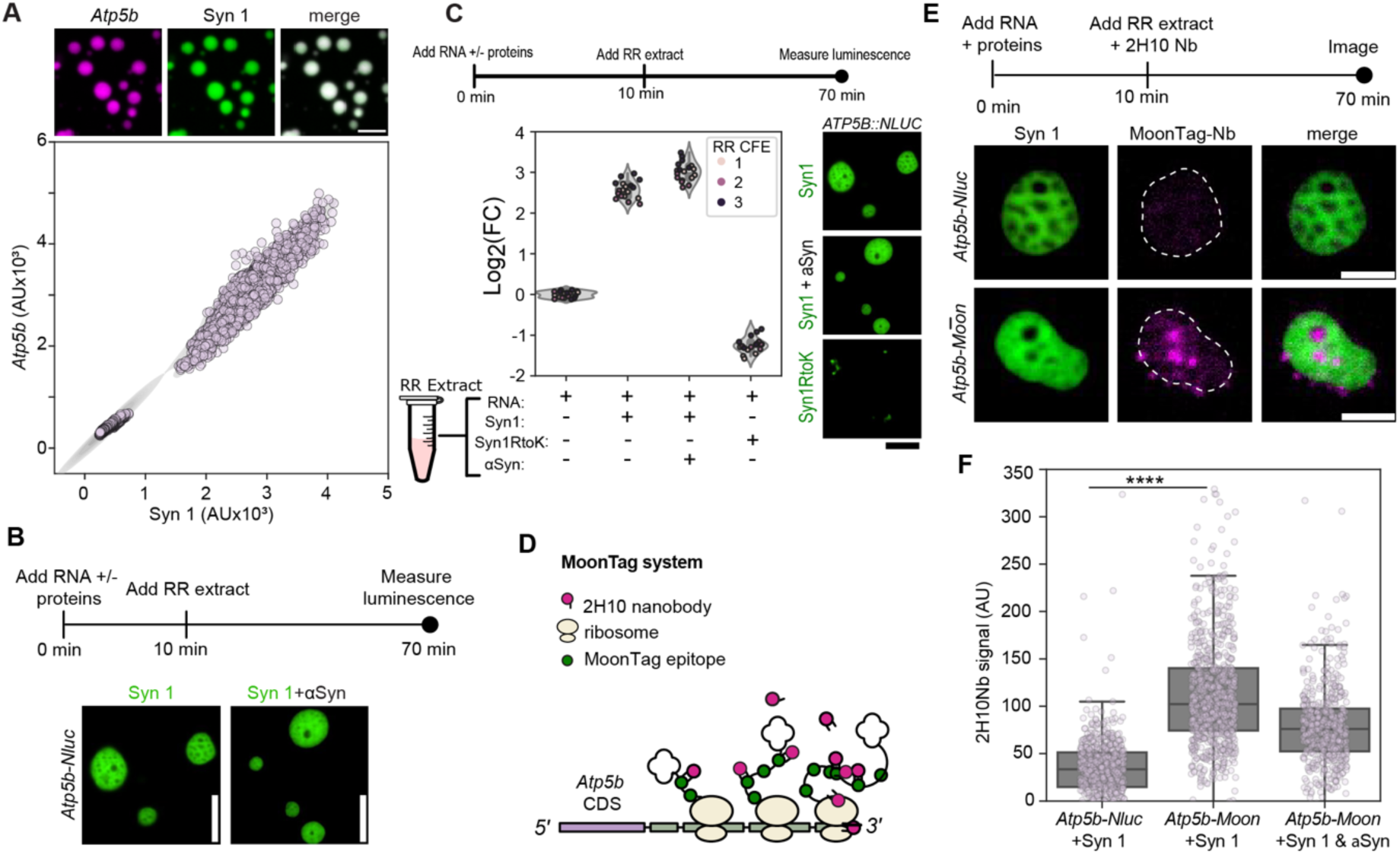
Synapsin-1 condensates promotes mRNA translation. **(A)** Above: images showing *Atp5b* synaptic mRNA (magenta) enriching synapsin-1 condensates (Syn 1, green) in in vitro system under physiological conditions and in the absence of any molecular crowders. Scale bar, 5 µm. Below: Scatter plot depicts quantified *Atp5b* RNA intensities as a function of Syn 1 intensities. Each circle represents individual condensate analyzed overlaying Kernel Density Estimate (KDE). Data were collected from three independent in vitro reconstitutions of in vitro transcribed Cy5 labelled *Atp5b* and purified EGFP-tagged synapsin-1. **(B)** Top: scheme represents simplified overview of the in vitro luciferase translation assay using *Atp5b*-nano-luciferase (*Atp5b-Nluc*) as an RNA reporter. Below: representative images of Syn 1 and Syn 1/α-Syn condensates in rabbit reticulocyte extract (RRE). Scale bar, 5 µm. **(C)** Top: scheme represents simplified overview of the in vitro luciferase translation assay using *Atp5b*-nano-luciferase (*APT5B:NLUC*) as a RNA reporter. Bottom: the violin plot showing the quantification of the *APT5B:NLUC* luminescence signal under different reaction conditions (only *APT5B:NLUC, APT5B:NLUC* + BSA*, APT5B:NLUC* + Syn 1, *APT5B:NLUC* + Syn 1 + α-Syn, and *APT5B:NLUC* + Syn 1RtoK). Within each violin plot, the box depicts the 25th through the 75th quartiles, with whiskers extending to min and max data point. Each individual circle represents a technical replicate. Data from from 3 independent extracts. Right: representative images of Syn 1, Syn 1/αSyn, and Syn1RtoK condensates after the addition of rabbit reticulocyte (RR) extract. **(D)** Scheme of MoonTag system developed to quantitatively detect translation events live *in cellulo* and in our work optimized for in vitro system of Syn 1 condensates in RR extract. **(E)** Top: Scheme of the MoonTag translation pipeline in RR extract using *Atp5b-*MoonTag (*Atp5b-Moon*) RNA reporter system. Bottom: Representative images of Syn 1 (green) and MoonTag(-Nb) nanobody signal (magenta) in condensates enriched with Syn 1 and *Atp5b-Nluc* used as negative control (upper image panel) and *Atp5b-MoonTag* reporter RNA (lower image panel) 60 min after the addition of RRE to pre-formed Synapsin-1/RNA condensates. Scale bar, 2 µm. **(F)** Bx plot showing anti-MoonTag nanobody (2H10Nb) fluorescence values for the different in vitro conditions (only *Atp5b-Nluc+*Synapsin-1 [negative control], *Atp5b-Moon*+ Syn 1*, Atp5b-Moon*+ Syn 1+ α-Syn) 60 min after the addition of RRE. For each box plot, the horizontal line in the box plot denotes the median value of data distribution between 25^th^ and 75^th^ percentiles. Whiskers extend to 1.5× the IQR. Each magenta circle represents individual condensate analyzed. Data were collected from 3 independent RR extracts. Paired, two-tailed t-tests were carried out comparing mCherry-2H10Nb signal values between reactions containing *Atp5b-Nluc* and two conditions: (1) *Atp5b-Moon*+Syn 1 and (2) *Atp5b-Moon*+Syn 1+ αSyn; P-values <0.0001 (****), for both conditions respectively. First comparison is highlighted on the graph (****). The same statistical test is performed to compare *Atp5b-Moon*+Syn 1 and *Atp5b-Moon*+Syn 1+aSyn conditions (P-value = 0.001166).

We next asked if this promotion of translation was happening within or at the synapsin condensate interface. To visualize translation events, we turned to a fluorescence-based approach called MoonTag (**Fig. 7D**) ^[56]^. This system relies on the strong association of a fluorescently labeled nanobody (2H10Nb) with nascently translated MoonTag (gp41) peptides. By fusing a sequence coding for 24 MoonTag peptides to *Atp5b* mRNA coding sequence, we can visualize the translation of mRNAs through the accumulation of fluorescently-labeled nanobodies. As above, we pre-formed synapsin-1/RNA condensates with *Atp5b-MoonTag* mRNA and then supplemented reactions with RRE and mCherry-labeled nanobodies. The MoonTag did not change the basic conditions at which synapsin-1 to phase separates (*Atp5b-MoonTag = 3052 nt; ED = 1025)*. To our surprise, we saw large nanobody foci contained within pores within synapsin-1 condensates as well as foci on the surface, complemented with detected fluorescence signal of Atp5b-Moon nascent peptide (visualized with 2H10Nb nanobody) in synapsin-1/*Atp5b-Moon* condensates as compared to synapsin-1/*Atpb5b-Nluc* negative control (**Fig. 7E F**). Notably, these same pores could be seen in condensates formed with *Atp5b-Nluc* when added to RRE, however the recruitment of nanobody was specific to condensates having RNAs with the MoonTag sequence, indicating that synapsin-1 condensates are not concentrating the nanobody itself (**Fig. 7D**).

Additionally, large dextrans readily pass through these condensates, suggesting translation machinery may also concentrate within these condensates (**S14**). Interestingly, the average nanobody signal associated with synapsin condensates in MoonTag reactions containing α-synuclein was lower than those containing synapsin alone (**Fig. 7F**). This is in contrast to the luciferase reaction containing both synapsin and α-synuclein, which showed an increase in mRNA translation over synapsin-only conditions (**Fig. 7B**). These results would seem to suggest either that (i) MoonTag signal provides a spatial marker for where translation is occurring but the average signal per condensate is not an informative metric with regards to translation efficiency or (ii) the addition of α-synuclein leads to the differences in residency of already translated, nanobody-bound MoonTag peptides, leading to a loss of nanobody signal from synapsin condensates. Regardless of the interpretation, the increase in translation of mRNAs in reactions containing synapsin-RNA condensates (**Fig. 7C and F**), and the observation of translational microdomains within synapsin condensates (**Fig 7E**), suggests that synapsin-1 condensates may act as translational crucibles for organizing protein synthesis.

## 3. Discussion

In this study, we discover that synapsin-1, a highly abundant protein in the presynapse, readily forms condensates with RNA well below the physiological concentrations of this protein. In vitro reconstitutions indicate increased synapsin recruitment in condensates formed with RNA with lower conformational heterogeneity. Consistent with some role for RNA structure, there is a moderate bias in RNAs that associate with synapsin-1 condensates in cells to be short and have lower relative conformational heterogeneity. Global disruption of RNA in cultured neurons is sufficient to disperse synapsin-1 and SVs at the presynapse. In addition to RNA playing a role in condensate formation/maintenance, we provide evidence that RNA translation may be promoted by synapsin-1 condensates.

### RNA structure heterogeneity can influence synapsin-1 condensates

Synapsin-1 lacks any classical RNA binding domain; thus, it is likely that binding and condensation of synapsin and RNA is driven primarily through electrostatic interactions between the highly positively charged synapsin-1 and the negative charge of RNA ^[57]^. Indeed, the IDR of synapsin-1, a necessary and sufficient region for its phase separation ^[42]^, was also sufficient to form condensates with RNAs, although of smaller size (**S4**). However, in addition, our data show that differences in RNA conformational heterogeneity can alter recruitment of client proteins such as α-synuclein and impacts the concentration of both RNA and proteins in synapsin-RNA condensates (**Fig. 2 and 3**). We find that condensates formed by less conformationally heterogeneous RNAs promote greater partitioning of synapsin-1 (**Fig. 2**). These results indicate that in addition to length and nucleotide composition, RNA structural heterogeneity may act as a knob by which neurons might tune the formation and dissolution of synaptic vesicle-containing condensates in the small volume of synaptic bouton. RNA modifications, editing and helicases could all shift the ED of a given sequence and impact the concentrations/properties of synapsin-1 condensates locally in the neuron. Furthermore, RNA itself may contribute to the composition of SV condensates in the complex milieu of the presynapse, where multiple competing interactions among proteins can occur.

### α-Synuclein competes with RNA for enrichment into synapsin/SV condensates

α-Synuclein, a prominent SV cluster component, is recruited into synapsin/SV condensates ^[49]^. Recently, α-synuclein was demonstrated to interact with few components of P-bodies, another prominent example of RNA granules. Moreover, many of these proteins are part of the decapping machinery, suggesting a role for α-synuclein role in modulating the stability of mRNA stored therein ^[58]^. Together, these two studies point to a direct relation between the SV cluster and mRNA processing, presumably modulated by α-synuclein. While neither α-synuclein alone nor SVs alone were sufficient to form condensates our data show that the partitioning of α-synuclein into synapsin condensates is sensitive to the presence of RNA, strongly suggesting RNAs could be both a substrate as well as competitor for interactions with synapsin-1 (**Fig. 2 and Fig. 3**). These intricate associations between proteins and RNAs could be important in the organization of the presynapse, with dysregulation of protein-RNA interactions potentially contributing to the onset or emergence of neurodegeneration ^[59]^ ^[60]^.

In addition to *CLN3* orthogonal system, we probed the effect of two structurally different synaptic mRNAs (*Scg3* and *Prnp*) ^[51]^ identified at the synapse on in vitro condensation of synapsin-1 and α-synuclein. Even though we observed a clear competition between mRNAs and α-synuclein (as with orthogonal mRNAs in **Fig. 2**), no difference between *Scg3* and *Prnp* synapsin-1/α-synuclein condensates were observed under our experimental conditions. This can be the consequence of these two transcripts having more similar ED values than the *CLN3* sequences, suggesting that more extreme structural differences or specific RNA sequence motifs are required to drive differential condensation into synapsin condensates. Nonetheless, we do not exclude the possibility that a different RNA/protein initial concentration, would reveal differences in dense phase recruitment as a function of ED and/or RNA secondary structure.

### Synapsin-RNA condensates – potential hubs for the local translation

Spatial control of translation was first implicated in neurons after the observation of preferential localization of polyribosomes at the base of dendritic spines^[61]^ reinforcing the notion that local protein synthesis occurred in dendrites but not in axons or presynaptic sites. This view evolved with the data demonstrating the local translation at the presynapse^[62],[63]^. Our studies demonstrate a potential role for synapsin-1 in regulating this process at the presynapse through the formation of biomolecular condensates. We show that, in contrast to some ribonucleoprotein condensates^[64],[65],[66]^, the presence of synapsin condensates enhances translation of RNAs by several orders of magnitude (**Fig. 7**). Though the mechanistic basis of this enhancement remains unknown, it is possible that the positive charge of synapsin-1, thought to be the major driving force in synapsin-RNA interactions, may also aid in capturing and concentrating negatively charged ribosomal components or other translational machinery. Such pronounced translation activity within synapsin condensate microdomains and at the interface of synapsin condensates^[67]^ may be the result of interfacial ion potential that can generate permissive chemical environments^[68]^ and necessary protein conformations^[69]^ to facilitate effective translation.

Our spatial analysis of translation in synapsin condensates support this hypothesis, as we observe sub-domains within synapsin-1 condensates that show signal for active translation (**Fig. 7**). Our dextran-based assessment (**S14**) of the mesh pore size within condensates suggests that pores are sufficiently large for translated proteins to diffuse out of the condensate, indicating that these fluorescently labeled MoonTag nanobodies depict the ‘hot spots’ of active translation within the condensates. These observations build further evidence of condensate functions beyond translational repression and RNA storage. For neurons, this translational enhancement may be particularly relevant in situations where concentrations of RNAs and ribosomal machinery are low ^[15]^. In these scenarios, biomolecular condensation may be indispensable in ensuring RNAs are translated to the proper extent and at the right time, despite a low abundance of translational machinery.

## 4. Conclusions

In summary, our study provides a direct insight into the architectural role of RNA in organizing the synapsin-1 condensates in a minimal system and modulating the synapsin-SV condensates distribution at the synaptic boutons. We also demonstrate that translation of mRNA can be enhanced in synapsin-1 condensates, indicating a mechanism for coupling of SV release and local translation, a mechanism recently proposed to underlie sustain synaptic activity^[63]^. Thus, this study demonstrates the importance of RNA in organizing the presynaptic condensates, where competing interactions among nucleic acids and proteins, such as α-synuclein, may play a critical role in regulating the spatial and temporal protein localization vital for synaptic physiology and disrupted in neurodevelopmental and neurodegenerative diseases.

## 5. Experimental Section

### Cloning

Synapsin-1 full-length (FL) (*Homo sapiens*) was cloned into the pmCherry-C1 vector using BglII-SacI restriction sites (Genscript). pmCherry-C1 were modified through NheI-XhoI sites to encode the 6×His-tag ^[42]^. Synapsin-1 FL (*Rattus norvegicus*) or IDR (*Homo sapiens*) were cloned in pEGFP-C1 vector using BglII-SacI restriction sites (Genscript). pEGFP-C1 was modified through NheI-XhoI sites to encode the 6×His-tag^[41]^. The untagged α-synuclein was cloned from Addgene plasmid #40822 into pBFP N1 backbone using BglII and SalI restriction sites. All the vectors contain CMV promoter and SV40 terminator. Untagged Synaptophysin was PCR amplified from synaptophysin-EGFP (kind gift of Pietro De Camilli) in order to delete EGFP tag and using HindIII and NotI restriction sites reinserted in the same backbone now free of EGFP-tag. α-synuclein (*Homo sapiens*) for protein expression was cloned into pET28 expression vector using restriction enzyme sites NcoI-NotI. All DNA sequences used for cloning are provided in Supplementary **Table S1**.

The modified *Ashbya gossypii CLN3* sequences used in in vitro reconstitution assays were purchased from Genewiz as gene fragments and cloned into the pJet vector (ThermoFisher Scientific K1231) using blunt end cloning. Primers for amplifying *CLN3* are listed in Supplementary **Table S2.** Directionality and sequence were confirmed by Sanger sequencing (Genewiz).

The *Mus musculus Atp5b*, *Prnp*, and *Scg3* sequences used for translation and in vitro reconstitution assays were ordered from Genewiz as gene fragments and cloned into the pJet vector (ThermoFisher Scientific K1231) using blunt end cloning. Primers for amplifying *Atp5b*, *Prnp*, and *Scg3* are listed in Supplementary **Table S2**. Directionality and sequence were confirmed by Sanger sequencing (Genewiz).

Full-length Nanoluciferase (Nluc; Addgene, plasmid #60140) was amplified using primers containing overlap with the 3’ end of *Atp5b*. The following Nluc fragments and *Atp5b* DNA were included in a PCR reaction, allowing the amplification of the *Atp5b*-Nluc. *Atp5b*-Nluc was then cloned into the pJet vector (Thermofisher Scientific K1231) using blunt end cloning. Primers for the amplification of *Atp5b*-Nluc are listed in Supplementary **Table S2**.

A custom MoonTag sequence was cloned into the pJet vector using blunt end cloning as described in ^[70]^. To insert *Atp5b* into the custom MoonTag cassette, *Atp5b* was amplified using primers containing a 5’ DraIII and 3’BsiWI restriction sites and cloned into the MoonTag cassette using sticky-end cloning. Primers use to amplify *Atp5b* for insertion into the MoonTag cassette are listed in Supplementary **Table S2**.

### Protein purification and labelling

#### EGFP-synapsin-1 purification

Synapsin-1 purification was performed using previously published protocols ^[42]^ ^[43]^ ^[71]^. In brief, 6×His-EGFP-rr-synapsin-1 full-length (EGFP-synapsin-1 FL) and 6×His-EGFP(A206K)-hs-synapsin-1-IDR (421-705) (EGFP-synapsin-1 IDR) were expressed in Expi293F™ cells (Thermo Fisher Scientific) for three days prior to induction. Cell lysis was performed by three freeze-thaw cycles in liquid nitrogen and at 37°C, respectively in a buffer containing 25 mM Tris-HCl (pH 7.4), 300 mM NaCl, 25 mM imidazole, 0.5 mM TCEP (buffer A) supplemented with EDTA-free protease inhibitor (Roche, cOmplete Mini, 11836170001), 10 µg/mL DNase I (AppliChem A3378) and 1 mM MgCl_2_. Next, the lysate was pelleted by centrifugation for 1 hour at 20,000 × g followed by two-step purification. First affinity purification of soluble supernatant was performed on a Ni-NTA column (HisTrap™HP, Cytiva, ÄKTA pure 25M). Washing steps were performed with buffer A containing 40 mM imidazole, followed by elution in buffer A supplemented with 400 mM imidazole. Post elution fractions were concentrated (30K MWCO protein concentrator, Pierce) and submitted to size-exclusion chromatography (Superdex™ 200 Increase 10/300, GE Healthcare, ÄKTA pure 25M). Size-exclusion chromatography was performed in 25 mM Tris-HCl (pH 7.4), 150 mM NaCl, 0.5 mM TCEP. All steps of protein purification were performed at +4°C. Prior to use purified proteins were snap-frozen in liquid nitrogen and stored at -80°C.

#### α-Synuclein purification

Purification of untagged human α-synuclein was performed as previously described ^[49]^ ^[72]^. Untagged hsSNCA (A140C) was expressed in BL21-(DE3) chemically competent *E. coli* cells from a pET28 expression plasmid (Novagen). Cell lysis was performed in two rounds using high-pressure French press (10 000-15 000 psi, The French®, SLM-AMINCO) in cold 1× PBS supplemented with EDTA-free protease inhibitor (Roche, cOmplete Mini, 11836170001). Next, the lysate was supplemented with lysozyme and transferred to Beckman tube (#357000) for further clarification. To clarify the lysate 4 rounds of high-speed centrifugation (JA 25-50 rotor) were applied, each 30 min at 50000 × g at +4°C. After the round 1 and 2 cleared supernatants were collected and supplemented first with DTT (1 mM) and streptomycin-sulphate (10 mg/mL) followed by addition of ammonium-sulphate (0.361 g/ml) after round 2. Following third round of centrifugation pellet was resuspended in cold 1× PBS supplemented with 1 mM DTT and boiled for 20 min, after which it was cooled down and subjected to the final round of high-speed centrifugation. Obtained supernatant was dialyzed overnight against 15 mM Tris-base solution, cleared through 0.2 µm filter (7 bar max, puradisc 30/0.2 CA S, Whatman, Cytiva) and as such used for 2-step purification process. First, ion-exchange chromatography of dialyzed supernatant was performed on HiPrep^TM^ Q FF 16/10 column (Cytiva, ÄKTA pure 25M) using anion-exchange buffer gradient (0-500 mM NaCl in 25 mM Tris-base, pH 7.4, 1 mM DTT). Elution was performed in 250 mM NaCl in 25 mM Tris-base, pH 7.4, 1 mM DTT and fractions were concentrated with 3K MWCO protein concentrator (Millipore), after which size-exclusion chromatography (Superdex™ 75, Increase 10/300, GE Healthcare, ÄKTA pure 25M) was applied in 25 mM Tris-HCl (pH 7.4), 150 mM NaCl, 0.5 mM TCEP. Prior to use, purified proteins were snap-frozen in liquid nitrogen and stored at -80°C.

#### α-Synuclein labelling

Purified α-synuclein was labelled with thiol-reactive probe Alexa Fluor™ 647 C_2_ Maleimide (Invitrogen, A20347) using manufacturer protocol. In brief, 130 µM of the starting protein solution (150 mM NaCl, 25 mM Tris buffer pH 7.4) was prepared at room temperature. Next, reduction of disulfide bonds was obtained adding TCEP to a final concentration of 1.3 mM. To label the protein 3× in excess of reactive dye to a starting protein concentration was added dropwise to the protein solution and labelling reaction was performed in dark at room temperature for 2 h. Upon reaction completion the sample was subjected to gel filtration on PD-10 Sephadex^TM^ G-25 M column (Cytiva) to separate labelled protein. The concentration of labelled protein was calculated using following formula:

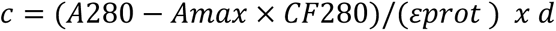

where A280 = absorbance of the protein at 280 nm, Amax = absorbance of the dye at the absorption maximum, εprot = extinction coefficient of the protein, CF280 = correction factor for the dye absorption, d = path length.

#### 2H10-mCherry purification

2H10-mCherry was expressed and purified from in BL21 (DE3) *E. coli* cells as described in ^[70]^. Briefly, cells were collected by centrifugation, and bacteria were lysed using lysis buffer (20 mM HEPES pH 7.4, 100 mM KCl, 1x EDTA free Rosche protease inhibitor tablet) and probe sonication (Qsonica Ultrasonic Sonicator; CL-334)). Protein was purified from the crude bacterial extract by incubating with cobalt resin (HisPur^TM^, Thermo Fisher Scientific), followed by extensive washing with lysis buffer (at least 10× column volumes). Protein was retrieved from the cobalt resin using elution buffer (20 mM HEPES pH 7.4, 100 mM KCl, 200 mM Imidazole. Flow through was collected and dialyzed overnight in storage buffer (lysis buffer excluding the Rosche protease inhibitor tablet) using a 10 000 MWCO cassette (Slide-A-Lyzer® Dialysis Cassette G2). After dialysis, the concentration of 2H10-mCherry was measured by Bradford assay. Purified protein was stored at -80℃ prior to use. Prior to the addition of 2H10-mCherry to MoonTag reactions, the N-terminal 6×His tag was cleaved off by TEV-protease by incubating 2H10-mCherry with TEV-protease (17:1) at room temperature for 30 min.

### RNA extraction, in vitro transcription and labelling

#### RNA extraction of total neuronal RNA

For in vitro reconstitution assays total neuronal RNA was purified from mouse primary hippocampal neuron culture using modified TRIzol extraction protocol. In brief, cultured hippocampal neurons were gently washed with cold 1× PBS (Gibco, 10010-015), upon which cells were resuspended in 350 µl TRIzol reagent (ambion, 15596026). To allow phase separation of RNA phase, 70 µl of chloroform (Roth, 4432.1) was added and suspension was vortexed vigorously for 5 s, followed by incubation on ice for 5 min. Next, phase separation was enhanced by centrifugation at 11 000× g for 45 min at +4°C, after which aqueous phase was collected and mixed with 175 µl of 2-propanol (Roth, 9866.1) to proceed with RNA precipitation. 2-propanol RNA precipitation was performed at 11 000× g for 30 min with the help of GlycoBlue coprecipitant (Invitrogen, AM9515), followed by three washes with 75% ethanol. Precipitation and washes were performed at +4°C. After washes RNA was air-dried and eluted in 20-30 µl of UltraPure^TM^ DNase/RNase free water (Invitrogen, 10977-035).

#### In vitro *transcription and labelling*

In vitro transcription of *CLN3, Atp5b, Scg3, Prnp, Atp5b-NLUC,* and *Atp5b-MOONTAG* RNA was carried out according to previously established protocols ^[35]^ ^[73]^ ^[74]^. cDNA template was amplified using PCR (see Supplementary **Table S3**) for all constructs listed except for *Atp5b-MOONTAG*, which required linearization with HindIII digestion. PCR products (or digestion reactions) were purified with a PCR purification kit (Qiagen, 28104). For *CLN3*, *Atp5b*, *Scg3* and *Prnp,* the in vitro transcription was carried out according to manufacturer’s instructions (NEB, E2040S), with the following modifications. Reactions included 1 µg of DNA template and were supplemented with approximately 0.4 µl of Cy3 or Cy5-labelled UTP (Jena Bioscience, NU-803-CY5-S, NU-803-CY3-S). Following incubation at 37℃ for 4 h, the reaction was halted with the addition of 30 µl ddH_2_O. Reactions were than treated with 2 µl of DNase I (Thermo-Fisher, EN0521) and incubated for an additional 20 min at 37℃ to remove DNA template. Reactions were then purified with 2.5 M LiCl precipitation. RNA concentration was determined by 260 nm absorbance and verified for purity and size using a denaturing agarose gel and Millenium RNA ladder (ThermoFisher Scientific, AM7151).

For in vitro transcription of *Atp5b-NLUC and Atp5b-MOONTAG RNA*, the same protocol was used with the following modifications. In vitro transcription was carried out using HiScribe T7 with CleanCap Reagent AG (NEB, E2080S) to produce Cap-1 *mRNA*. These reactions did not include labelled-UTPs, to reduce the impact of bulky adducts on translation. Furthermore, following the first purification and precipitation with 2.5 M LiCl, RNA poly(A) tails were added according to manufacturer specifications (NEB, M0276L). Reactions were then purified a second time with 2.5 M LiCl precipitation. RNA concentration was determined by 260 nm absorbance and verified for purity and size as described above.

To determine RNA labelling efficiency, following absorbance measurements at 260 nm, the same RNA was measured for Cy3 concentration using absorbance at 550, with an extinction coefficient of 150,000 (l/mole-cm), and corrected for absorbance at 260 and 280 using correction factors 0.04, and 0.05 respectively. For Cy5, absorbance was measured at 650, with an extinction Coefficient of 2500,00 (l/mole-cm), and corrected for absorbance at 260 and 280 using correction factors 0.00, and 0.05 respectively. The labelling efficiency was than derived using the formula:

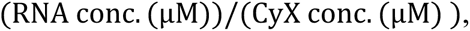

where conc. stands for concentration and X for indicating 3 or 5 Cy conjugate. Labelling efficiency was used to normalize grey values acquired in the Cy3 or Cy5 channel (see Microscopy).

### Mammalian cell culture

HEK293 cells were maintained in DMEM complete medium (DMEM, high glucose, GlutaMAX™ Supplement, Gibco-10566016; 10% Fetal Bovine Serum (FBS), Sigma Aldrich-12106C; 1% Penicillin/Streptomycin, 10 000 U/ml, Gibco-15140122; 1% non-essential amino acids (MEM), Sigma Aldrich- M7145) at 37°C and 5% CO_2_ under sterile conditions. Cells were split every 2 days using 0.05% trypsin (Gibco, 25300-054). For the ectopic reconstitutions of SV-like condensates, approximately 2.4 million cells were seeded on a 10 cm Petri dish and harvested for condensate purification at 90-95 % confluency. Transfection of plasmids for condensate purification or immunocytochemistry (ICC) was performed at 80% of cell confluency using Lipofectamine^TM^ 2000 (Invitrogen^TM^, 11668027) 20 h prior to fixing the cells for ICC.

### Primary neuronal culture

#### Mouse primary culture

Hippocampal neurons were prepared following the previously published procedure ^[42]^. In brief, brains of P0/1 WT mice (C57BL/6) were dissected, and hippocampi were selected for further assessment. After dissection hippocampi were gently washed two times with cold supplemented Hank’s balanced salt solution (HBSS-supp, Gibco, 14170112; supplemented with 10 mM HEPES (Gibco, 14170112), 50 U/ml penicillin/streptomycin (Gibco, 151-40-122), 1 mM sodium pyruvate (Gibco, 11360-039) and 6 mM MgCl_2_ (Roth, KK36.2)]. Next, hippocampi were digested for 20 min at 37℃ with Papain (Worthington, LS003126) to a final concentration of ∼0.47 mgP/ml in HBSS-supp supplemented with 100 µg/ml of DNase 1 (AppliChem, A3378) and 0.6 mg/ml of L-Cysteine Hidrochloride (Sigma Aldrich, C7880). Afterwards, the enzymatic reaction was inactivated and tissue was washed by the plating medium (Neurobasal Medium A (NB-A Gibco, 10888022), supplemented with 5% FBS (Sigma, 12106C), 1% B27 (Gibco, 17504-044), 1% GlutaMAX™ (Gibco, 35-050-038), and 50 U/ml penicillin/streptomycin). Further, digested tissue was gently triturated with a P1000 pipette, sieved through 70 µm cell sieve (Sarstedt, 83.3945.070) and 40 000 cells were seeded in a plating medium on previously coated 18 mm glass coverslips with 0.1 mg/ml poly-L-lysine (PLL; Sigma, P6282). Note for RNA extraction 200 000 cells were seeded on 25 mm coverslips. Lastly, after 4-5 h post seeding plating medium was exchanged with feeding medium devoid of serum (NB-A supplemented with 1% B27, 1% GlutaMAX™, and 50 U/ml penicillin/streptomycin) in which neuronal culture was maintained at 37°C temperature and 5% CO_2_ for a minimum of 14 days in vitro (DIV) prior to next experiments.

#### Rat primary culture

Primary neuronal cultures were obtained from hippocampi of P1 rats (*Rattus norvegicus*) as previously described ^[75]^ ^[76]^. Cells were then grown in 12-well plates on 18 mm glass coverslips in Neurobasal-A medium (Gibco, Paisley, Scotland), pH 7.5, at 37°C in 5% CO_2_. Transfections were performed at DIV 3-7 using Lipofectamine™ 2000 Transfection Reagent (Invitrogen, Carlsbad, CA, USA) following a protocol according to the recommendations of the manufacturer with modifications as previously described ^[77]^.

### Synaptic vesicle preparation

Synaptic vesicles (SVs) were isolated as described in previous studies ^[78]^ ^[79]^ ^[80]^. In brief, 20 rat brains dissected from adult male rats were homogenized in ice-cold sucrose buffer (320 mM sucrose, 4 mM HEPES-KOH, pH 7.4 supplemented with 0.2 mM, phenylmethylsulfonylfluoride and 1 mg/ml pepstatin A. Cellular debris were removed by centrifugation (10 min at 900× g), after which the obtained supernatant was subjected to 10 min centrifugation at 12,000× g. Next, synaptosome pellet was washed by resuspending it in sucrose buffer followed by centrifugation step for 15 min at 14500 × g. Synaptosomes were lysed by hypo-osmotic shock. From synaptosomes released SVs were fractionated by centrifugation of the lysate for 20 min at 20,000× g. The supernatant with SVs was further ultracentrifuged for 2 h at 230 000× g to obtain a crude synaptic vesicle pellet. SVs were purified by resuspending the pellet in 40 mM sucrose followed by centrifugation for 3 h at 110 880× g on a continuous sucrose density gradient (50–800 mM sucrose). Next, SVs were collected from the gradient and subjected to size-exclusion chromatography on controlled pore glass beads (300 nm diameter), equilibrated in glycine buffer (300 mM glycine, 5 mM HEPES, pH 7.4, adjusted using KOH), to separate synaptic vesicles from residual larger membrane contaminants. SVs were pelleted by centrifugation for 2 h at 230 000× g and resuspended in sucrose buffer by homogenization before being aliquoted into single-use fractions and snap frozen in liquid nitrogen. All the steps of SV prep were performed at +4°C.

### Pharmacological treatments

RNase A treatment of rat primary neurons was performed at DIV 10-14 using a modified photochemical internalization (PCI) method used primarily to promote endosomal release of macromolecules such as drugs, peptides, nucleic acids in cytoplasm ^[81]^. In our experimental set up PCI was applied to permeabilize the plasma membrane to enable an efficient RNase A delivery from surrounding medium into the cytosol of photochemically-activated neurons co-expressing mScarlet-synapsin-1 and synaptophysin-EGFP. Photosensitizer, aluminium phthalocyanine disulfonate - AIPcS_2a_, was used to trigger RNase A internalization. AIPcS_2a_ was added to primary culture at a final concentration of 10 ng/ml in calcium-free imaging medium (25 mM HEPES pH 7.35, 124 mM NaCl, 5 mM KCl, 1 mM MgCl_2_, 30 mM D-glucose, 5 mM EGTA) and incubated for 10 min to allow its integration into a cell membrane. Afterwards cells were washed with fresh imaging medium and supplemented with 150 µg/ml of RNase A (PureLink^TM^, Invitrogen, 12091-039). Next, to initiate membrane permeabilization, the region of interest was illuminated with red light for 10 s using an inverted Nikon Ti epifluorescence microscope (Nikon Corporation, Chiyoda, Tokyo, Japan) equipped with a Plan Apochromat 60×, 1.4 NA oil immersion objective, an HBO-100W Lamp and 620/60 ET Bandpass filter. After light activation neurons were incubated for 30 min. Image acquisition was performed immediately and 30 min post illumination. All the steps were performed under controlled conditions at 37°C. For BSA control experiments in neurons and method validation in HEK293 cell line the same workflow was applied.

For the validation of EIF4E antibody specificity, Na-arsenite (TRC-S080868) treatment of HEK293 cells was performed in order to promote stress granules formation, i.e., RNA enrichment in stress granules was used as positive marker. At 80% of confluency HEK293 cells were treated with arsenite to a final concentration of 330 μM for 30 min (5% CO_2_ at 37°C). In parallel no treated control cells underwent the same conditions without addition of arsenite.

### In silico RNA structure prediction and ensemble diversity analysis

To calculate the ensemble diversity (ED) of sequences identified in the Epple et al. ^[51]^ synaptic transcriptome, gene names corresponding to the study were downloaded from the supplemental materials sections and referenced to transcriptome datasets for each. For the Epple et al. dataset, the gene names in table S1 in the supplementary file ‘12035_2021_2296_MOESM2_ESM.xlsx’ were referenced to ReqSeq mRNA file refMrna.fa.gz from the mm10 mouse genome dataset (http://hgdownload.soe.ucsc.edu/goldenPath/mm10/bigZips/) from which RNA sequences were retrieved. The gene2refseq data was downloaded from (https://ftp.ncbi.nlm.nih.gov/gene/DATA/README) to link accession numbers to gene names. The python script ‘epple-el-al_mouse_transcriptome_filter.py’ implemented the described tasks. The ED for each sequence was calculated using the python script ‘epple-et-alS1_transcriptome_properties.py’. The script uses the ViennaRNA secondary structure prediction software python plugin to calculate the ensemble diversity of each sequence at 37℃^[48]^. ED values for *CLN3* 362 and *CLN3* 657 were obtained from^[46]^. Results were plotted using the custom python script.

### In vitro reconstitution assays

#### Total neuronal RNA and synapsin-1

Total neuronal RNA and purified EGFP-synapsin-1 (later referred as synapsin-1) were used for in vitro reconstitutions assays with RNA and synapsin-1 varying concentrations (see phase diagram in S1) in 10 μl-reaction buffer: 150 mM NaCl, 25 mM Tris-HCl pH 7.4 and 0.5 mM TCEP (referred further as “the reaction buffer” and used for all the in vitro reconstitutions except for reconstitutions with *CLN3* mRNAs).

#### In vitro *assays with orthogonal CLN3 mRNAs*

Synapsin-1 condensates were formed with *CLN3 RNA* by the addition of 2.5 µM synapsin-1 to reaction buffer (50 mM HEPES, 150 mM KCl) containing 35 nM *CLN3-Cy5*. The *RNA* concentration used for these reconstitutions was determined by balancing *Rattus norvegicus* synapsin-1 charge at pH 7.4 (+20.523)) with the negative charge provided by the length of phosphate backbone of the RNA (-1588 for *CLN3*). Reactions were incubated for 10 min prior to imaging. For the in vitro reconstitutions with pre-formed protein droplets containing synapsin-1/αSyn, the final concentration of proteins in the reaction mix were 4.5 µM and 1.5 µM, for synapsin-1 and αSyn respectively, maintaining a physiological ratio of _3:1_^[40]^.

#### In vitro *assays with synaptic mRNAs*

To form synapsin-1 condensates with synaptic mRNAs, *Cy3*-labelled *Prnp*, *Scg3* or *Atp5b* were mixed with synapsin-1 to a final concentration of .5 μM synapsin-1 and 42 nM *Scg3* or *Prnp* RNA or 49 nM of *Atp5b* resulting in a polyelectrolyte protein:RNA ratio of 1:1. In vitro reconstitutions with *Scg3* and *Prnp* RNAs together with synapsin-1 and αSyn were performed in two different set ups: *type I* and *type II* reactions. For the *type I* reactions protein-protein condensates of EGFP-synapsin-1 and αSyn-AF647 were pre-formed. upon which mRNA was added approx. 7 min post protein-protein condensate formation. Images were acquired before and after 15 min of RNA addition (25 min post protein-protein condensate formation) to assess the enrichment of proteins in the condensates. RNA enrichment was analyzed almost immediately upon RNA addition and after approx. 15 of incubation. In *type II* reactions first purified EGFP-synapsin-1 was mixed with RNA. Next, after 1 min of incubation in the tube αSyn was added and 10 µl reaction was aliquoted onto glass bottom dish for further imaging. 5- and 25-min post reaction mix enrichment of synapsin-1, αSyn and RNA was assessed. In both reaction set-ups synapsin-1 and αSyn were mixed at the ratio close to physiological range^[49]^ to a final concentration of 4.5 µM and 1 µM, respectively. *Scg3* or *Prnp* (each to a final concentration of 42 nM) were added in 1:1 polyelectrolyte charge ratio to synapsin-1 (*Rattus norvegicus*, +20.523 at pH 7.4). All the in vitro reconstitutions with synaptic RNAs were performed in the reaction buffer.

#### RNase A *in vitro* treatments

First 10 µM to a final concentration of EGFP-synapsin-1 and 300 ng of total neuronal RNA were mixed in 10 µl of the reaction buffer. Approximately after 6 min when majority of condensates reached around 3 µm in diameter time laps was started. After 2^nd^ lap acquisition was paused and RNase A was added to a final concentration 0.1 or 1 µg/µl. Almost immediately acquisition was continued for the next 10 min of RNase A incubation time.

In independent reaction set up 300 ng of total neuronal RNA was pre-digested with 1 µg/µl of RNase A for 10 min. Such pre-digested RNA was mixed with EGFP-synapsin-1 to a final concentration of 10 µM in the reaction buffer and the effect on the formation of synapsin-1/RNA condensates was assessed.

#### In vitro *assays with SVs*

In vitro reconstitutions of SVs were performed according to published protocol^1^. Briefly, first condensates of 10 µM EGFP-synapsin-1 and 300 ng of total neuronal RNA were formed. When condensates reached a size of approx. 3-5 µm FM4-64-labelled SVs (to a final concentration of 3 nM) were added onto synapsin-1/RNA condensates.

#### Dextran assays

Dextran assays were carried out following the previously published procedure^[42]^. Briefly, the commercially available TMR-dextran of specific molecular weight (Sigma Aldrich T1037, T1162, T1287) were added to PEG-driven or RNA-driven synapsin-1 phases. In brief, 4.5 µM synapsin-1 phases were formed either with 3% PEG 8K (Roth, 263.1) or 150 ng total neuronal RNA in 10 µl of the reaction buffer. Next, TMR-dextran of specific molecular weight to a final concentration of 0.04 mg/ml was added to the reaction mixture when the condensates reached an average diameter of 2 - 3 µm (approx. after 7 min). Enrichment or exclusion of TMR-dextran in synapsin-1 phases was followed approx. for 15 min post dextran addition.

### *Atp5b* smFISH probe design and labelling

*Atp5b* smFISH probes were designed applying the smFISH finder tool from Gaspar et al., 2017. For the probe design *Mus musculus* transcript of *Atp5b,* (NM_016774.3, see **S12D**) was used as template. Thirty 18-20mer oligonucleotides were designed and 3’-end labelled with amino-11-ddUTP conjugated to NHS-ester ATTO647N (ATTO-TEC Gmbh, AD 647-31), upon which labeled *Atp5b* 30× probe set (see Supplementary **Table S4)** was purified with ethanol precipitation. Probe labelling and purification was performed as previously described^[82]^.

### SmFISH and immunocytochemistry in neurons

Prior to *Atp5b* smFISH and ICC primary mouse neurons were washed gently with 1× PBS and fixed in 4% PFA, 4% Sucrose, 1 mM MgCl_2_ and 0.1 mM CaCl_2_ in 1× PBS for 20 min at room temperature. Permeabilization was performed for 3 min in 1× PBS supplemented with 0.1% of Triton-X 100 (PBT). Afterwards, neurons were pre-hybridized in hybridization (Hyb) buffer (300 mM NaCl, 30 mM sodium citrate pH 7.0, 15 v/v% ethylene carbonate, 1 mM EDTA, 50 μg/ml heparin, 100 μg/ml salmon sperm DNA, 1 v/v% Triton X-100)) for 20 min at 37°C, followed by 30× *Atp5b* probe mix addition (12.5-15 nM per individual oligonucleotide) and hybridization for 4 h at 37°C in humidified chamber. After hybridization step subsequent washes were performed: 1× 12-min wash in Hyb-buffer at 37°C, followed by 2× 7 min washes in Hyb-buffer: PBT (1:1) and PBT at room temperature. Next, neurons were subjected to blocking in 1× β-casein supplemented with 0.1% Triton-X 100 for 20 min at room temperature. Primary antibody incubation (MAP2 and vGLUT1 antibodies at a dilution 1:1000 and 1:500, respectivelly) was done for 30 min at room temperature in the same buffer as used for the blocking step. Next, 3 consecutive washes were performed (each 7 min in PBT). Secondary antibodies (anti-chiken AF488 and anti-rabbit AF568 both at a dilution 1:500) were incubated for 30 min under the same buffer and temperature conditions, followed by 3× 7-min washes in PBT. Lastly, neurons were mounted in Fluoroshield supplemented with DAPI (Sigma Aldrich F6057). The antibody list and concentration in use is provided in Supplementary **Table S5**.

### Poly(A) FISH and immunocytochemistry in neurons and HEK cells

For poly(A) FISH commercially available Stellaris probe was used (T30-ATTO647N-1, stock concentration 12.5 µM). Probe was diluted 1:100 in Hyb-buffer and hybridized overnight at 37°C in humidified chamber. Pre-hybridization step and post-hybridization washes as well as post-FISH immunostaining and sample mounting were performed as described above.

### Ectopic expression of SV-like condensates and immunocytochemistry in HEK cells

HEK293 cells ectopically expressing synapsin-1 condensates (plasmid set: 6×His-TEV-mCherry-PP-hsSYN1 + pCMV SYPH-FL) or synapsin-1/α-synuclein condensates: (plasmid set: 6×His-TEV-mCherry-PP-hsSYN1 + pCMV SYPH-FL + pCMV hsSNCA) were fixed 20 h post-transfection in 4% PFA, 4% Sucrose, 1 mM MgCl_2_ and 0.1 mM CaCl_2_ in 1× PBS for 15 min. Fixation was followed by 3-5 min permeabilization in PBT and subsequent blocking in 1× β-casein supplemented with 0.1% Triton-X 100 for 30 min. The same protocol was applied for untransfected HEK cells which underwent poly(A) FISH prior to immunostining. Primary antibody incubation (EIF4E antibody at a dilution 1:250) was done for 2 h at room temperature in a blocking buffer, after which 3 consecutive 10-min washes were executed in PBT. Secondary antibody (anti-rabbit STAR488 at a dilution 1:500) was incubated for 60 min under the same buffer and temperature conditions, followed by 3× 10-min washes in PBT. Next, cells were mounted in Fluoroshield supplemented with DAPI. The antibody list and concentration in use is provided in Supplementary **Table S5**.

### Luciferase in vitro translation assay

For luciferase assays, *Atp5b-NLUC* and synapsin-1, or synapsin-1-RtoK were pre-mixed at 70 nM and 9 µM respectively and incubated for 10 min at room temperature prior to the dilution of the reaction 1:1 with Rabbit Reticulocyte extract (RRE; Promega, L4960). The reactions were incubated at 30°C for 1.5 hours before the addition of substrate, Vivazine (Promega, N2580). Vivazine was diluted 1:50 in translation buffer (50 mM Hepes at 7.4, 140 mM KCl, 2 mM magnesium glutamate), and 15 µl of substrate was added for every 20 µl of luciferase reaction. The luminescence for each reaction was read out on a Promega GloMax® Explorer with a 1 s integration time. For reactions containing, α-synuclein, 3 µM α-synuclein was added during the pre-mix. For reactions containing BSA, BSA was diluted in translation buffer (50 mM Hepes at 7.4, 140 mM KCl, 2 mM magnesium glutamate) and added to the pre-mix step to a final concentration of 9 µM.

### Moon-Tag-spatial in vitro translation assay

The MoonTag system *in extract* was adapted from^[70]^. Briefly, mixtures of synapsin-1 (or synapsin-1 and α-synuclein) and *Atp5b-MOONTAG RNA* were pre-mixed as described in the Luciferase assay and left to incubate for 10 min at room temperature. After incubation, RRE containing was added to the mixture in a ratio of 1:1 along with 2H10-mCherry, to a final concentration of 500 nM. MoontTag reactions were left to incubate for 1 hour at 30°C prior to visualization.

### Correlative light and electron microscopy

Upon 24 h post-culturing HEK293 cells on poly-L-lysine coated 6 mm sapphire discs, the cells were transiently transfected with plasmids coding for synapsin-1-GFP and synapthophysin using jetPEI® DNA transfection reagent (Polyplus) according to manufacturer’s protocol. Briefly, a ratio of jetPEI®:DNA of 3 µL: 1 µg for 24 h was used. The discs were covered with a gold-plated 6 mm type-B carrier containing 20% BSA in PHEM buffer, which served as a cryoprotectant. Cells were cryo-fixed using a high-pressure freezer (EM ICE; Leica). Freeze substitution was performed in acetone supplemented with 0.2% uranyl acetate and 2% H₂O for 60 h at −90°C and 12 h at −40°C using an automated freeze substitution system (AFS2; Leica). Samples were washed with ethanol and embedded in HM20 resin at −40°C. Polymerization was carried out under UV light for 48 h at −40°C followed by 48 h at 20°C.

Ultrathin sections (200 nm) were cut using an ultramicrotome (UC6; Leica) and collected on formvar-coated NiH100 grids. Sections were stained with DAPI and imaged in PBS using a fluorescence microscope (Axio Observer Z1; Carl Zeiss) equipped with ZEN software and a Prime 96B digital camera (Photometrics). Z-stack images were acquired using a 100×/1.30 NA objective (Carl Zeiss). GFP fluorescence was detected using a 470/40 nm bandpass excitation filter, a 495 nm dichroic mirror, and a 525/50 nm bandpass emission filter. DAPI fluorescence was detected using a 365 nm excitation filter and a 445/50 nm bandpass emission filter.

Following fluorescence imaging, grids were post-stained with uranyl acetate and lead citrate and labeled with 10 nm gold particles. Regions of interest identified from fluorescence data were imaged by transmission electron microscopy (TALOS 120C; Thermo Scientific) using MAPS software. Fluorescence and TEM images were correlated using the eC-CLEM plugin (Paul-Gilloteaux et al., 2017) in Icy (http://icy.bioimageanalysis.org). Tomograms were reconstructed and rendered in 3D using the IMOD software package. Ribosome density was quantified using FIJI. Cubic volumes of 100 nm³ were defined with the rectangle selection tool, and the number of ribosomes within each selected volume was counted.

### Microscopy

All the images of in vitro reconstitutions of synapsin-1 phases with total neuronal RNA or synaptic RNAs ± α-synuclein were acquired with Nikon Spinning Disk Confocal CSU-X with emission filters 452/45, 525/50, 600/50, 685/40 using iXon3 DU-888 Ultra EMCCD camera detector (1024×1024-pixel resolution, 13 μm pixel size, 26 fps full frame, usable area ∼512×512 pixels, 93 fps) and 60× oil objective (Plan Apo, NA = 1.4, WD = 0.13 mm).

Imaging of in vitro reconstitutions of synapsin-1 condensates with *CLN3* RNAs ± α-synuclein were obtained on a Nikon Yokogawa CSU-W1 SoRa spinning disk confocal (confocal mode) equipped with an ORCA-Fusion BT Digital CMOS camera using a 100× oil objective (CFI Plan Lambda, NA = 1.45, WD = 0.13 mm). Scaling factor = 15.39 pixels/µm.

For MoonTag reactions, 3D fluorescence microscopy was performed on a Nikon Ti2-E inverted microscope equipped with a Yokogawa CSU-W1 spinning disk confocal and an ORCA-Fusion BT Digital CMOS camera. Images were collected using a 100× oil objective (CFI Plan Lambda, NA = 1.45, WD = 0.13 mm) with Nyquist sampling.

Mouse neurons and HEK293 cells expressing SV-like condensates were imaged with Nikon Spinning Disk Confocal CSU-X with emission filters 452/45, 525/50, 600/50, 685/40 using pco.edge 4.2 bi sCMOS camera detector (2048×2048 pixel resolution, 6.5 μm pixel size, 40 fps full frame, usable area 1024×1024 pixels) and 60× oil objective (Plan Apo , NA = 1.4, WD = 0.13 mm). Images were acquired in 12-bit mode. Exposure and laser intensities were adapted to antibodies and FISH signal requirements. Same imaging set up as above was applied rat neurons to assess poly(A) signal.

Non-transfected arsenite-treated HEK293 cells were imaged on Nikon Laser Scanning Confocal A1Rsi+. Images were collected using galvano scanner together with PMT and hybrid GaAsP PMT detectors and 60× oil objective (Plan Apo, NA = 1.4, WD = 0.13 mm) with Nyquist sampling.

### Data analysis, statistics and visualization

#### Electrostatic charge potential map of synapsin-1

Visualization of the electrostatic charge potential of synapsin-1 (*Rattus norvegicus*) was carried out according to these steps: First, synapsin-1 3D structure was downloaded (AF-P09951-F1-v4, PDB). To compute the atomic structure of synapsin-1 with the associated charges, the syanpsin-1 PDB file was read into PDB2PQR (available at https://www.poissonboltzmann.org/), where calculations were carried out at pH 7.4 (PROPKA protonation states) with a “PARSE” forcefield. Subsequently, files generated from PDB2PQR calculations were read into APBS (available at https://www.poissonboltzmann.org/) for final calculations of synapsin-1 electrostatic densities. To assemble the synapsin-1 electrostatic density map, the electrostatic potential was aligned with the surface rendering of synapsin-1 using ChimeraX (https://www.cgl.ucsf.edu/chimerax/).

#### Phase diagrams and synapsin-1 charge / total neuronal RNA charge ratio calculation

To obtain polyelectrolyte ratio of synapsin-1 and total neuronal RNA charges used in phase diagram in vitro reconstitutions following equation was applied:

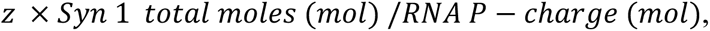

where z = (+)20.532 — synapsin-1 net charge. To obtain charge approximation of total neuronal RNA total moles of phosphate where used as equal to total moles of negatively charged nucleotides. For each amount of RNA used in reaction RNA P-charge is calculated as: *P* (*mol*) = *total neuronal RNA mass* (*g*)/ *average nucleotide MW* (*g*/*mol*), where av. nucleotide MW = 340 g/mol. Charge ratio histogram was plotted using RStudio (2023.03.0+386 “Cherry Blossom” release for Windows).

#### Normalized synapsin-1 fluorescence enrichment measurements of *in vitro* RNase A treatment over time

For measuring the intensity of condensates just before addition of RNase A and over time of RNase A treatment 10 individual condensates were selected from 3 independent reconstitutions. First, x-y shifts during the time-lapse acquisition were corrected for individual condensates using the StackRegJ plugin^[83]^. Next, a circular region of interest (ROI was created on condensates and their change in fluorescence intensity over time was computed using the create spectrum jru v1 plugin ^[83]^. To measure the change in fluorescence intensity of the dilute phase (out) 10 regions of interest (size: 29 by 25 pixels) for each replicate were selected outside of the condensate and the change in the fluorescence intensity of the dilute phase over time was computed using the create spectrum jru v1 plugin. Normalized synapsin-1 fluorescence enrichment was calculated as the ratio of mean intensity inside condensates to the average mean intensity outside of condensates (In/Out) over the course of time of 10 min. Data was collected from three independent in vitro reconstitutions and plotted using GraphPad Prism software v 9.1.

#### *Synapsin-1 fluorescence enrichment analysis for before and after RNase A treatment* in vitro

To quantify synapsin-1 fluorescence enrichment in the condensates a custom Fiji workflow was used (reference). In brief, all the acquired images were first pre-processed. An area of interest (size: 60 µm x 60 µm) was selected at the center of the image frame and an equatorial z-plane was manually selected (to minimize any bias from out-of-plane droplets). Such obtained data was further used for processing and analysis. For quantifying the fluorescence intensity within the synapsin-1 dense phase, condensates were segmented using the Otsu algorithm after Gaussian convolution and further mean intensity was determined within condensates. For determining the fluorescence intensity within the dilute phase 10 rectangular regions of interest (size: 29 by 25 pixels) for each reconstitution were selected outside of the condensates and the mean fluorescence intensity of the dilute phase was computed. Data was collected from three independent in vitro reconstitutions and plotted using RStudio (2023.03.0+386 “Cherry Blossom” release for Windows).

#### RNA and protein enrichment quantification of CLN3- and synaptic RNA-driven synapsin condensates

Image analysis of synapsin-1 condensates was carried out using a custom macro in Fiji. The macro performed the following operations: Each image-stack was first analyzed for the 2D-cross section which most closely approximated the midplane of synapsin-1 condensates. After selecting a z-plane for analysis, a list of ROIs corresponding to the condensates was generated using an Otsu threshold, and subsequently particle detection, on the synapsin-1 channel. The ROIs corresponding to the synapsin-1 condensates were overlaid on the fluorescence channels corresponding to synapsin-1, RNA, and αSyn, and intensities values for each ROI were collected. Raw fluorescence values extracted from Fiji were then loaded into Python 3.11.3 for wrangling, statistical analysis, and data visualization. Data were collected from three independent in vitro reconstitutions.

#### Synapsin-1 condensates *± SVs* diameter assessment

For assessing the diameter of RNA-driven synapsin-1 condensates with and without SVs acquired images were segmented using the Otsu algorithm after Gaussian convolution and further diameters of condensates that had circularity within the range of 0.9-1.0 were determined. Data were collected from three independent in vitro reconstitutions and relative frequency distribution (fractions) as a function of diameter was plotted using GraphPad Prism software v 9.1.

#### Line profiles assessment

A line of 3 µm for cells or 6 µm for reconstituted condensates was drawn across the region of interest and the fluorescence intensity profiles were recorded for each acquired channel using in Fiji made macro. The obtained data was normalized and plotted using GraphPad prism software v9.1.

#### FRAP data acquisition and analysis

During the FRAP experiments, ROI for photobleaching was set to 2 µm. FRAP movies were acquired in such way that prior to the bleach 6 images were acquired at 100 ms and 1 second interval for synapsin-1/RNA and synapsin-1/RNA ± SVs in vitro reconstitutions respectively. This step was followed by bleaching at maximum laser power intensity with 488 nm wave length on Nikon Spinning Disk Confocal CSU-X. After the bleach images were acquired at 100 ms or 1 s interval (depending on in vitro set up-see above) continuously for 10 min. In order to quantify the intensity of condensates over time, x-y shifts during the acquisition of FRAP movie were corrected for bleached condensates using the StackRegJ plugin^31^. Next, a circular ROI was created on bleached condensates and their change in intensity over time was recorded using the create spectrum jru v1 plugin^31^. A correction factor was calculated based on the change in fluorescence outside of the bleached ROI and applied to correct the intensities of bleached condensates over time. The corrected fluorescence intensities were further normalized based on the minimum and maximum intensity values and plotted using GraphPad Prism software v 9.1.

#### Image analysis of FISH + ICC in primary neurons and ICC in HEK cells

Images of neurons with *Atp5b* smFISH + ICC were analyzed using previously described pipeline ^[84]^. Images were analyzed using radial symmetry-FISH (RS-FISH) ^[84]^. MAP2 was used to generate a mask for neurons. *Atp5b* foci were detected using the RS-FISH Fiji plugin. Detected *Atp5b* foci were then used as reference to manually screen for co-localization with presynaptic marker vGLUT1 using Fiji (ImageJ).

For poly(A) FISH + ICC in neurons line profile analysis of fluorescence intensities was performed on maximum projections to manually asses co-localization of poly(A) FISH signal and immunostained vGLUT1 in axons. MAP2 was used as neuronal marker.

Images of HEK293 cells co-expressing SV-like condensates and immunostained for EIF4E. All condensates were manually tracked through whole z-stack to assess their co-localization with EIF4E. Representative line profiles of fluorescence intensities of mCherry-synapsin-1 and EIF4E (STAR488) were obtained from maximum projections of 2-3 z-planes of the widest condensate area of the region of interest. All imaging analyses were performed in Fiji (ImageJ) ^[85]^.

#### Luciferase assay and MoonTag image analysis

For luminescence data wrangling, visualization and statistical analysis, raw luminescence values were read into Python 3.11.3. To minimize the impact of RRE translation-potential variation on luminescence values collected, the reactions were performed as follows: for each luminescence condition (no synapsin-1, synapsin-1 only, synapsin-1 + αSyn, BSA), data were collected from six technical replicates per a condition, per an extract, for a total of three extracts or biological replicates. All four conditions were carried out in each extract.

For image-stacks collected of synapsin-1 condensates containing ATP::MoonTag RNA and mCherry-2H10Nb, intensity values corresponding to synapsin-1 and mCherry-2H10Nb were extracted using a custom Fiji macro described above in *RNA and protein enrichment quantification of CLN3- and synaptic RNA-driven synapsin condensates*. After image segmentation, intensity values were read into Python 3.11.3 for data wrangling, visualization and statistical analysis.

### Statistics and Reproducibility

The number of biological and technical replicates and the number of analyzed molecules are indicated in the figure legends and Source Data. Statistical tests were performed using R-studio and Prism 9 (Graphpad).

## Acknowledgements

We thank Advanced Medical Bioimaging Core Facility (AMBIO) at Charité University Clinic (Berlin, Germany); to Silvio O. Rizzoli (Göttingen) for the constructive advice for the RNase experiments. The work is supported the grants from the German Research Foundation (MI 2104 and SFB1286/B10), the ERC Grant MemLessInterface (101078172), the HFSP Grant (RGEC32/2023), and Horizons Europe MSCA DN Grant (101167843 - BICEPS) to DM; Air Force Office of Scientific Research FA9550-20-1-0241, NIH grant 7R01GM081506-13, and 1R35GM156800 to ASG; F32 GM151858-02 to ZMG; Innovative Minds Program of the German Dementia Association to CH.

## Conflict of Interests

The authors declare no competing financial interests.

## Author contributions

Conceptualization: DM, ASG, BR, ZMG.

Methodology: BR, ZMG, CH, SR, RdB, GvdB.

Investigation and Visualization: BR, ZMG, AC, IS, CH, SR, GAP, AnC, LTH.

Resources support: KF, MJ, FH and VMJ, KN, VD, MMM.

Writing—original draft: DM, ASG, BR and ZMG.

Writing—review and editing: all contributing authors.

Funding acquisition: DM and ASG.

Project administration: DM and ASG.

## Data Availability

Source data are provided with this paper. All data generated or analyzed for this study are available within the paper and its associated supplementary information files. All other data presented are available upon reasonable request from the corresponding author.

## Code Availability

All codes written for this study are available upon reasonable request from the corresponding authors.

## Notes

### Competing Interest Statement

The authors have declared no competing interest.

### Summary of Updates

This is a revised version of the manuscript with the new data and experiments to strengthen the conclusions of the study.

