## Supplementary Data for "RNA promotes synapsin phase separation providing a platform for local translation"

### Supplementary Data File includes:

- Supplementary Figures S1-S14;
- Supplementary Tables S1-S5.

### Supplementary Data Figures and Figure legends

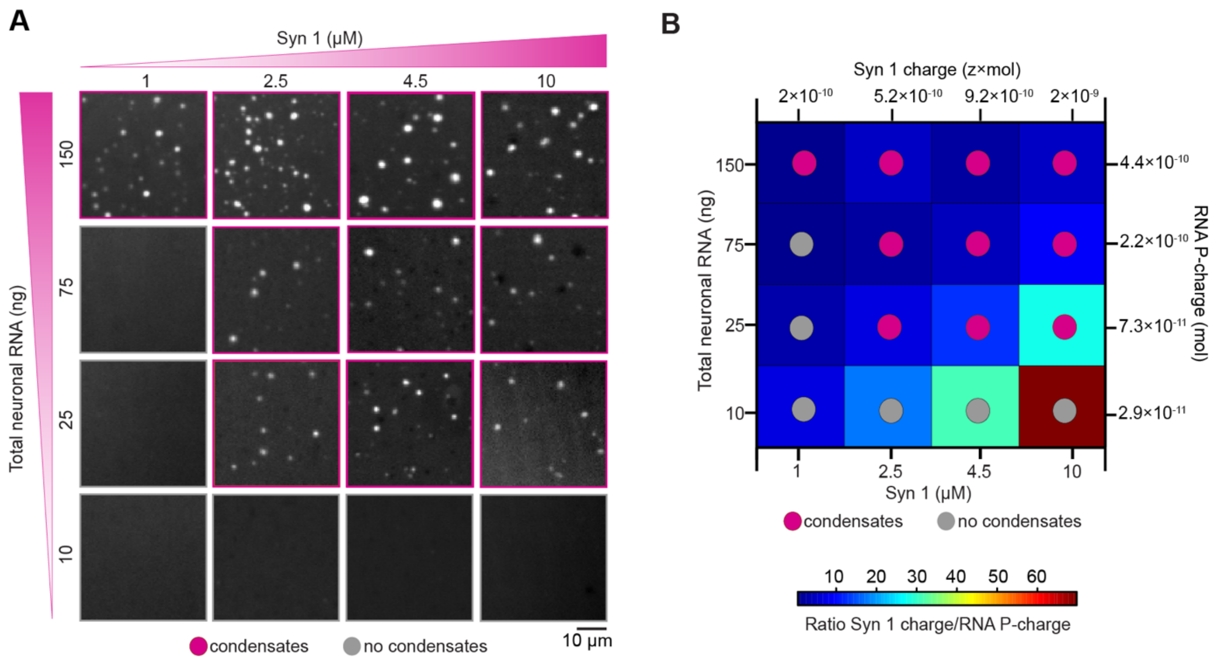

**Supplementary Figure 1| Phase diagram of RNA-driven synapsin-1 condensates.**
**(A)** Phase separation of purified recombinant EGFP-synapsin-1 full-length (Syn 1) protein was assessed with respect of increasing concentrations of protein and total neuronal RNA. *In vitro* reconstitution of 10 µl-reaction mix of Syn 1 and RNA was prepared under physiological salt conditions and in the absence of any molecular crowders (150 mM NaCl, 25 mM Tris-HCl pH 7.4 and 0.5 mM TCEP). These representative images correspond to the phase diagram plot in **Fig. 1B**. (**B)** Phase diagram as a function of total charge of synapsin-1, calculated as number of synapsin-1 moles in the reaction mixture multiplied with z (+20.523) for Syn 1, and RNA, calculated as a charge of the total neuronal RNA phosphate backbone charge. The colored histogram depicts the ratio of charges in Syn 1 and RNA.

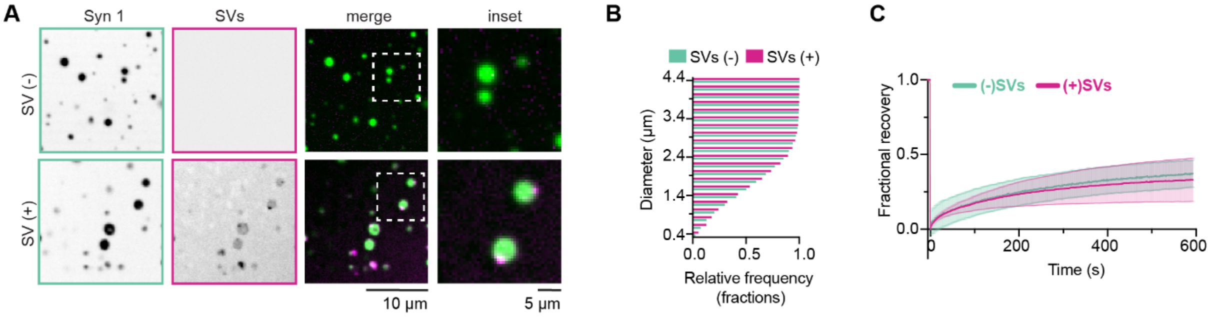

**Supplementary Figure 2 | Synapsin-1/RNA condensates recruit synaptic vesicles *in vitro*.**
**(A)** Representative micrographs of *in vitro* reconstitutions of 10 µM EGFP-synapsin-1 (Syn 1, in green) with 300 ng total neuronal RNA condensates with (+) and without (-) SV addition. To visualize SVs membrane staining FM4-64 styryl dye was used (in magenta).
**(B)** Quantification of the diameter of Syn 1/RNA condensates with (+) and without (-) SV addition. The plot represents frequency distribution of Syn 1 condensates diameter with (+) or without (-) SVs.
**(C)** FRAP curves of Syn 1/RNA condensates (+) or (-) SVs. Fluorescent recovery after photobleaching was recorded for 600 s with an interval of 1 s. No significant effect was observed on the Syn 1/RNA condensates dynamics upon addition of SVs. Fractional recovery ±SD was plotted as a function of time. Data were collected from three independent FRAP experiments.

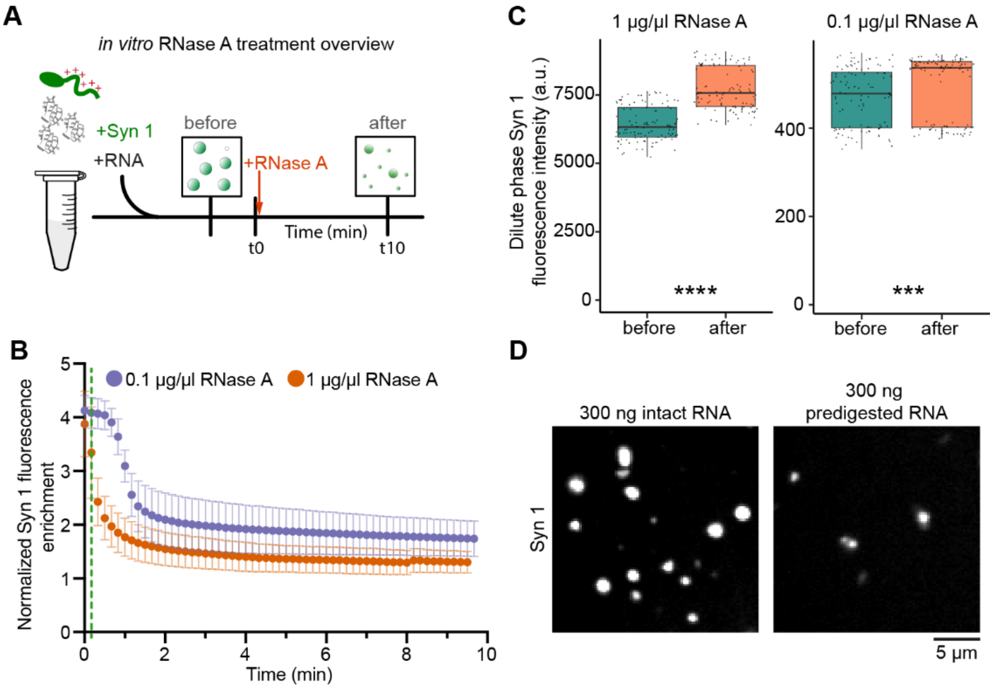

**Supplementary Figure 3 | RNase A treatment of synapsin-1/RNA condensates.**
**(A)** Schematic representation of *in vitro* RNase A assay workflow.
**(B)** Quantification of EGFP-synapsin-1 (Syn 1) Syn 1 fluorescence enrichment in Syn 1/RNA condensates (10 µM Syn 1 to a final concentration and 300 ng of total neuronal RNA) just before and upon RNase A addition (to a final concentration of 0.1-green, and 1 µg/µl-magenta) was assessed in the course of 10 min incubation time. Partition coefficient (±SD) of Syn 1 for 10 manually selected condensates per technical replicate for each condition was analyzed. Red dashed line indicates first value of fluorescence intensities of Syn 1 condensates quantified immediately upon RNase A addition. Data were collected from three independent *in vitro* reconstitutions (total 30 selected condensates). This quantification corelates with representative images of selected condensate in **Fig. 1D**.
**(C)** Quantification of Syn 1 fluorescence intensity of a dilute phase before and 10 min after the treatment with 1 (left) or 0.1 µg/µl (right) of RNase A (complementary to data in **Fig. 1E**). The jitter box plot encompasses 25th and 75th quartiles with median marked by central line. Minimum and maximum of quantified Syn 1 fluorescence intensities are denoted by whiskers. Black circles (above the whiskers) represent quantified fluorescence intensities outside the range of adjacent values. Each point represents the value, which was slightly shifted for better visualization of overlapping data points. **** indicates $P \leq 0.0001$, *** $P \leq 0.001$ by Wilcoxon signed-rank test. Data was collected from three independent in vitro reconstitutions.
**(D)** Representative micrographs indicating that 10 µM Syn 1 readily forms condensates with intact RNA (left) but fails to phase separate in the presence of nucleotides (i.e., predigested RNA with 1 µg/µl of RNase A; right).

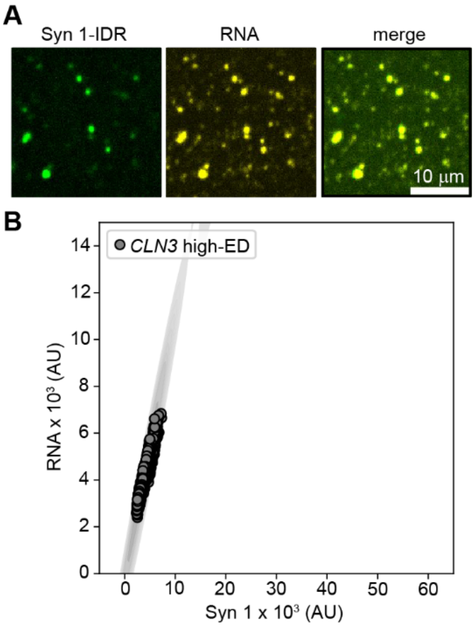

**Supplementary Figure 4 | Syn 1-IDR can recruit more unstructured mRNAs *in vitro*.**
**(A)** Representative images of *in vitro* reconstituted EGFP-synapsin-1-IDR (Syn 1-IDR) condensates (green) enriched with Cy5-labelled high-ED *CLN3* mRNA (yellow) of *A. gossypii*.
**(B)** Plot showing fluorescence intensities quantification of colocalizing Syn 1-IDR and *CLN3* high-ED in corresponding Syn 1-IDR/*CLN3* high-ED condensates. In the scatter plot overlayed with Kernel Density Estimate (KDE) each circle represents Syn 1-IDR/RNA condensate analyzed. Data were collected from three independent *in vitro* reconstitutions.

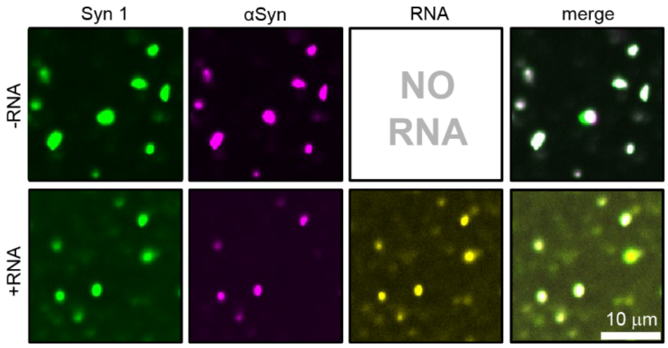

**Supplementary Figure 5 | In vitro reconstitution of synapsin-1 and α-synuclein with *CLN3* high ED mRNA.**
Representative micrographs of *in vitro* reconstituted synapsin-1/α-synuclein (Syn 1/αSyn) condensates with or without *CLN3* high ED mRNA upon 15 min of incubation under physiological salts conditions and in the absence of molecular crowders.

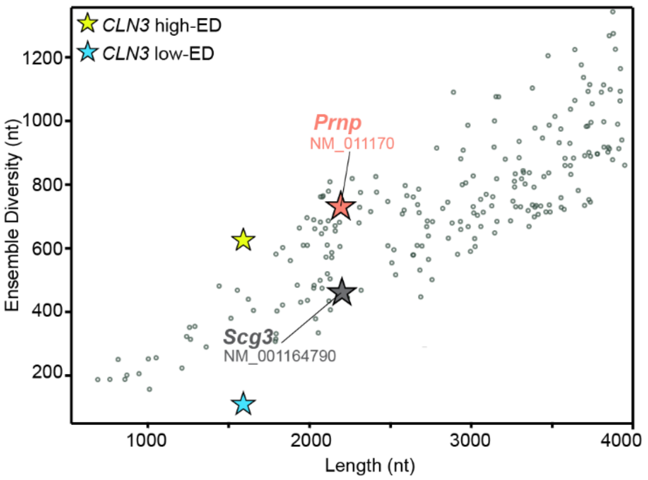

**Supplementary Figure 6 | ED signature of the synaptic RNAome data set.**
Ensemble diversity (ED) analysis plot showing synaptic transcripts from [57] S1 data set in correlation with *CLN3* low-ED and *CLN3* high-ED *Ashbya gossypii* mRNAs. *Prnp* and *Scg3* mRNAs (highlighted in the plot with a orange and grey star, respectively) were used in the *in vitro* reconstitution experiments of **Fig. 3**. The scatter plot shows transcript ED values as a function of transcript length. Each circle represents an individual mRNA. nt stands for nucleotides.

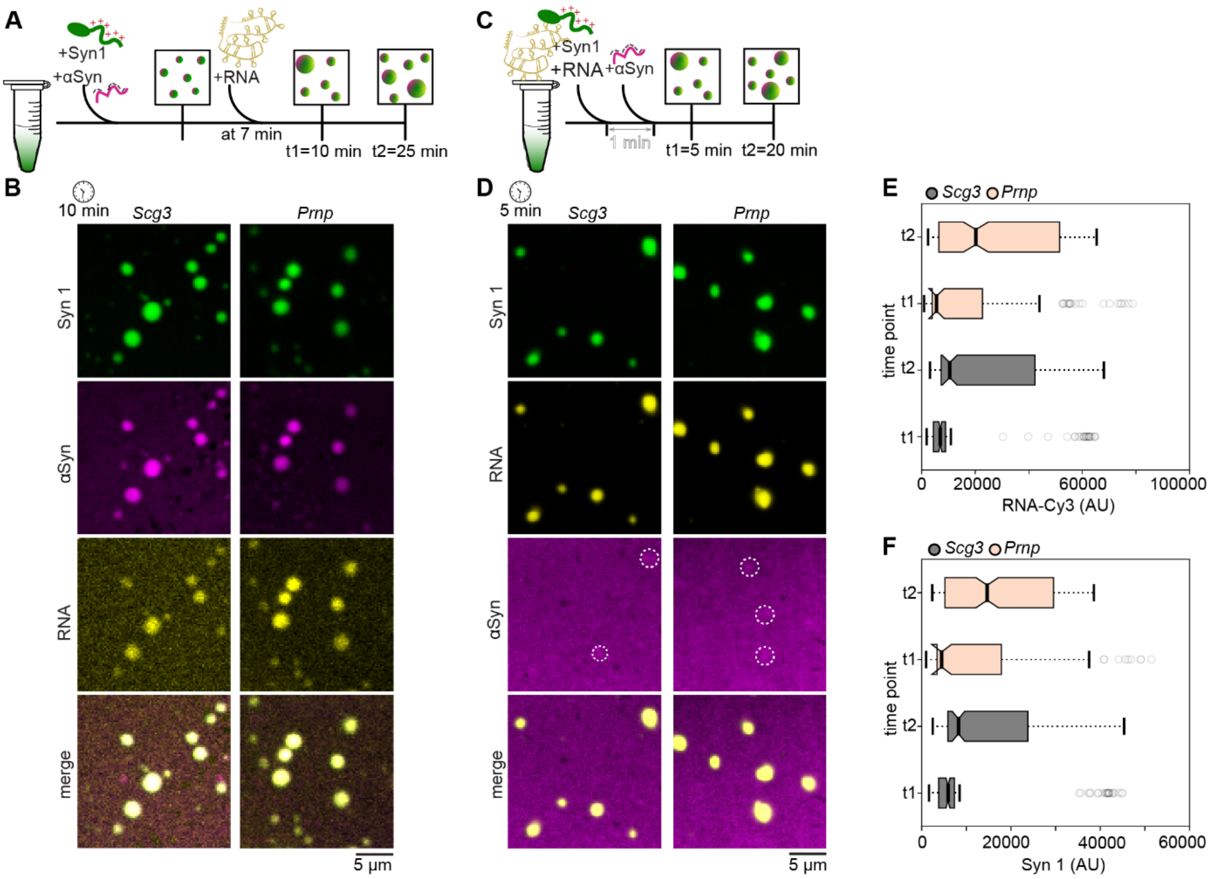

**Supplementary Figure 7 | Synaptic mRNAs affect α-syn recruitment into synapsin-1 condensates.** **(A)** Representative scheme of type I *in vitro* reaction set up (see Methods): First protein-protein condensates of synapsin-1 (Syn 1) and α-synuclein **(**αSyn) were formed and loaded on glass bottom dish for subsequent imaging. After approx. 7 min mRNA of interest was added precisely into the reaction droplet. Next, for mRNA and protein enrichment assessment images were taken 10- and 25-min post proteins mixing step.

**(B)** Representative images of type I *in vitro* reaction set up at 10 min after synaptic mRNA (*Scg3* or *Prnp*) addition.

**(C)** Representative scheme of type II *in vitro* reaction set up (see Methods): First Syn 1 and mRNA of interest were pre-mixed in the reaction tube and incubated for 1 min. Next, αSyn was added and the reaction mix was aliquoted on the glass bottom dish for subsequent imaging. For mRNA and protein enrichment assessment images were taken 5- and 25-min post reaction set up.

**(D)** Representative images of type II *in vitro* reaction set up at min 5 after mixing all the reaction components.

**(E)** Quantifications of mRNA enrichment (*Scg3* or *Prnp*) in *in vitro* reconstituted RNA-protein droplets at time points t1 and t2 (reaction set up type II, see Methods). Box plots (e, f) encompass the $25^{th}$ and $75^{th}$ quartiles with the median marked by central line. Whiskers extend to 1.5 times the interquartile range (IQR). Circles indicate outliers. Paired two-tailed T-test comparing the intensities of RNA within condensates at t1 and t2 results in a P-value of <0.00001 for reactions containing either *Scg3* or *Prnp*.

**(F)** Quantifications of Syn 1 enrichment in *in vitro* reconstituted RNA-protein droplets at time points t1 and t2 (reaction set up type II, see Methods). Paired two-tailed T-test comparing the intensities of Syn1 within condensates at t1 and t2 resulted in a P-value of 0.000165 and 0.000168 for reactions containing *Scg3* and *Prnp*, respectively. Data were collected from three independent *in vitro* reconstitutions. Data presented here supplements *in vitro* data from **Fig. 3**.

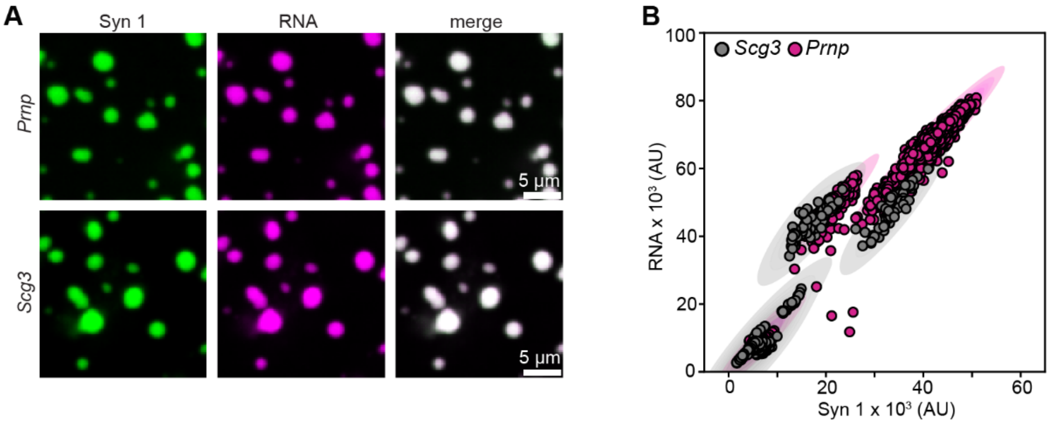

**Supplementary Figure 8 | Synaptic mRNAs phase separates with synapsin-1 *in vitro*.**
**(A)** Representative images of *in vitro* reconstituted EGFP-synapsin-1 (Syn 1) condensates with Cy3-labelled synaptic mRNAs (*Prnp* or *Scg3*) after approx. 20 min of incubation upon reaction set up in the reaction buffer with physiological salt conditions and the absence of any molecular crowder (see Methods).
**(B)** Quantification of RNA fluorescence intensities (*Scg3* in grey and *Prnp* in magenta) as a function of Syn 1 fluorescence intensities. In the scatter plot overlayed with Kernel Density Estimate (KDE) each circle represents Syn 1/RNA condensate analyzed. Data were collected from three independent *in vitro* reconstitutions.

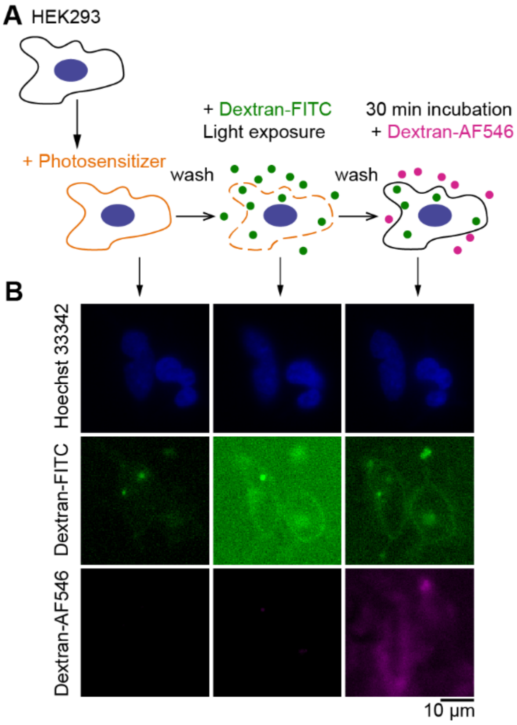

**Supplementary Figure 9 | Validation of PCI method in mammalian cells.**
**(A)** Schematic representation of validation of the PCI method in HEK293 cell line.
**(B)** Representative images of HEK293 cells at each step of PCI. Dextran-FITC (green) and dextran-AlexaFluor (AF)546 (magenta) were used to assess internalization of small molecules and their retention 30 min post photo-activation, as well as membrane sealing following photo-activation. Nuclei (blue) were stained with Hoechst 33342.

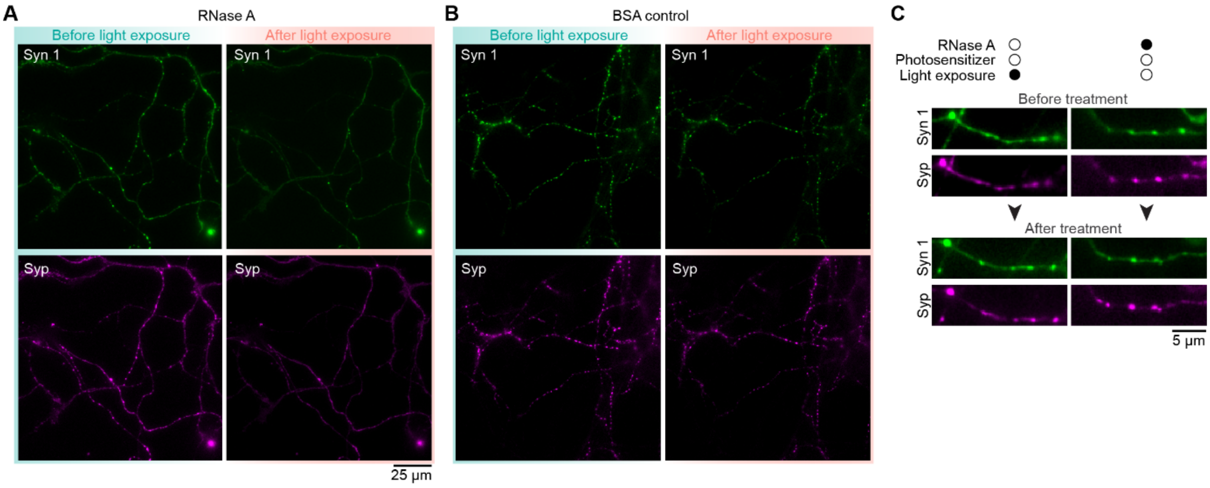

**Supplementary Figure 10 | PCI mediated RNase A treatment in primary rat neurons.**
**(A)** Representative micrographs of full frame axonal network before and after RNase A treatment and **(B)** before and after BSA treatment showing mScarlet-synapsin 1 and EGFP-synaptophysin signal distribution at the synaptic boutons. These images correspond to the axonal segments in **Fig. 4**.
**(C)** Representative micrographs of axonal segments before and after treatment by light exposure (left) or incubation with RNase A (right) in the same experimental conditions as in results presented in **Fig. 4**, but in the absence of the photosensitizer. Black circles indicate presence of the specific condition while white circles indicate the absence of the same.

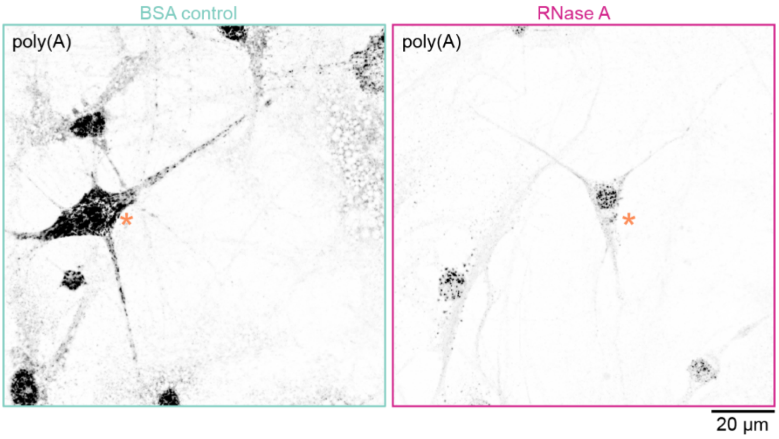

**Supplementary Figure 11 | Quality control of PCI mediated RNase A treatment in primary neurons.** Representative images for quality control of RNase A treatment in primary rat neurons showing that poly(A) signal was RNase A-sensitive. Left: image shows poly(A) FISH signal of neurons treated with BSA for 30 min. Right: image shows primary neuron underwent RNase A treatment for 30 min to a final concentration of RNase A of 150 µg/ml. Orange asterisks indicate neuronal soma. Glial cells could be detected in the background with noticeable poly(A) signal depletion in treated example. Images of BSA and RNase A-treated neurons were acquired under the exact same imaging conditions.

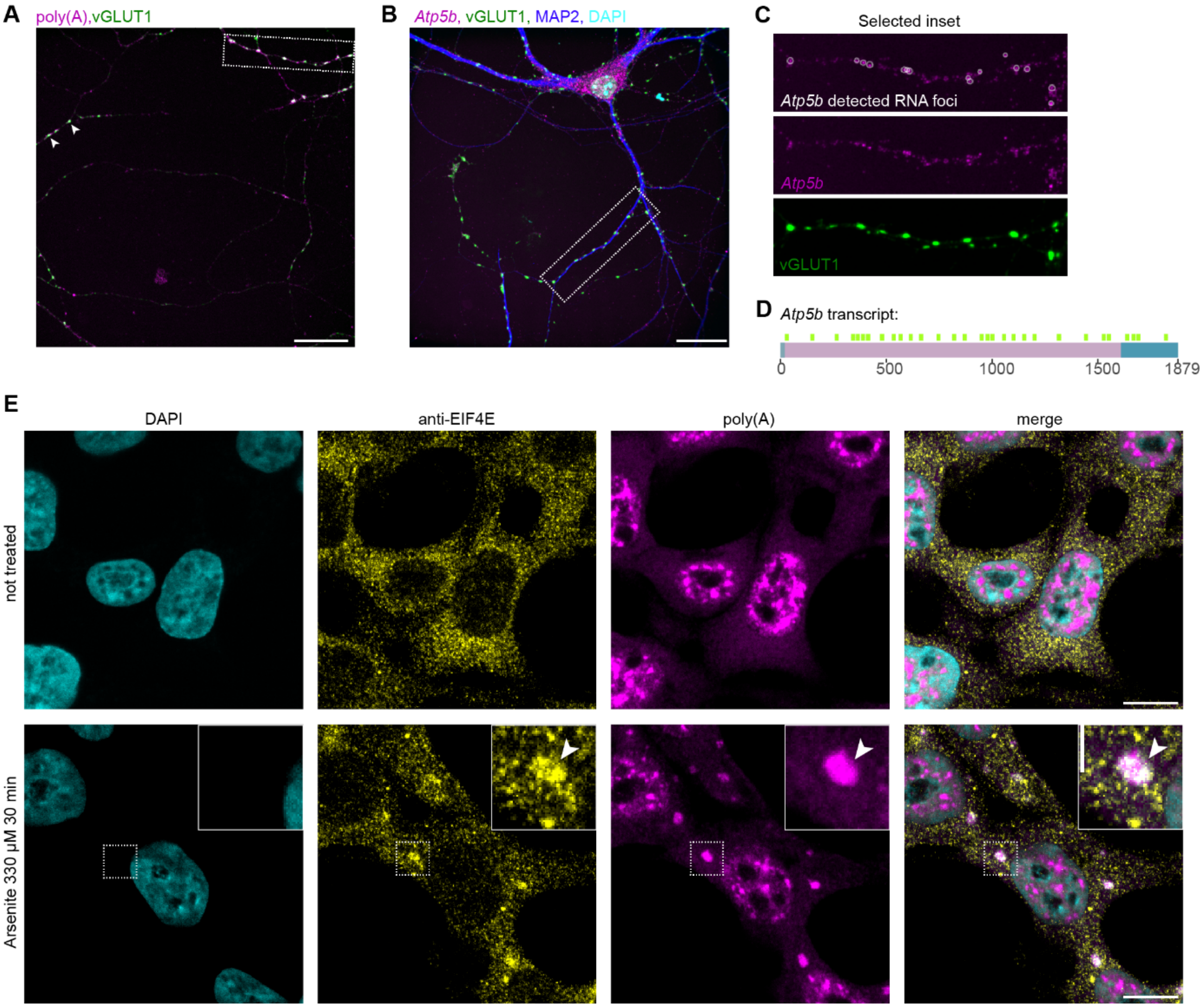

**Supplementary Figure 12 | Synapsin-1 condensates harbor mRNAs and EIF4E.**

**(A)** Representative full frame image of hippocampal axon showing poly(A) FISH and vGLUT1 immunostaining. Selected region corresponds to the neurite segment of mouse primary neurons in **Fig. 5A**. Arrows indicate additional examples of poly(A) (magenta) co-localizing with vGLUT1 signal (green). Scale bar: 20 µm.

**(B)** Representative full frame image of hippocampal neurons showing Atp5b smFISH signal (magenta) and vGLUT1 immunostaining (green). MAP2, a neuron specific marker is in blue and DAPI in cyan. Selected region corresponds to the neurite segment in **Fig. 5C**. Scale bar: 20 µm.

**(C)** Representative images of neurite segment selected in **(B)** indicating detected *Atp5b* foci (using radial symmetry-FISH; white circles overlay) from **Fig. 5C** and corresponding raw smFISH signal of *Atp5b* in magenta (image below). vGLUT1 is indicated in green (last image of panel C).

**(D)** Schematic representation of *Atp5b* transcript (*Mus musculus*, NM_016774.3) and 30 selected 18-20mer sequences used to design smFISH probes (light green) spanning over the *Atp5b* CDS and 3'UTR.

**(E)** Validation of anti-EIF4E antibody (abcam; ab33766) in not treated and 30 min Arsenite (330 µM) treated HEK293 cells. As expected EIF4E re-localize in stress granules also positive for poly(A) enrichment (magenta). DAPI is in cyan. Scale bar: 10 µm and 3 µm in selected inset. Arrows in selected inset indicate EIF4E (yellow; immunostained with anti-EIF4E) and poly(A) (magenta; FISH signal) positive stress granule.

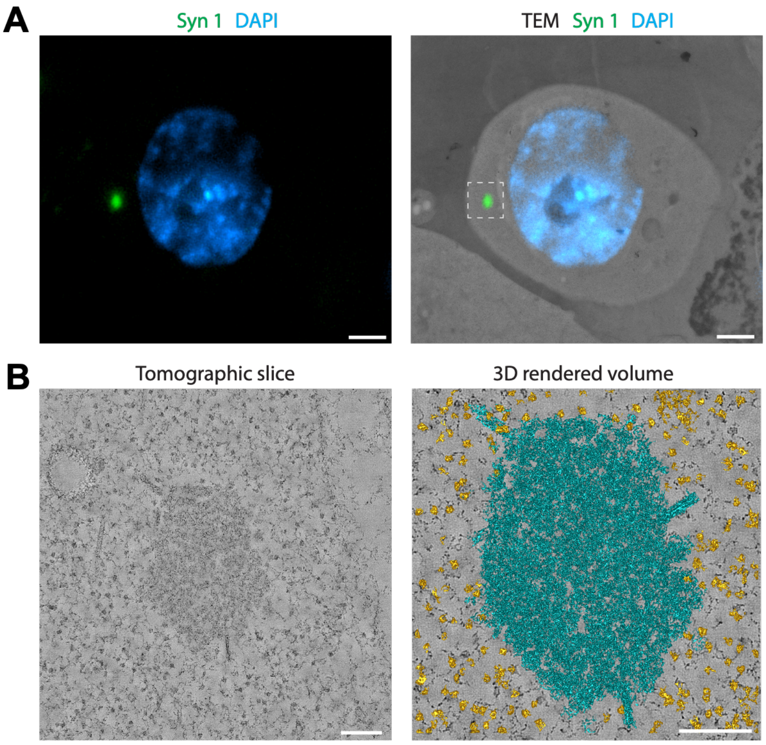

**Supplementary Figure 13 | Ribosomes localize in the vicinity of synapsin-1 condensates.**
**(A)** Fluorescence microscopy image of a 200 nm section of cryo-fixed HEK293 cells expressing EGFP-synapsin-1 and synaptophysin, overlaid with a TEM image of the same region. Scale bar, 2 µm.
**(B)** Tomographic reconstruction and 3D-rendered volume of the region indicated by the dashed box in *A*. Vesicular condensates are shown in cyan and ribosomes in orange. Scale bar, 200 nm.

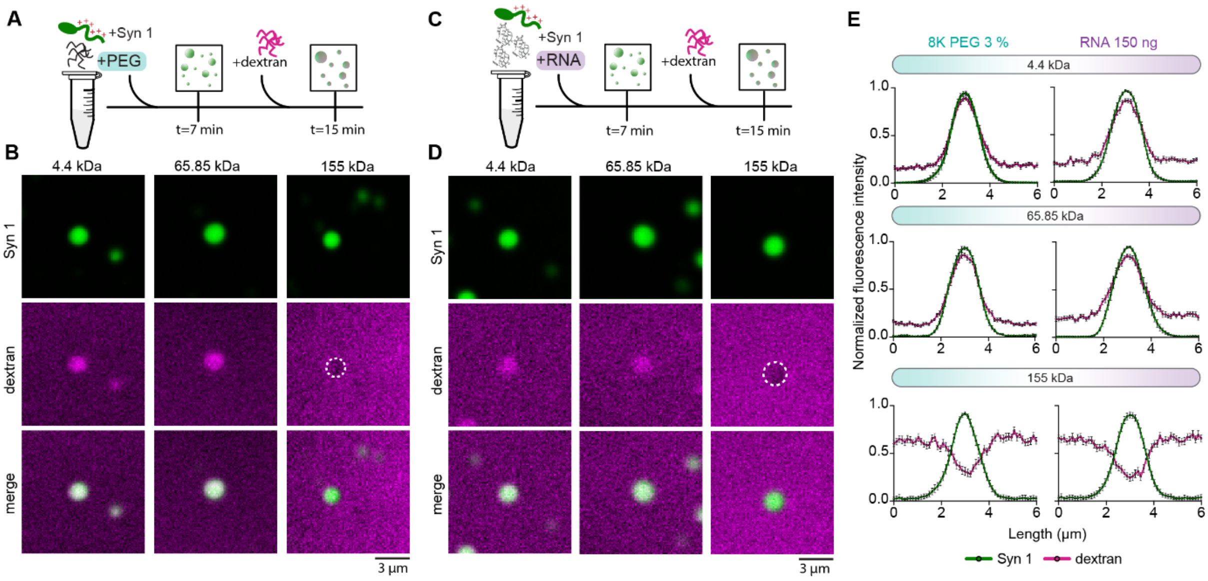

**Supplementary Figure 14 | RNA driven synapsin-1 condensates are permissive for micro to mesoscale molecules.**

**(A)** Representative scheme of *in vitro* reconstitution with Syn 1 (green), PEG 8000 and TMR-dextran (magenta) of specific molecular weight.

**(B)** Representative images of dextran assay for assessing the pore size of 4.5 µM Syn 1 condensates in the presence of 3 % molecular crowder PEG 8000 in the reaction buffer (150 mM NaCl, 25 mM Tris-HCl pH 7.4 and 0.5 mM TCEP).

**(C)** Representative scheme of *in vitro* reconstitution with Syn 1 (green), total neuronal RNA and TMR-dextran (magenta) of specific molecular weight.

**(D)** Representative images of dextran assay for assessing the pore size of 4.5 µM Syn 1 condensates in the presence of 150 ng of total neuronal RNA in reaction buffer (150 mM NaCl, 25 mM Tris-HCl pH 7.4 and 0.5 mM TCEP).

**(E)** Quantification of TMR dextran (sizes: 4.4, 65.85 and 155 kDa) enrichment in PEG- (left-hand panel plots) or RNA-driven (right-hand panel plots) Syn 1 condensates. Each plot represents line profiles of normalized fluorescence intensity signals with ±SEM of EGFP-Syn 1 (green) and TMR-dextran of different sizes (magenta). Data were collected from 3 independent *in vitro* reconstitutions.

**Supplementary Table S1: Plasmids used in this study.**

| Table S1. Plasmids in use | | | |
|---|---|---|---|
| **Plasmid Name** | **Expression system** | **Source** | **Sequence** |
| **mCherry-hsSYNAPSIN 1** | *mammalian* | Hoffman, Sansevrino et al., 2021 | ATGGGCCATCACCATCACCATCACAGCGGCAGCGAGAACCTGTACTT<br>CCAGGGCGTGAGCAAGGGCGAGGAGGATAACATGGCCATCATCAAG<br>GAGTTCATGCGCTTCAAGGTGCACATGGAGGGCTCCGTGAACGGCCA<br>CGAGTTCGAGATCGAGGGCGAGGGCGAGGGCCGCCCCTACGAGGGC<br>ACCCAGACCGCCAAGCTGAAGGTGACCAAGGGTGGCCCCCTGCCCTT<br>CGCCTGGGACATCCTGTCCCCTCAGTTCATGTACGGCTCCAAGGCCT<br>ACGTGAAGCACCCCGCCGACATCCCCGACTACTTGAAGCTGTCCTTC<br>CCCGAGGGCTTCAAGTGGGAGCGCGTGATGAACTTCGAGGACGGCG<br>GCGTGGTGACCGTGACCCAGGACTCCTCCCTGCAGGACGGCGAGTTC<br>ATCTACAAGGTGAAGCTGCGCGGCACCAACTTCCCCTCCGACGGCCC<br>CGTAATGCAGAAGAAGACCATGGGCTGGGAGGCCTCCTCCGAGCGG<br>ATGTACCCCGAGGACGGCGCCCTGAAGGGCGAGATCAAGCAGAGGC<br>TGAAGCTGAAGGACGGCGGCCACTACGACGCTGAGGTCAAGACCAC<br>CTACAAGGCCAAGAAGCCCGTGCAGCTGCCCGGCGCCTACAACGTC<br>AACATCAAGTTGGACATCACCTCCCACAACGAGGACTACACCATCGT<br>GGAACAGTACGAACGCGCCGAGGGCCGCCACTCCACCGGCGGCATG<br>GACGAGCTGTACAAGTCCGGACTCGGCGGCCACCGCTGGATCCTGG<br>AGGTGCTGTTCCAGGGCCCCAGATCTATGAACTACCTGCGCCGCCGC<br>CTGAGCGACAGCAACTTCATGGCCAACCTGCCCAACGGCTACATGAC<br>CGACCTGCAGCGCCCCCAGCCCCCCCCCCCCCCCCCCGGCGCCCACA<br>GCCCCGGCGCCACCCCCGGCCCCGGCACCGCCACCGCCGAGCGCAG<br>CAGCGGCGTGGCCCCCGCCGCCAGCCCCGCCGCCCCCAGCCCCGGCA<br>GCAGCGGCGGCGGCGGCTTCTTCAGCAGCCTGAGCAACGCCGTGAA<br>GCAGACCACCGCCGCCGCCGCCGCCACCTTCAGCGAGCAGGTGGGC<br>GGCGGCAGCGGCGGCGCCGGCCGCGGCGGCGCCGCCAGCCGCGTGC<br>TGCTGGTGATCGACGAGCCCCACACCGACTGGGCCAAGTACTTCAAG<br>GGCAAGAAGATCCACGGCGAGATCGACATCAAGGTGGAGCAGGCCG<br>AGTTCAGCGACCTGAACCTGGTGGCCCACGCCAACGGCGGCTTCAGC<br>GTGGACATGGAGGTGCTGCGCAACGGCGTGAAGGTGGTGCGCAGCC<br>TGAAGCCCGACTTCGTGCTGATCCGCCAGCACGCCTTCAGCATGGCC<br>CGCAACGGCGACTACCGCAGCCTGGTGATCGGCCTGCAGTACGCCG<br>GCATCCCCAGCGTGAACAGCCTGCACAGCGTGTACAACTTCTGCGAC<br>AAGCCCTGGGTGTTCGCCCAGATGGTGCGCCTGCACAAGAAGCTGG<br>GCACCGAGGAGTTCCCCCTGATCGACCAGACCTTCTACCCCAACCAC<br>AAGGAGATGCTGAGCAGCACCACCTACCCCGTGGTGGTGAAGATGG<br>GCCACGCCCACAGCGGCATGGGCAAGGTGAAGGTGGACAACCAGCA<br>CGACTTCCAGGACATCGCCAGCGTGGTGGCCCTGACCAAGACCTACG<br>CCACCGCCGAGCCCTTCATCGACGCCAAGTACGACGTGCGCGTGCAG<br>AAGATCGGCCAGAACTACAAGGCCTACATGCGCACCAGCGTGAGCG<br>GCAACTGGAAGACCAACACCGGCAGCGCCATGCTGGAGCAGATCGC<br>CATGAGCGACCGCTACAAGCTGTGGGTGGACACCTGCAGCGAGATA<br>TTCGGCGGCCTGGACATCTGCGCCGTGGAGGCCCTGCACGGCAAGG<br>ACGGCCGCGACCACATCATCGAGGTGGTGGGCAGCAGCATGCCCCT<br>GATCGGCGACCACCAGGACGAGGACAAGCAGCTGATCGTGGAGCTG |

| | | | |
|---|---|---|---|
| | | | GTGGTGAACAAGATGGCCCAGGCCCTGCCCCGCCAGCGCCAGCGCG<br>ACGCCAGCCCCGGCCGCGGCAGCCACGGCCAGACCCCCAGCCCCGG<br>CGCCCTGCCCCTGGGCCGCCAGACCAGCCAGCAGCCCGCCGGCCCCC<br>CCGCCCAGCAGCGCCCCCCCCCCAGGGCGGCCCCCCCAGCCCGGC<br>CCCGGCCCCCAGCGCCAGGGCCCCCCCTGCAGCAGCGCCCCCCCC<br>CCAGGGCCAGCAGCACCTGAGCGGCCTGGGCCCCCCCGCCGGCAGC<br>CCCCTGCCCCAGCGCCTGCCCAGCCCCACCAGCGCCCCCCAGCAGCC<br>CGCCAGCCAGGCCGCCCCCCCCACCCAGGGCCAGGGCCGCCAGAGC<br>CGCCCCGTGGCCGGCGGCCCCGGCGCCCCCCCCGCCGCCCGCCCCCC<br>CGCCAGCCCCAGCCCCCAGCGCCAGGCCGGCCCCCCCCAGGCCACCC<br>GCCAGACCAGCGTGAGCGGCCCCGCCCCCCCCAAGGCCAGCGGCGC<br>CCCCCCCGGCGGCCAGCAGCGCCAGGGCCCCCCCCAGAAGCCCCCC<br>GGCCCCGCCGGCCCCACCCGCCAGGCCAGCCAGGCCGGCCCCGTGC<br>CCCGCACCGGCCCCCCCACCACCCAGCAGCCCCGCCCCAGCGGCCCC<br>GGCCCCGCCGGCCGCCCCAAGCCCCAGCTGGCCCAGAAGCCCAGCC<br>AGGACGTGCCCCCCCCCGCCACCGCCGCCGCCGGCGGCCCCCCCCAC<br>CCCCAGCTGAACAAGAGCCAGAGCCTGACCAACGCCTTCAACCTGCC<br>CGAGCCCGCCCCCCCCCGCCCCAGCCTGAGCCAGGACGAGGTGAAG<br>GCCGAGACCATCCGCAGCCTGCGCAAGAGCTTCGCCAGCCTGTTCAG<br>CGACTGA |
| **hsSNCA** | *mammalian* | this study | ATGGATGTATTCATGAAAGGACTTTCAAAGGCCAAGGAGGGAGTTG<br>TGGCTGCTGCTGAGAAAACCAAACAGGGTGTGGCAGAAGCAGCAGG<br>AAAGACAAAAGAGGGTGTTCTCTATGTAGGCTCCAAAACCAAGGAG<br>GGAGTGGTGCATGGTGTGGCAACAGTGGCTGAGAAGACCAAAGAGC<br>AAGTGACAAATGTTGGAGGAGCAGTGGTGACGGGTGTGACAGCAGT<br>AGCCCAGAAGACAGTGGAGGGAGCAGGGAGCATTGCAGCAGCCACT<br>GGCTTTGTCAAAAAGGACCAGTTGGGCAAGAATGAAGAAGGAGCCC<br>CACAGGAAGGAATTCTGGAAGATATGCCTGTGGATCCTGACAATGA<br>GGCTTATGAAATGCCTTCTGAGGAAGGGTATCAAGACTACGAACCTG<br>AAGCCTAA |
| **hsSNCA-BFP** | *mammalian* | Hoffman, Sansevrino et al., 2021 | ATGAGCGAGCTGATTAAGGAGAACATGCACATGAAGCTGTACATGG<br>AGGGCACCGTGGACAACCATCACTTCAAGTGCACATCCGAGGGCGA<br>AGGCAAGCCCTACGAGGGCACCCAGACCATGAGAATCAAGGTGGTC<br>GAGGGCGGCCCTCTCCCCTTCGCCTTCGACATCCTGGCTACTAGCTTC<br>CTCTACGGCAGCAAGACCTTCATCAACCACACCCAGGGCATCCCCGA<br>CTTCTTCAAGCAGTCCTTCCCTGAGGGCTTCACATGGGAGAGAGTCA<br>CCACATACGAAGACGGGGGCGTGCTGACCGCTACCCAGGACACCAG<br>CCTCCAGGACGGCTGCCTCATCTACAACGTCAAGATCAGAGGGGTGA<br>ACTTCACATCCAACGGCCCTGTGATGCAGAAGAAAACACTCGGCTGG<br>GAGGCCTTCACCGAGACGCTGTACCCCGCTGACGGCGGCCTGGAAG<br>GCAGAAACGACATGGCCCTGAAGCTCGTGGGCGGGAGCCATCTGAT<br>CGCAAACATCAAGACCACATATAGATCCAAGAAACCCGCTAAGAAC<br>CTCAAGATGCCTGGCGTCTACTATGTGGACTACAGACTGGAAAGAAT<br>CAAGGAGGCCAACAACGAGACCTACGTCGAGCAGCACGAGGTGGCA<br>GTGGCCAGATACTGCGACCTCCCTAGCAAACTGGGGCACAAGCTTAA<br>TTCCGGACTCGGCGGCCACCGCTGGATCCTGGAGGTGCTGTTCCAGG<br>GCCCCAGATCTCGAGCTCAAGCTTCGATGGATGTATTCATGAAAGGA<br>CTTTCAAAGGCCAAGGAGGGAGTTGTGGCTGCTGCTGAGAAAACCA<br>AACAGGGTGTGGCAGAAGCAGCAGGAAAGACAAAAGAGGGTGTTCT<br>CTATGTAGGCTCCAAAACCAAGGAGGGAGTGGTGCATGGTGTGGCA<br>ACAGTGGCTGAGAAGACCAAAGAGCAAGTGACAAATGTTGGAGGAG<br>CAGTGGTGACGGGTGTGACAGCAGTAGCCCAGAAGACAGTGGAGGG<br>AGCAGGGAGCATTGCAGCAGCCACTGGCTTTGTCAAAAAGGACCAG<br>TTGGGCAAGAATGAAGAAGGAGCCCCACAGGAAGGAATTCTGGAAG |

| | | | |
|---|---|---|---|
| | | | ATATGCCTGTGGATCCTGACAATGAGGCTTATGAAATGCCTTCTGAG<br>GAAGGGTATCAAGACTACGAACCTGAAGCCTAA |
| **mCherry-empty** | *mammalian* | kind gift of Pietro De Camilli | ATGGGCCATCACCATCACCATCACAGCGGCAGCGAGAACCTGTACTT<br>CCAGGGCGTGAGCAAGGGCGAGGAGGATAACATGGCCATCATCAAG<br>GAGTTCATGCGCTTCAAGGTGCACATGGAGGGCTCCGTGAACGGCCA<br>CGAGTTCGAGATCGAGGGCGAGGGCGAGGGCCGCCCCTACGAGGGC<br>ACCCAGACCGCCAAGCTGAAGGTGACCAAGGGTGGCCCCCTGCCCTT<br>CGCCTGGGACATCCTGTCCCCTCAGTTCATGTACGGCTCCAAGGCCT<br>ACGTGAAGCACCCCGCCGACATCCCCGACTACTTGAAGCTGTCCTTC<br>CCCGAGGGCTTCAAGTGGGAGCGCGTGATGAACTTCGAGGACGGCG<br>GCGTGGTGACCGTGACCCAGGACTCCTCCCTGCAGGACGGCGAGTTC<br>ATCTACAAGGTGAAGCTGCGCGGCACCAACTTCCCCTCCGACGGCCC<br>CGTAATGCAGAAGAAGACCATGGGCTGGGAGGCCTCCTCCGAGCGG<br>ATGTACCCCGAGGACGGCGCCCTGAAGGGCGAGATCAAGCAGAGGC<br>TGAAGCTGAAGGACGGCGGCCACTACGACGCTGAGGTCAAGACCAC<br>CTACAAGGCCAAGAAGCCCGTGCAGCTGCCCGGCGCCTACAACGTC<br>AACATCAAGTTGGACATCACCTCCCACAACGAGGACTACACCATCGT<br>GGAACAGTACGAACGCGCCGAGGGCCGCCACTCCACCGGCGGCATG<br>GACGAGCTGTACAAG |
| **SYP-FL** | *mammalian* | this study | ATGGACGTGGTGAATCAGCTGGTGGCTGGGGGTCAGTTCCGGGTGGT<br>CAAGGAGCCCCTTGGCTTCGTGAAGGTGCTGCAGTGGGTCTTTGCCA<br>TCTTCGCCTTTGCTACGTGCGGCAGCTACACCGGAGAGCTTCGGCTG<br>AGCGTGGAGTGTGCCAACAAGACGGAGAGTGCCCTCAACATCGAAG<br>TCGAATTTGAGTACCCATTCAGGCTGCACCAAGTGTACTTTGATGCA<br>CCCTCCTGCGTTAAAGGGGGCACTACCAAGATCTTCCTAGTTGGTGA<br>CTACTCCTCCTCGGCTGAATTCTTTGTCACCGTGGCTGTGTTTGCCTT<br>CCTCTACTCCATGGGGGCCCTGGCCACCTACATCTTCCTGCAGAACA<br>AGTACCGAGAGAACAACAAAGGGCCAATGATGGACTTCCTGGCCAC<br>AGCAGTGTTCGCTTTCATGTGGCTAGTTAGCTCATCCGCCTGGGCCA<br>AAGGCCTGTCCGATGTGAAGATGGCCACTGACCCAGAGAACATTATC<br>AAGGAGATGCCTATGTGCCGCCAGACAGGAAACACATGCAAGGAAC<br>TGAGGGACCCTGTGACTTCAGGACTCAACACCTCGGTGGTGTTTGGC<br>TTCCTGAACCTGGTGCTCTGGGTTGGCAACCTATGGTTCGTGTTCAAG<br>GAGACAGGCTGGGCCGCCCCATTCATGCGCGCACCTCCAGGCGCCCC<br>AGAAAAGCAACCAGCTCCTGGCGATGCCTACGGCGATGCGGGCTAT<br>GGGCAGGGCCCCGGAGGCTATGGGCCCCAGGACTCCTACGGGCCTC<br>AGGGTGGTTATCAACCCGATTACGGGCAGCCAGCCAGCGGCGGTGG<br>CGGTGGCTACGGGCCTCAGGGCGACTATGGGCAGCAAGGCTACGGC<br>CAACAGGGTGCGCCCACCTCCTTCTCCAATCAGATG |
| **SYP-FL-EGFP** | *mammalian* | kind gift of Pietro De Camilli | ATGGACGTGGTGAATCAGCTGGTGGCTGGGGGTCAGTTCCGGGTGGT<br>CAAGGAGCCCCTTGGCTTCGTGAAGGTGCTGCAGTGGGTCTTTGCCA<br>TCTTCGCCTTTGCTACGTGCGGCAGCTACACCGGAGAGCTTCGGCTG<br>AGCGTGGAGTGTGCCAACAAGACGGAGAGTGCCCTCAACATCGAAG<br>TCGAATTTGAGTACCCATTCAGGCTGCACCAAGTGTACTTTGATGCA<br>CCCTCCTGCGTTAAAGGGGGCACTACCAAGATCTTCCTAGTTGGTGA<br>CTACTCCTCCTCGGCTGAATTCTTTGTCACCGTGGCTGTGTTTGCCTT<br>CCTCTACTCCATGGGGGCCCTGGCCACCTACATCTTCCTGCAGAACA<br>AGTACCGAGAGAACAACAAAGGGCCAATGATGGACTTCCTGGCCAC<br>AGCAGTGTTCGCTTTCATGTGGCTAGTTAGCTCATCCGCCTGGGCCA<br>AAGGCCTGTCCGATGTGAAGATGGCCACTGACCCAGAGAACATTATC<br>AAGGAGATGCCTATGTGCCGCCAGACAGGAAACACATGCAAGGAAC<br>TGAGGGACCCTGTGACTTCAGGACTCAACACCTCGGTGGTGTTTGGC |

| | | | |
|---|---|---|---|
| | | | TTCCTGAACCTGGTGCTCTGGGTTGGCAACCTATGGTTCGTGTTCAAG<br>GAGACAGGCTGGGCCGCCCCATTCATGCGCGCACCTCCAGGCGCCCC<br>AGAAAAGCAACCAGCTCCTGGCGATGCCTACGGCGATGCGGGCTAT<br>GGGCAGGGCCCCGGAGGCTATGGGCCCCAGGACTCCTACGGGCCTC<br>AGGGTGGTTATCAACCCGATTACGGGCAGCCAGCCAGCGGCGGTGG<br>CGGTGGCTACGGGCCTCAGGGCGACTATGGGCAGCAAGGCTACGGC<br>CAACAGGGTGCGCCCACCTCCTTCTCCAATCAGATGTCGGATCCACC<br>GGTCGCCACCATGGTGAGCAAGGGCGAGGAGCTGTTCACCGGGGTG<br>GTGCCCATCCTGGTCGAGCTGGACGGCGACGTAAACGGCCACAAGTT<br>CAGCGTGTCCGGCGAGGGCGAGGGCGATGCCACCTACGGCAAGCTG<br>ACCCTGAAGTTCATCTGCACCACCGGCAAGCTGCCCGTGCCCTGGCC<br>CACCCTCGTGACCACCCTGACCTACGGCGTGCAGTGCTTCAGCCGCT<br>ACCCCGACCACATGAAGCAGCACGACTTCTTCAAGTCCGCCATGCCC<br>GAAGGCTACGTCCAGGAGCGCACCATCTTCTTCAAGGACGACGGCA<br>ACTACAAGACCCGCGCCGAGGTGAAGTTCGAGGGCGACACCCTGGT<br>GAACCGCATCGAGCTGAAGGGCATCGACTTCAAGGAGGACGGCAAC<br>ATCCTGGGGCACAAGCTGGAGTACAACTACAACAGCCACAACGTCT<br>ATATCATGGCCGACAAGCAGAAGAACGGCATCAAGGTGAACTTCAA<br>GATCCGCCACAACATCGAGGACGGCAGCGTGCAGCTCGCCGACCAC<br>TACCAGCAGAACACCCCCATCGGCGACGGCCCCGTGCTGCTGCCCGA<br>CAACCACTACCTGAGCACCCAGTCCGCCCTGAGCAAAGACCCCAAC<br>GAGAAGCGCGATCACATGGTCCTGCTGGAGTTCGTGACCGCCGCCGG<br>GATCACTCTCGGCATGGACGAGCTGTACAAGTAA |
| **mScarlet-mmSYNAPSIN 1** | *mammalian* | this study | ATGGTGAGCAAGGGCGAGGCAGTGATCAAGGAGTTCATGCGGTTCA<br>AGGTGCACATGGAGGGCTCCATGAACGGCCACGAGTTCGAGATCGA<br>GGGCGAGGGCGAGGGCCGCCCCTACGAGGGCACCCAGACCGCCAAG<br>CTGAAGGTGACCAAGGGTGGCCCCCTGCCCTTCTCCTGGGACATCCT<br>GTCCCCTCAGTTCATGTACGGCTCCAGGGCCTTCATCAAGCACCCCG<br>CCGACATCCCCGACTACTATAAGCAGTCCTTCCCCGAGGGCTTCAAG<br>TGGGAGCGCGTGATGAACTTCGAGGACGGCGGCGCCGTGACCGTGA<br>CCCAGGACACCTCCCTGGAGGACGGCACCCTGATCTACAAGGTGAA<br>GCTCCGCGGCACCAACTTCCCTCCTGACGGCCCCGTAATGCAGAAGA<br>AGACAATGGGCTGGGAAGCGTCCACCGAGCGGTTGTACCCCGAGGA<br>CGGCGTGCTGAAGGGCGACATTAAGATGGCCCTGCGCCTGAAGGAC<br>GGCGGCCGCTACCTGGCGGACTTCAAGACCACCTACAAGGCCAAGA<br>AGCCCGTGCAGATGCCCGGCGCCTACAACGTCGACCGCAAGTTGGA<br>CATCACCTCCCACAACGAGGACTACACCGTGGTGGAACAGTACGAA<br>CGCTCCGAGGGCCGCCACTCCACCGGCGGCATGGACGAGCTGTACA<br>AGTCCGGACTCAGATCTATGAACTACCTGCGGCGCCGCCTGTCGGAC<br>AGCAACTTCATGGCCAATCTGCCGAATGGGTACATGACAGACCTGCA<br>GCGCCCGCAACCGCCCCCGCCGCCTCCCTCGGCCGCCAGCCCTGGGG<br>CCACGCCCGGCTCCGCGACAGCCTCTGCCGAGAGGGCCTCCACAGCT<br>GCTCCAGTGGCTTCTCCAGCAGCCCCTAGTCCTGGGTCCTCGGGGGG<br>CGGCGGCTTCTTCTCGTCGCTGTCTAACGCGGTCAAGCAAACCACAG<br>CAGCCGCAGCCGCCACCTTCAGCGAGCAGGTGGGCGGTGGCTCTGG<br>GGGCGCAGGCCGCGGGGGCGCCGCCGCCAGGGTGCTGCTGGTCATC<br>GACGAACCGCACACCGACTGGGCAAAATACTTCAAAGGGAAGAAGA<br>TCCATGGAGAAATTGACATTAAAGTAGAGCAAGCTGAATTCTCTGAT<br>CTCAATCTTGTGGCTCATGCCAATGGTGGATTCTCTGTGGACATGGA<br>AGTTCTTCGGAATGGAGTCAAAGTTGTGAGGTCTCTGAAGCCAGACT<br>TTGTGCTGATCCGCCAGCATGCCTTCAGCATGGCACGTAATGGAGAC<br>TACCGAAGTTTGGTCATTGGGCTGCAGTATGCTGGAATCCCCAGTGT<br>AAACTCTTTGCATTCTGTCTACAACTTCTGTGACAAACCCTGGGTGTT<br>TGCCCAGATGGTTCGACTACACAAGAAGCTTGGAACAGAGGAATTC<br>CCTCTGATTGATCAGACTTTCTATCCTAATCACAAAGAGATGCTCAG |

| | | | |
|---|---|---|---|
| | | | CAGCACAACATACCCTGTGGTTGTGAAGATGGGCCACGCACATTCTG<br>GGATGGGCAAGGTCAAGGTAGACAACCAACATGACTTCCAGGATAT<br>TGCAAGTGTTGTGGCACTGACTAAGACATATGCCACTGCTGAGCCCT<br>TCATTGATGCTAAATATGATGTGCGTGTCCAGAAGATTGGGCAGAAC<br>TACAAGGCCTACATGAGGACATCAGTGTCGGGTAACTGGAAGACCA<br>ATACAGGTTCTGCTATGCTTGAGCAGATTGCCATGTCTGACAGGTAC<br>AAGTTGTGGGTAGACACGTGCTCAGAGATTTTTGGGGGACTTGACAT<br>CTGCGCAGTGGAAGCGCTGCATGGCAAGGACGGAAGGGATCACATT<br>ATTGAGGTGGTGGGCTCCTCCATGCCACTCATTGGTGATCACCAGGA<br>TGAAGACAAGCAGCTCATCGTGGAACTTGTGGTCAACAAGATGACTC<br>AGGCTCTGCCTCGGCAGCCGCAGCGGGATGCTTCCCCTGGCAGGGGC<br>TCCCACAGCCAGTCTTCATCCCCAGGAGCCCTGACCTTGGGCCGCCA<br>GACCTCCCAGCAGCCTGCAGGTCCTCCTGCTCAACAACGACCCCCAC<br>CCCAGGGAGGCCCTCCACAGCCAGGCCCAGGACCTCAGCGCCAGGG<br>ACCCCCGCTGCAGCAGCGCCCACCCCCACAAGGCCAGCAACATCTTT<br>CTGGCCTTGGACCGCCAGCTGGCAGCCCTCTGCCTCAGCGCCTACCA<br>AGTCCCACCGCAGCACCTCAGCAGTCTGCCTCTCAGGCCACACCAGT<br>GACCCAGGGTCAAGGCCGCCAGTCGCGGCCAGTGGCAGGAGGCCCT<br>GGAGCACCTCCAGCAGCGCGCCCACCAGCCTCCCCATCTCCACAGCG<br>TCAGGCGGGGGCCCCGCAGGCTACCCGTCAGGCATCTATCTCTGGTC<br>CAGCTCCAACGAAGGCCTCAGGAGCCCCACCCGGAGGGCAGCAGCG<br>CCAGGGCCCTCCCCAAAAACCCCCAGGCCCTGCTGGTCCCACTCGTC<br>AGGCCAGTCAGGCAGGTCCCGGACCTCGCACTGGGCCTCCCACCACA<br>CAGCAGCCCCGGCCCAGCGGCCCAGGTCCTGCTGGACGTCCCGCCAA<br>ACCACAGCTGGCCCAGAAACCCAGCCAGGATGTGCCACCACCCATC<br>ACCGCCGCTGCCGGGGGACCCCCGCACCCCCAGCTCAACAAATCCCA<br>GTCTCTGACCAATGCCTTCAACCTTCCAGAGCCAGCCCCTCCCAGGC<br>CCAGCCTTAGCCAGGACGAGGTGAAAGCTGAGACCATCCGCAGCCT<br>GAGGAAGTCTTTCGCCAGCCTCTTCTCCGACTGA |
| **pET28-hsSNCA** | *E. coli* | Wallace et al., 2024 | ATGGATGTATTCATGAAAGGACTTTCAAAGGCCAAGGAGGGAGTTG<br>TGGCTGCTGCTGAGAAAACCAAACAGGGTGTGGCAGAAGCAGCAGG<br>AAAGACAAAAGAGGGTGTTCTCTATGTAGGCTCCAAAACCAAGGAG<br>GGAGTGGTGCATGGTGTGGCAACAGTGGCTGAGAAGACCAAAGAGC<br>AAGTGACAAATGTTGGAGGAGCAGTGGTGACGGGTGTGACAGCAGT<br>AGCCCAGAAGACAGTGGAGGGAGCAGGGAGCATTGCAGCAGCCACT<br>GGCTTTGTCAAAAAGGACCAGTTGGGCAAGAATGAAGAAGGAGCCC<br>CACAGGAAGGAATTCTGGAAGATATGCCTGTGGATCCTGACAATGA<br>GGCTTATGAAATGCCTTCTGAGGAAGGGTATCAAGACTACGAACCTG<br>AAGCCTAA |
| **pJet1.2-cln3.657** | IVT_template | this study | TAATACGACTCACTATAGGGAGAGTCTGCATACCAAAGATCAGCCGC<br>TTGCATCACATGAGACCCTGCGGCACTCTCTTATTTTGCGTAGTTTCT<br>TGGCCAAACTTAGGTAACCCACACTAAGAGGACACTGCCTGCATAA<br>ATACTGAAATTCTCTGTTCTCATCATCCGAGCAGTTAATAACTCCTAA<br>CGGACCCCATTTGCATCTTATTCGCACATCTGCGAAAGACTCTAGGA<br>AACACCCACGCTCTAAAGCATCAATACTCCTATTGTCCGTAAATGGT<br>ACATCCGTGTGGGAAAGTCACGCGCTCCATCGCTGGTTCCAGACCAG<br>AACGATGGAAGCGCATCAATATGCTTGCCAGCATGCCAGTGTTGCAC<br>TCCGCAAGCATCTCAGTGATCTGGAGAACCAGAAGCGGGCGTCCGG<br>GACTGGCTTCCCGTACCTGCCGCAACTAGGTATCGGGATGTTCAACT<br>TGAGTTTGCGCCACTTCATGTCCTTCCAAACTCGCTTGCGACTGTCCT<br>AATCCACTCGCCTATCATGGCAAGACATCATGGGGCTTTTCAGCTCC<br>AAGATGGTTTTTACTTCTACCCCCTTTCACCTACATACGCTTACGGCG<br>GTAAGCCTGTAGTGATAGTTCTTACATAAAAACACCCGCACCCCGAG<br>CCCGCAGCGCAATTGCTGCGACAACCGCTCACATGGGCGGTACCATG<br>ACCCGGAGCTGACCAACTTCTAGACGCTTCCCTGACCTGCGTATTCA |

| | | | |
|---|---|---|---|
| | | | GCTCCGAGTCATGTTTGCGCGCTTCATAACCTACAGATAACCGAGAC<br>CCCCAAGCCGTGCTCCATGCTCCTTAATCGAAACAGTTGTGAGCATG<br>GAGCAGGCATAATTGGGTCAGTATGCAGCTCTTACCCAGGAGTTTAC<br>TATCATAACTGCTCCAACGCCATTCATATGGCGAGCGTTACCCCTAT<br>AACAATCGCATCCCGATATGCTGACCCCAAAGGGTCAGCATGCTGCA<br>GGCCGTGTTTAACAGATGGAAATCTCATGGCGGTCTGCAGCCCCCAT<br>CTTTGCCTACAGGCCTGCGATGTCAAATTCCGGGCCTGTCTTTCCCTC<br>TAGTCGGGATCCTGGAGCAGCGTGCGAATCTAAAGCACGACTATGTC<br>TCCTCGGTAACCTTAATACCGCCAGTTAGCCGATGAGATCAATCTGT<br>AGTATGGTTATGCAAGCGCTTCCGTGGGCTCGCCAAGTGCGGGTCTA<br>GCTCCGGCCACTGCGGCCAGCGGCTCTCCCGCTGTCCGCATGGCAGG<br>CAGCTAGACCCCGGCCTATGTCCCGACATCCGCTCTGGTGACCCTTA<br>CGCGGCATATGCCGACGATCAAGGCTTTTATGCCGCTCACTCCAACT<br>ACTTCTCGCAAGCCAAAGCGCCTAGTGCTCTGGCTTAGTGAAAAGTA<br>CCAGACGAGGCCCGCGCCTGTGGAGGTTACATCATAGGCGCGCGCA<br>AAAGGCACTACCCATCAACATTCACCGCCTCTCCAGGAGATCGCGCA<br>GTTGCTGCATCAGTAGCAACATCAGCGCGACCCCATGGCCGTGGACC<br>TCCAGTTCAATGCCCGGGAACCACTCCTGGTCAAGAAACTACGCTAG<br>CATAC |
| **pJet1.2-cln3.362** | IVT_template | this study | TAATACGACTCACTATAGGGAGAGTCTGCATACCAAAGATCAGCCGC<br>TTGCATTGAAAGGGACGAACGGTGGCTCATGTCTTTGCCATCCTTTCT<br>TCTCCAATCCCCGAAGCACTTCAGACACACACGTAACCTTGCATGAA<br>ATATAGTATTTATTGTCATCGCCGGCTGGACTGTTACTATCTCAAGCC<br>CAAAGACCATTGCATATCACACACAAATTCGTCACAAACACTTAAAG<br>ATTGTCCTGCGGTCAGGCTTCTCTGCTACACCGGGCCTCTGTCAGTAT<br>GTCTGTTTGGGAACGTAAGTGTTCGCATCACCGGCTAATAAAAGTGG<br>ACGTGTTTGCGTAACAAACTGACTGCTCGGAGAACCGCATTTTAACC<br>CTTACGCTTCTGATTGATCTCGTGTATCACACCCGGTAATCATGTACT<br>TGTTTCCAGTCCCGACCCTTAGTTACCGTGGCGATTCGCAAATTGATT<br>GATGATTCTTGCCCTAGTTTCCCGCCATCCTTGTGCTTCCCCTCTTCG<br>ACGCTCTTTATAGGCAGCGCCAAATTTTATCGGTAGCGCGCTATGTT<br>CATTGTTATGTCACCCACATATCTTCTTCTTGGGCTTGACACTATTTG<br>GCTGACGTCCAAGTTCGCAGATCTAATGCCCCGACTATCGAGCTTCC<br>AAAGCCTCCGCCGCCACGAGCAAAAGAAACCGCATTTCACGACATC<br>AGAGCCGACAATCCCCAACACCCTTCACTGGTCAGTATGTACTACGG<br>CGACTCTCTATTCATAGGCGGAGATCAACCGGCAAACAAAGATAGG<br>CAGTCTCCAGTTCTGGCCCATCTCTATGAGCAGTTTAAGGTCTCGAG<br>CACCGAATCTTTGCAAGATATTCCCGTATGACCCAGACCCGAAATAC<br>AATACCAAGTTCAGCGCCGTTCACTTCGCAAGCGTAAGAGTTCTAAA<br>GACCGTACGCAGGCTCGCTGTGCTCTCCTAGGTCATGGTTTGCTGAC<br>GATGTACAACTGACAAAAATCCCATTCCCATCCGCACCCAGGTTCTA<br>GAGGTAGTCGCCAACCCGTCCAACGCAAACTCCAGATCTCCACCCAA<br>TTGCCTTTCAGGTGGTGCCTCGATCCTTGCCGTCTAAATTATAGCCAG<br>CCTGTGCCCATTTAACCGACGGTTAAGTAATGAGAAAAAGCGTTCCT<br>GGCGAGATGCAGGCACGGGGGTGGGAATCGAATGGCCAGGCGCCGC<br>AGTGCTCCGTGCTACCTGCCTCTCCCCCGATATCAGCAGTGACGGCG<br>ACTCTAGCTCGGGCTCTGTTCCACCATTTACTCTGGACCGGCCCGCCC<br>CGCACAAGACGACACTCAAGACAGCTGTGGCCCCCCTAACATCTACT<br>CCGAACGACTCACCGCGCACTCATCTCACGTATCGTTCGATGCCCCA<br>GACGCCACCCACTATTATGTCCGTTGCAGCTCAGGCGCGTGCACAAT<br>GCCTCCCAGACCTACACTCCACCACGACTAAGAACCCCCTTCACTTG<br>CTGCATTCTGAAGAATCTAAGGACTCCGCCACGGGTATCGACCATGA<br>GCTATGCGACATTGTTCAGGACTCGGTCAAGAAACTACGCTAGCATA<br>C |

| | | | |
|---|---|---|---|
| **pJet1.2-atp5b** | IVT_template | this study | TAATACGACTCACTATAGGTCTCCACCCGGATTCCGCCATGTTGAGT<br>CTTGTGGGGCGTGTGGCCTCGGCCTCGGCCTCCGGGGCCTTGCGGGG<br>ACTCAGCCCTTCGGCGGCTCTGCCACAGGCGCAGCTTCTACTGCGAG<br>CTGCTCCCGCCGGGGTTCATCCTGCCAGAGACTATGCGGCGCAGGCG<br>TCTGCGGCCCCGAAGGCAGGCACTGCCACCGGGCGAATCGTGGCAG<br>TCATCGGCGCTGTGGTGGACGTCCAGTTCGATGAGGGATTACCACCC<br>ATCCTAAATGCCCTGGAAGTGCAAGGCAGGGACAGCAGACTGGTTTT<br>GGAGGTGGCCCAGCATTTGGGGGAGAGCACGGTCAGAACTATTGCT<br>ATGGATGGCACTGAAGGCTTGGTTAGAGGCCAGAAAGTACTGGATT<br>CAGGGGCACCAATCAAAATTCCTGTTGGTCCTGAGACCTTGGGCAGA<br>ATCATGAATGTCATTGGAGAACCTATTGATGAGAGAGGTCCTATCAA<br>AACCAAACAATTTGCTCCTATTCATGCTGAGGCTCCTGAGTTCATAG<br>AGATGAGTGTTGAGCAGGAGATTCTGGTGACTGGGATAAAGGTTGT<br>GGATCTGCTGGCCCCATACGCCAAGGGTGGGAAAATCGGACTCTTTG<br>GAGGTGCTGGCGTTGGAAAGACAGTACTGATCATGGAGCTAATCAA<br>CAATGTCGCCAAAGCCCATGGTGGTTACTCTGTATTTGCTGGTGTTG<br>GTGAGAGGACCCGTGAGGGCAATGATTTATACCATGAAATGATTGA<br>ATCTGGTGTTATCAATCTAAAAGATGCCACTTCCAAGGTAGCGTTGG<br>TATATGGACAGATGAACGAACCACCTGGCGCTCGAGCCCGGGTAGC<br>TCTGACTGGTTTGACCGTTGCTGAATACTTCAGAGACCAGGAGGGCC<br>AAGATGTCCTGCTGTTTATTGACAACATCTTCCGCTTTACCCAGGCTG<br>GCTCAGAGGTGTCTGCCTTATTGGGCAGAATCCCTTCTGCTGTAGGC<br>TACCAGCCCACCCTAGCCACCGACATGGGCACAATGCAGGAAAGGA<br>TCACCACCACCAAGAAGGGATCGATCACCTCGGTGCAGGCTATCTAT<br>GTGCCTGCTGATGACCTGACTGACCCTGCCCCTGCAACCACCTTTGC<br>CCATTTGGATGCTACCACTGTGTTGTCCCGGGCTATTGCTGAGTTGGG<br>CATCTATCCAGCTGTGGATCCACTGGACTCCACCTCTCGAATTATGG<br>ATCCCAACATTGTTGGCAATGAGCATTATGACGTCGCCCGAGGAGTG<br>CAGAAAATCCTGCAGGACTACAAATCTCTCCAGGACATCATTGCCAT<br>CTTGGGTATGGATGAACTTTCTGAGGAAGATAAATTGACTGTGTCCC<br>GGGCAAGAAAGATACAGCGCTTCTTGTCTCAGCCATTCCAAGTTGCT<br>GAGGTCTTCACGGGTCACATGGGGAAGCTGGTGCCCTTGAAGGAGA<br>CCATTAAAGGATTCCAGCAGATTTTAGCAGGTGAATATGACCATCTC<br>CCAGAACAAGCCTTCTACATGGTGGGACCCATTGAAGAAGCCGTGG<br>CAAAGGCTGATAAGCTGGCAGAAGAGCACGGGTCGTGAGGGACTCC<br>AGCCAAAGGCAGCACTGCAACTGATCTCTCCATATCAAGCGAGAGCT<br>CAGGTTTCCTTCCATGCAGGCCACACAAGAGCCTTGATTGAAGATGT<br>GATGTTCTCTCTGAAGAGTATTTAAAGTTTTCAATAAAGTATATACCC<br>CTCATTTATGTCTGTTTATGTTGCTTCTGAAACAGCTTATAATTGAGC<br>TCACAGTGGTGGGCCACATTTAGGAAATTGTTAGAAACAGGATTATT<br>ATTAAAAAGATCAATTTATTAATG |
| **pJet1.2-scg3** | IVT_template | this study | TAATACGACTCACTATAGGGAGCATGCGCCTCCTCCTCATTTCCTCTC<br>CTGGAGCTAGTGGAGTAAAGCTACGCCCAGGCCCGCGTCCGCTGGC<br>GGCGCAGGAACTTCAGCACCCGCGGGGCGGACAGCGCCTACCGCAC<br>CTGCTCACCTGCTCTGGGCGCCAGAAGAGCCTGCATCCTCCTTCCAG<br>CCCGGAGCAACTGCGCCGGAGGCGCCCAGACCCTCTCCCTTCCCGCA<br>CCCAGGCTCCTGTCCCTTCCAGCTTCTTAACTCCCCTTCTCATTCATA<br>ACAAAAGCTACAGCTCAGGGGCCCAGCGCCAAGCTCTTTCCAGCAA<br>AGCACAGAAGAGCAAGAAAGAATGGGGTTCCTTTGGACCGGCTCTT<br>GGATACTGGTGTTGGTGCTCAACAGCGGCCCAATTCAAGCTTTCCCC<br>AAACCCGAAGGCAGCCAAGACAAATCCCTGCATAATAGAGAATTAA<br>GTGCAGAAAGACCTTTGAATGAACAGATCGCTGAGGCAGAGGCAGA<br>CAAGATTAAAAAGGCATTCCCTTCAGAAAGCAAGCCGAGTGAAAGC<br>AATTATTCTTCTGTCGATAACTTGAATCTGCTGAGGGCAATAACAGA<br>AAAGGAAACCGTTGAGAAAGAGAGACAATCCATAAGAAGCCCCCCG |

| | | | |
|---|---|---|---|
| | | | TTTGATAACCAACTGAACGTGGAAGACGCTGATTCAACCAAAAATCG<br>GAAACTGATCGATGAGTACGATTCCACCAAGAGTGGACTGGACCAC<br>AAGTTTCAAGATGACCCAGACGGCCTTCATCAACTGGATGGAACTCC<br>TTTAACTGCTGAAGACATCGTCCATAAGATTGCCACCAGGATTTATG<br>AGGAGAACGACAGAGGAGTGTTTGACAAAATTGTTTCTAAACTGCTG<br>AATCTTGGCCTGATCACTGAAAGCCAGGCACATACTCTGGAAGATGA<br>AGTAGCAGAAGCTTTACAAAAACTGATTTCAAAAGAGGCCAACAAT<br>TATGAGGAGACCCTGGATAAACCCACAAGCAGGACCGAGAATCAGG<br>ATGGGAAAATACCAGAGAAAGTGACTCCGGTGGCAGCAGTCCAAGA<br>TGGCTTCACTAACCGTGAAAACGATGAGACGGTGTCTAACACCTTGA<br>CCTTGTCCAATGGCTTGGAAAGGAGAACTAACCCCCACAGGGAAGA<br>CGACTTTGAGGAACTCCAGTATTTCCCCAACTTCTATGCACTACTGAC<br>AAGCATCGACTCAGAAAAAGAAGCAAAAGAGAAAGAAACCCTGATC<br>ACCATCATGAAGACATTGATTGACTTCGTGAAAATGATGGTGAAATA<br>CGGTACGATATCTCCAGAGGAAGGCGTGTCCTACCTTGAAAACTTGG<br>ATGAAACAATTGCTCTGCAGACCAAGAACAAGCTAGAAAAAAATAC<br>TACTGATAGCAAAACTCCACCAGAGAAGAGTCAGGAAGAAACAGAC<br>AGTACCAAGGAAGAAGCCGCCAAGATGGAAAAGGAATACGGAAGC<br>CTAAAAGACTCTACAAAAGATGATAACTCCAACCTAGGAGGAAAGA<br>CAGATGAAGCCACAGGGAAGACAGAAGCCTACTTGGAAGCCATTAG<br>AAAAAACATCGAATGGCTGAAGAAACATAACAAGAAGGGCAACAA<br>AGAAGATTACGACCTTTCAAAGATGAGGGACTTTATCAACCAACAA<br>GCTGACGCTTATGTGGAGAAGGGCATCCTCGACAAGGAAGAAGCCA<br>ACGCCATCAAACGCATCTACAGCAGCCTGTGAAAATGGCGGGCAGC<br>TTGAGCCTTCCTGTTGTTCCAGCAAAAACAATATAGCTTACAAACTA<br>ATTCGGCGGTTAAAGGGTTACCAGCCCAGAAGTATTAGGATGTGCTG<br>AATTTATAGTAGTTAATCCCTTAGAAATGAGTAAAATAGAGCTCTCT<br>TGCCATAAATACCTTATGAAAAGCAAAGCTGTAGAGAAGCCGAGGT<br>TTTTCTATATAGAATCCTTATTTCCTCTTGAATTTACATTTTGTAATCA<br>GAGATGTGCTGCTCTGGAAAAGACTCTAATGGGTTGAACATAAGTCT<br>GAACCTACTCCCCACTGTCCTCAGCCCCCTGAAGCTCTGAGAGGCCC<br>TGTCTCGGCATGCTAGACACCTGAGCACCTCACTGGATGTTTGTCAT<br>AGGATGTCGTTTCCACTAGTCGATCTCTGTTGGGCACGGAAATAAAC<br>CCACGTCTCTTCATATCCTTGA |
| **pJet1.2-prnp** | IVT_template | this study | TAATACGACTCACTATAGCCCCTTTCCACTCCCGGCTCCCCCGCGTTG<br>TCGGATCAGCAGACCGATTCTGGGCGCTGCGTCGCATCGGTGGCAGG<br>ACTCCTGAGTATATTTCAGAACTGAACCATTTCAACCGAGCTGAAGC<br>ATTCTGCCTTCCTAGTGGTACCAGTCCAATTTAGGAGAGCCAAGCAG<br>ACTATCAGTCATCATGGCGAACCTTGGCTACTGGCTGCTGGCCCTCTT<br>TGTGACTATGTGGACTGATGTCGGCCTCTGCAAAAAGCGGCCAAAGC<br>CTGGAGGGTGGAACACCGGTGGAAGCCGGTATCCCGGGCAGGGAAG<br>CCCTGGAGGCAACCGTTACCCACCTCAGGGTGGCACCTGGGGGCAG<br>CCCCACGGTGGTGGCTGGGGACAACCCCATGGGGGCAGCTGGGGAC<br>AACCTCATGGTGGTAGTTGGGGTCAGCCCCATGGCGGTGGATGGGGC<br>CAAGGAGGGGGTACCCATAATCAGTGGAACAAGCCCAGCAAACCAA<br>AAACCAACCTCAAGCATGTGGCAGGGGCTGCGGCAGCTGGGGCAGT<br>AGTGGGGGGCCTTGGTGGCTACATGCTGGGGAGCGCCATGAGCAGG<br>CCCATGATCCATTTTGGCAACGACTGGGAGGACCGCTACTACCGTGA<br>AAACATGTACCGCTACCCTAACCAAGTGTACTACAGGCCAGTGGATC<br>AGTACAGCAACCAGAACAACTTCGTGCACGACTGCGTCAATATCACC<br>ATCAAGCAGCACACGGTCACCACCACCACCAAGGGGGAGAACTTCA<br>CCGAGACCGATGTGAAGATGATGGAGCGCGTGGTGGAGCAGATGTG<br>CGTCACCCAGTACCAGAAGGAGTCCCAGGCCTATTACGACGGGAGA<br>AGATCCAGCAGCACCGTGCTTTTCTCCTCCCCTCCTGTCATCCTCCTC<br>ATCTCCTTCCTCATCTTCCTGATCGTGGGATGAGGGAGGCCTTCCTGC |

| | | | |
|---|---|---|---|
| | | | TTGTTCCTTCGCATTCTCGTGGTCTAGGCTGGGGGAGGGGTTATCCAC<br>CTGTAGCTCTTTCAATTGAGGTGGTTCTCATTCTTGCTTCTCTGTGTCC<br>CCCATAGGCTAATACCCCTGGCACTGATGGGCCCTGGGAAATGTACA<br>GTAGACCAGTTGCTCTTTGCTTCAGGTCCCTTTGATGGAGTCTGTCAT<br>CAGCCAGTGCTAACACCGGGCCAATAAGAATATAACACCAAATAAC<br>TGCTGGCTAGTTGGGGCTTTGTTTTGGTCTAGTGAATAAATACTGGTG<br>TATCCCCTGACTTGTACCCAGAGTACAAGGTGACAGTGACACATGTA<br>ACTTAGCATAGGCAAAGGGTTCTACAACCAAAGAAGCCACTGTTTGG<br>GGATGGCGCCCTGGAAAACAGCCTCCCACCTGGGATAGCTAGAGCA<br>TCCACACGTGGAATTCTTTCTTTACTAACAAACGATAGCTGATTGAA<br>GGCAACAGGAAAAAAAAAATCAAATTGTCCTACTGACGTTGAAAGC<br>AAACCTTTGTTCATTCCCAGGGCACTAGAATGATCTTTAGCCTTGCTT<br>GGATTGAACTAGGAGATCTTGACTCTGAGGAGAGCCAGCCCTGTAA<br>AAAGCTTGGTCCTCCTGTGACGGGAGGGATGGTTAAGGTACAAAGG<br>CTAGAAACTTGAGTTTCTTCATTTCTGTCTCACAATTATCAAAAGCTA<br>GAATTAGCTTCTGCCCTATGTTTCTGTACTTCTATTTGAACTGGATAA<br>CAGAGAGACAATCTAAACATTCTCTTAGGCTGCAGATAAGAGAAGT<br>AGGCTCCATTCCAAAGTGGGAAAGAAATTCTGCTAGCATTGTTTAAA<br>TCAGGCAAAATTTGTTCCTGAAGTTGCTTTTTACCCCAGCAGACATA<br>AACTGCGATAGCTTCAGCTTGCACTGTGGATTTTCTGTATAGAATAT<br>ATAAAACATAACTTCAAGCTTATGTCTTCTTTTTAAAACATCTGAAGT<br>ATGGGACGCCCTGGCCGTTCCATCCAGTACTAAATGCTTACCGTGTG<br>ACCCTTGGGCTTTCAGCGTGCACTCAGTTCCGTAGGATTCCAAAGCA<br>GACCCCTAGCTGGTCTTTGAATCTGCATGTACTTCACGTTTTCTATAT<br>TTGTAACTTTGCATGTATTTTGTTTTGTCATATAAAAAGTTTATAAAT<br>GTTTGCTATCAGACTGACATTAAATAGAAGCTATGATGAACACCTGG |
| **pJet1.2-atp5b::nluc** | IVT_template | this study | TAATACGACTCACTATAAGGCCACCATGTTGAGTCTTGTGGGGCGTG<br>TGGCCTCGGCCTCGGCCTCCGGGGCCTTGCGGGGACTCAGCCCTTCG<br>GCGGCTCTGCCACAGGCGCAGCTTCTACTGCGAGCTGCTCCCGCCGG<br>GGTTCATCCTGCCAGAGACTATGCGGCGCAGGCGTCTGCGGCCCCGA<br>AGGCAGGCACTGCCACCGGGCGAATCGTGGCAGTCATCGGCGCTGT<br>GGTGGACGTCCAGTTCGATGAGGGATTACCACCCATCCTAAATGCCC<br>TGGAAGTGCAAGGCAGGGACAGCAGACTGGTTTTGGAGGTGGCCCA<br>GCATTTGGGGGAGAGCACGGTCAGAACTATTGCTATGGATGGCACTG<br>AAGGCTTGGTTAGAGGCCAGAAAGTACTGGATTCAGGGGCACCAAT<br>CAAAATTCCTGTTGGTCCTGAGACCTTGGGCAGAATCATGAATGTCA<br>TTGGAGAACCTATTGATGAGAGAGGTCCTATCAAAACCAAACAATTT<br>GCTCCTATTCATGCTGAGGCTCCTGAGTTCATAGAGATGAGTGTTGA<br>GCAGGAGATTCTGGTGACTGGGATAAAGGTTGTGGATCTGCTGGCCC<br>CATACGCCAAGGGTGGGAAAATCGGACTCTTTGGAGGTGCTGGCGTT<br>GGAAAGACAGTACTGATCATGGAGCTAATCAACAATGTCGCCAAAG<br>CCCATGGTGGTTACTCTGTATTTGCTGGTGTTGGTGAGAGGACCCGT<br>GAGGGCAATGATTTATACCATGAAATGATTGAATCTGGTGTTATCAA<br>TCTAAAAGATGCCACTTCCAAGGTAGCGTTGGTATATGGACAGATGA<br>ACGAACCACCTGGCGCTCGAGCCCGGGTAGCTCTGACTGGTTTGACC<br>GTTGCTGAATACTTCAGAGACCAGGAGGGCCAAGATGTCCTGCTGTT<br>TATTGACAACATCTTCCGCTTTACCCAGGCTGGCTCAGAGGTGTCTG<br>CCTTATTGGGCAGAATCCCTTCTGCTGTAGGCTACCAGCCCACCCTA<br>GCCACCGACATGGGCACAATGCAGGAAAGGATCACCACCACCAAGA<br>AGGGATCGATCACCTCGGTGCAGGCTATCTATGTGCCTGCTGATGAC<br>CTGACTGACCCTGCCCCTGCAACCACCTTTGCCCATTTGGATGCTACC<br>ACTGTGTTGTCCCGGGCTATTGCTGAGTTGGGCATCTATCCAGCTGTG<br>GATCCACTGGACTCCACCTCTCGAATTATGGATCCCAACATTGTTGG<br>CAATGAGCATTATGACGTCGCCCGAGGAGTGCAGAAAATCCTGCAG<br>GACTACAAATCTCTCCAGGACATCATTGCCATCTTGGGTATGGATGA |

| | | | |
|---|---|---|---|
| | | | ACTTTCTGAGGAAGATAAATTGACTGTGTCCCGGGCAAGAAAGATAC<br>AGCGCTTCTTGTCTCAGCCATTCCAAGTTGCTGAGGTCTTCACGGGTC<br>ACATGGGGAAGCTGGTGCCCTTGAAGGAGACCATTAAAGGATTCCA<br>GCAGATTTTAGCAGGTGAATATGACCATCTCCCAGAACAAGCCTTCT<br>ACATGGTGGGACCCATTGAAGAAGCCGTGGCAAAGGCTGATAAGCT<br>GGCAGAAGAGCACGGGTCGATGGTCTTCACACTCGAAGATTTCGTTG<br>GGGACTGGCGACAGACAGCCGGCTACAACCTGGACCAAGTCCTTGA<br>ACAGGGAGGTGTGTCCAGTTTGTTTCAGAATCTCGGGGTGTCCGTAA<br>CTCCGATCCAAAGGATTGTCCTGAGCGGTGAAAATGGGCTGAAGATC<br>GACATCCATGTCATCATCCCGTATGAAGGTCTGAGCGGCGACCAAAT<br>GGGCCAGATCGAAAAAATTTTTAAGGTGGTGTACCCTGTGGATGATC<br>ATCACTTTAAGGTGATCCTGCACTATGGCACACTGGTAATCGACGGG<br>GTTACGCCGAACATGATCGACTATTTCGGACGGCCGTATGAAGGCAT<br>CGCCGTGTTCGACGGCAAAAAGATCACTGTAACAGGGACCCTGTGG<br>AACGGCAACAAAATTATCGACGAGCGCCTGATCAACCCCGACGGCT<br>CCCTGCTGTTCCGAGTAACCATCAACGGAGTGACCGGCTGGCGGCTG<br>TGCGAACGCATTCTGGCGGGTGGGGGATCATTGTACAAATAA |
| **pJet1.2-atp5b::moon** | IVT_template | this study | TAATACGACTCACTATAAGGCCACCATGTTGAGTCTTGTGGGGCGTG<br>TGGCCTCGGCCTCGGCCTCCGGGGCCTTGCGGGGACTCAGCCCTTCG<br>GCGGCTCTGCCACAGGCGCAGCTTCTACTGCGAGCTGCTCCCGCCGG<br>GGTTCATCCTGCCAGAGACTATGCGGCGCAGGCGTCTGCGGCCCCGA<br>AGGCAGGCACTGCCACCGGGCGAATCGTGGCAGTCATCGGCGCTGT<br>GGTGGACGTCCAGTTCGATGAGGGATTACCACCCATCCTAAATGCCC<br>TGGAAGTGCAAGGCAGGGACAGCAGACTGGTTTTGGAGGTGGCCCA<br>GCATTTGGGGGAGAGCACGGTCAGAACTATTGCTATGGATGGCACTG<br>AAGGCTTGGTTAGAGGCCAGAAAGTACTGGATTCAGGGGCACCAAT<br>CAAAATTCCTGTTGGTCCTGAGACCTTGGGCAGAATCATGAATGTCA<br>TTGGAGAACCTATTGATGAGAGAGGTCCTATCAAAACCAAACAATTT<br>GCTCCTATTCATGCTGAGGCTCCTGAGTTCATAGAGATGAGTGTTGA<br>GCAGGAGATTCTGGTGACTGGGATAAAGGTTGTGGATCTGCTGGCCC<br>CATACGCCAAGGGTGGGAAAATCGGACTCTTTGGAGGTGCTGGCGTT<br>GGAAAGACAGTACTGATCATGGAGCTAATCAACAATGTCGCCAAAG<br>CCCATGGTGGTTACTCTGTATTTGCTGGTGTTGGTGAGAGGACCCGT<br>GAGGGCAATGATTTATACCATGAAATGATTGAATCTGGTGTTATCAA<br>TCTAAAAGATGCCACTTCCAAGGTAGCGTTGGTATATGGACAGATGA<br>ACGAACCACCTGGCGCTCGAGCCCGGGTAGCTCTGACTGGTTTGACC<br>GTTGCTGAATACTTCAGAGACCAGGAGGGCCAAGATGTCCTGCTGTT<br>TATTGACAACATCTTCCGCTTTACCCAGGCTGGCTCAGAGGTGTCTG<br>CCTTATTGGGCAGAATCCCTTCTGCTGTAGGCTACCAGCCCACCCTA<br>GCCACCGACATGGGCACAATGCAGGAAAGGATCACCACCACCAAGA<br>AGGGATCGATCACCTCGGTGCAGGCTATCTATGTGCCTGCTGATGAC<br>CTGACTGACCCTGCCCCTGCAACCACCTTTGCCCATTTGGATGCTACC<br>ACTGTGTTGTCCCGGGCTATTGCTGAGTTGGGCATCTATCCAGCTGTG<br>GATCCACTGGACTCCACCTCTCGAATTATGGATCCCAACATTGTTGG<br>CAATGAGCATTATGACGTCGCCCGAGGAGTGCAGAAAATCCTGCAG<br>GACTACAAATCTCTCCAGGACATCATTGCCATCTTGGGTATGGATGA<br>ACTTTCTGAGGAAGATAAATTGACTGTGTCCCGGGCAAGAAAGATAC<br>AGCGCTTCTTGTCTCAGCCATTCCAAGTTGCTGAGGTCTTCACGGGTC<br>ACATGGGGAAGCTGGTGCCCTTGAAGGAGACCATTAAAGGATTCCA<br>GCAGATTTTAGCAGGTGAATATGACCATCTCCCAGAACAAGCCTTCT<br>ACATGGTGGGACCCATTGAAGAAGCCGTGGCAAAGGCTGATAAGCT<br>GGCAGAAGAGCACGGGTCGATGCGTACGGGCAAGAATGAGCAGGA<br>ACTGCTGGAACTGGACAAATGGGCAAGCCTGGGGAGCGGCTCCGGA<br>AAGAACGAGCAGGAACTGCTGGAACTGGACAAGTGGGCTTCTCTGG<br>GCTCCGGGTCCGGGAAAAACGAACAGGAGCTGCTGGAGCTGGATAA |

| | | | |
|---|---|---|---|
| | | | GTGGGCCAGCCTGGGCTCCGGGTCCGGGAAGAATGAACAGGAACTG<br>CTGGAACTGGATAAATGGGCCTCTCTGGGCAGCGGGTCCGGAAAGA<br>ATGAACAGGAGCTGCTGGAACTGGACAAGTGGGCGTCCCTGGGGAG<br>CGGGTCCGGAAAGAATGAACAGGAACTGCTGGAACTGGATAAGTGG<br>GCTTCCCTGGGAAGCGGCTCCGGAAAAAACGAGCAGGAGCTGCTGG<br>AGCTGGACAAGTGGGCCTCTCTGGGCTCCGGCTCCGGGAAGAATGA<br>ACAGGAGCTGCTGGAACTGGATAAGTGGGCCTCTCTGGGGTCCGGA<br>AGCGGTAAAAACGAGCAGGAACTGCTGGAACTGGACAAGTGGGCAA<br>GCCTGGGGTCTGGCTCCGGCAAAAACGAACAGGAACTGCTGGAACT<br>GGACAAATGGGCCAGCCTGGGCTCCGGGTCCGGGAAAAATGAGCAG<br>GAACTGCTGGAACTGGATAAATGGGCGAGCCTGGGCAGCGGGTCCG<br>GAAAAAACGAACAGGAGCTGCTGGAGCTGGACAAATGGGCGAGTCT<br>GGGCTCCGGAAGCGGCAAGAATGAACAGGAACTGCTGGAGCTGGAC<br>AAATGGGCCAGCCTGGGCAGCGGCTCCGGAAAGAATGAGCAGGAAC<br>TGCTGGAGCTGGACAAATGGGCGTCTCTGGGCTCCGGCTCCGGGAAA<br>AACGAGCAGGAACTGCTGGAGCTGGACAAGTGGGCTTCCCTGGGCA<br>GCGGCTCCGGAAAAAATGAACAGGAACTGCTGGAACTGGACAAGTG<br>GGCCAGCCTGGGGTCCGGGTCTGGGAAGAACGAGCAGGAACTGCTG<br>GAGCTGGACAAGTGGGCTTCTCTGGGGTCTGGCTCTGGCAAGAACGA<br>GCAGGAACTGCTGGAACTGGACAAGTGGGCGAGCCTGGGCAGCGGG<br>TCTGGCAAGAATGAACAGGAGCTGCTGGAACTGGACAAATGGGCCA<br>GCCTGGGGTCTGGGTCTGGAAAGAATGAGCAGGAACTGCTGGAGCT<br>GGACAAGTGGGCCTCTCTGGGCTCTGGGTCTGGCAAGAACGAGCAG<br>GAGCTGCTGGAGCTGGATAAGTGGGCGTCCCTGGGGTCCGGGTCCG<br>GCAAGAATGAACAGGAGCTGCTGGAGCTGGATAAATGGGCTTCCCT<br>GGGCTCTGGGTCTGGGAAGAATGAGCAGGAGCTGCTGGAACTGGAT<br>AAGTGGGCGAGCCTGGGCAGCGGGTCTGGCAAGAATGAGCAGGAAC<br>TGCTGGAGCTGGACAAGTGGGCTTCTCTGTAA |
| **pET30b-2H10-mCherry** | *E. coli* | Geisterfer&Jalihal et al., 2024 | ATGCACCATCATCATCATCATTCTTCTGGTGAAAACCTGTATTTTCAG<br>GGCGCCGAGGTGCAGCTGGTGGAATCTGGGGGCGGACTGGTGCAGC<br>CCGGGGGATCTCTGCGGCTGTCCTGCGCCGCCTCTGGCTCCATCTCTA<br>GCGTGGATGTGATGTCCTGGTACAGGCAGGCCCCTGGCAAGCAGAG<br>GGAACTGGTGGCCTTCATTACTGATAGGGGAAGAACCAATTACAAA<br>GTGAGCGTGAAAGGCCGCTTCACTATCAGCCGGGATAATTCTAAGAA<br>TATGGTGTATCTGCAGATGAACAGCCTGAAGCCAGAAGACACTGCC<br>GACTATCTGTGCAGGGCTGAGTCCCGGACTTCCTGGTCCAGCCCATC<br>TCCTCTGGATGTGTGGGGGCGCGGCACTCAGGTGACTGTGTCCTCCG<br>AATTGGATCCAGGTGGAGGTGGAAGCGGTGTGAGCAAGGGCGAGGA<br>GGATAACATGGCCATCATCAAGGAGTTCATGCGCTTCAAGGTGCACA<br>TGGAGGGCTCCGTGAACGGCCACGAGTTCGAGATCGAGGGCGAGGG<br>CGAGGGCCGCCCCTACGAGGGCACCCAGACCGCCAAGCTGAAGGTG<br>ACCAAGGGTGGCCCCCTGCCCTTCGCCTGGGACATCCTGTCCCCTCA<br>GTTCATGTACGGCTCCAAGGCCTACGTGAAGCACCCCGCCGACATCC<br>CCGACTACTTGAAGCTGTCCTTCCCCGAGGGCTTCAAGTGGGAGCGC<br>GCGATGAACTTCGAGGACGGCGGCGTGGTGACCGTGACCCAGGACT<br>CCTCCCTGCAGGACGGCGAGTTCATCTACAAGGTGAAGCTGCGCGGC<br>ACCAACTTCCCCTCCGACGGCCCCGTAATGCAGAAGAAGACCATGG<br>GCTGGGAGGCCTCCTCCGAGCGGATGTACCCCGAGGACGGCGCCCT<br>GAAGGGCGAGATCAAGCAGAGGCTGAAGCTGAAGGACGGCGGCCA<br>CTACGACGCTGAGGTCAAGACCACCTACAAGGCCAAGAAGCCCGTG<br>CAGCTGCCCGGCGCCTACAACGTCAACATCAAGTTGGACATCACCTC<br>CCACAACGAGGACTACACCATCGTGGAACAGTACGAACGCGCCGAG<br>GGCCGCCACTCCACCGGCGGCATGGACGAGCTGTACAAGTAA |

**Supplementary Table S2: cDNA and Primers for in vitro transcription.**

| Table S2 cDNA and Primers for in vitro transcription | | | |
|---|---|---|---|
| **Plasmid** | **cDNA template** | **Forward primer** | **Reverse primer** |
| **pJet1.2-cln3.657** | TAATACGACTCACTATAGGGAGAGTCTGCATAC<br>CAAAGATCAGCCGCTTGCATCACATGAGACCCT<br>GCGGCACTCTCTTATTTTGCGTAGTTTCTTGGCC<br>AAACTTAGGTAACCCACACTAAGAGGACACTGC<br>CTGCATAAATACTGAAATTCTCTGTTCTCATCAT<br>CCGAGCAGTTAATAACTCCTAACGGACCCCATT<br>TGCATCTTATTCGCACATCTGCGAAAGACTCTAG<br>GAAACACCCACGCTCTAAAGCATCAATACTCCT<br>ATTGTCCGTAAATGGTACATCCGTGTGGGAAAG<br>TCACGCGCTCCATCGCTGGTTCCAGACCAGAAC<br>GATGGAAGCGCATCAATATGCTTGCCAGCATGC<br>CAGTGTTGCACTCCGCAAGCATCTCAGTGATCTG<br>GAGAACCAGAAGCGGGCGTCCGGGACTGGCTTC<br>CCGTACCTGCCGCAACTAGGTATCGGGATGTTC<br>AACTTGAGTTTGCGCCACTTCATGTCCTTCCAAA<br>CTCGCTTGCGACTGTCCTAATCCACTCGCCTATC<br>ATGGCAAGACATCATGGGGCTTTTCAGCTCCAA<br>GATGGTTTTTACTTCTACCCCCTTTCACCTACAT<br>ACGCTTACGGCGGTAAGCCTGTAGTGATAGTTC<br>TTACATAAAAACACCCGCACCCCGAGCCCGCAG<br>CGCAATTGCTGCGACAACCGCTCACATGGGCGG<br>TACCATGACCCGGAGCTGACCAACTTCTAGACG<br>CTTCCCTGACCTGCGTATTCAGCTCCGAGTCATG<br>TTTGCGCGCTTCATAACCTACAGATAACCGAGA<br>CCCCCAAGCCGTGCTCCATGCTCCTTAATCGAAA<br>CAGTTGTGAGCATGGAGCAGGCATAATTGGGTC<br>AGTATGCAGCTCTTACCCAGGAGTTTACTATCAT<br>AACTGCTCCAACGCCATTCATATGGCGAGCGTT<br>ACCCCTATAACAATCGCATCCCGATATGCTGAC<br>CCCAAAGGGTCAGCATGCTGCAGGCCGTGTTTA<br>ACAGATGGAAATCTCATGGCGGTCTGCAGCCCC<br>CATCTTTGCCTACAGGCCTGCGATGTCAAATTCC<br>GGGCCTGTCTTTCCCTCTAGTCGGGATCCTGGAG<br>CAGCGTGCGAATCTAAAGCACGACTATGTCTCC<br>TCGGTAACCTTAATACCGCCAGTTAGCCGATGA<br>GATCAATCTGTAGTATGGTTATGCAAGCGCTTCC<br>GTGGGCTCGCCAAGTGCGGGTCTAGCTCCGGCC<br>ACTGCGGCCAGCGGCTCTCCCGCTGTCCGCATG<br>GCAGGCAGCTAGACCCCGGCCTATGTCCCGACA<br>TCCGCTCTGGTGACCCTTACGCGGCATATGCCGA<br>CGATCAAGGCTTTTATGCCGCTCACTCCAACTAC<br>TTCTCGCAAGCCAAAGCGCCTAGTGCTCTGGCTT<br>AGTGAAAAGTACCAGACGAGGCCCGCGCCTGTG<br>GAGGTTACATCATAGGCGCGCAAAAGGCACT<br>ACCCATCAACATTCACCGCCTCTCCAGGAGATC<br>GCGCAGTTGCTGCATCAGTAGCAACATCAGCGC<br>GACCCCATGGCCGTGGACCTCCAGTTCAATGCC<br>CGGGAACCACTCCTGGTCAAGAAACTACGCTAG<br>CATAC | taatacgactc actataggg | gtatgctagcg tagtttctt |

| | | | |
|---|---|---|---|
| **pJet1.2-cln3.362** | TAATACGACTCACTATAGGGAGAGTCTGCATAC<br>CAAAGATCAGCCGCTTGCATTGAAAGGGACGAA<br>CGGTGGCTCATGTCTTTGCCATCCTTTCTTCTCC<br>AATCCCCGAAGCACTTCAGACACACACGTAACC<br>TTGCATGAAATATAGTATTTATTGTCATCGCCGG<br>CTGGACTGTTACTATCTCAAGCCCAAAGACCATT<br>GCATATCACACACAAATTCGTCACAAACACTTA<br>AAGATTGTCCTGCGGTCAGGCTTCTCTGCTACAC<br>CGGGCCTCTGTCAGTATGTCTGTTTGGGAACGTA<br>AGTGTTCGCATCACCGGCTAATAAAAGTGGACG<br>TGTTTGCGTAACAAACTGACTGCTCGGAGAACC<br>GCATTTTAACCCTTACGCTTCTGATTGATCTCGT<br>GTATCACACCCGGTAATCATGTACTTGTTTCCAG<br>TCCCGACCCTTAGTTACCGTGGCGATTCGCAAAT<br>TGATTGATGATTCTTGCCCTAGTTTCCCGCCATC<br>CTTGTGCTTCCCCTCTTCGACGCTCTTTATAGGC<br>AGCGCCAAATTTTATCGGTAGCGCGCTATGTTCA<br>TTGTTATGTCACCCACATATCTTCTTCTTGGGCTT<br>GACACTATTTGGCTGACGTCCAAGTTCGCAGAT<br>CTAATGCCCCGACTATCGAGCTTCCAAAGCCTCC<br>GCCGCCACGAGCAAAAGAAACCGCATTTCACGA<br>CATCAGAGCCGACAATCCCCAACACCCTTCACT<br>GGTCAGTATGTACTACGGCGACTCTCTATTCATA<br>GGCGGAGATCAACCGGCAAACAAAGATAGGCA<br>GTCTCCAGTTCTGGCCCATCTCTATGAGCAGTTT<br>AAGGTCTCGAGCACCGAATCTTTGCAAGATATT<br>CCCGTATGACCCAGACCCGAAATACAATACCAA<br>GTTCAGCGCCGTTCACTTCGCAAGCGTAAGAGT<br>TCTAAAGACCGTACGCAGGCTCGCTGTGCTCTCC<br>TAGGTCATGGTTTGCTGACGATGTACAACTGAC<br>AAAAATCCCATTCCCATCCGCACCCAGGTTCTA<br>GAGGTAGTCGCCAACCCGTCCAACGCAAACTCC<br>AGATCTCCACCCAATTGCCTTTCAGGTGGTGCCT<br>CGATCCTTGCCGTCTAAATTATAGCCAGCCTGTG<br>CCCATTTAACCGACGGTTAAGTAATGAGAAAAA<br>GCGTTCCTGGCGAGATGCAGGCACGGGGGTGGG<br>AATCGAATGGCCAGGCGCCGCAGTGCTCCGTGC<br>TACCTGCCTCTCCCCCGATATCAGCAGTGACGGC<br>GACTCTAGCTCGGGCTCTGTTCCACCATTTACTC<br>TGGACCGGCCCGCCCCGCACAAGACGACACTCA<br>AGACAGCTGTGGCCCCCCTAACATCTACTCCGA<br>ACGACTCACCGCGCACTCATCTCACGTATCGTTC<br>GATGCCCCAGACGCCACCCACTATTATGTCCGTT<br>GCAGCTCAGGCGCGTGCACAATGCCTCCCAGAC<br>CTACACTCCACCACGACTAAGAACCCCCTTCACT<br>TGCTGCATTCTGAAGAATCTAAGGACTCCGCCA<br>CGGGTATCGACCATGAGCTATGCGACATTGTTC<br>AGGACTCGGTCAAGAAACTACGCTAGCATAC | taatacgactc<br>actataggg | gtatgctagcg<br>tagtttctt |
| **pJet1.2-atp5b** | TAATACGACTCACTATAGGTCTCCACCCGGATTC<br>CGCCATGTTGAGTCTTGTGGGGCGTGTGGCCTCG<br>GCCTCGGCCTCCGGGGCCTTGCGGGGACTCAGC<br>CCTTCGGCGGCTCTGCCACAGGCGCAGCTTCTAC<br>TGCGAGCTGCTCCCGCCGGGGTTCATCCTGCCA<br>GAGACTATGCGGCGCAGGCGTCTGCGGCCCCGA<br>AGGCAGGCACTGCCACCGGGCGAATCGTGGCAG | taatacgactc<br>actataggtct<br>ccacccgg | cattaataaatt<br>gatctttttaat<br>aataatcctgtt<br>tctaacaatttc<br>ctaaatgtggc<br>c |

| | | | |
|---|---|---|---|
| | TCATCGGCGCTGTGGTGGACGTCCAGTTCGATG<br>AGGGATTACCACCCATCCTAAATGCCCTGGAAG<br>TGCAAGGCAGGGACAGCAGACTGGTTTTGGAGG<br>TGGCCCAGCATTTGGGGGAGAGCACGGTCAGAA<br>CTATTGCTATGGATGGCACTGAAGGCTTGGTTA<br>GAGGCCAGAAAGTACTGGATTCAGGGGCACCAA<br>TCAAAATTCCTGTTGGTCCTGAGACCTTGGGCAG<br>AATCATGAATGTCATTGGAGAACCTATTGATGA<br>GAGAGGTCCTATCAAAACCAAACAATTTGCTCC<br>TATTCATGCTGAGGCTCCTGAGTTCATAGAGATG<br>AGTGTTGAGCAGGAGATTCTGGTGACTGGGATA<br>AAGGTTGTGGATCTGCTGGCCCCATACGCCAAG<br>GGTGGGAAAATCGGACTCTTTGGAGGTGCTGGC<br>GTTGGAAAGACAGTACTGATCATGGAGCTAATC<br>AACAATGTCGCCAAAGCCCATGGTGGTTACTCT<br>GTATTTGCTGGTGTTGGTGAGAGGACCCGTGAG<br>GGCAATGATTTATACCATGAAATGATTGAATCT<br>GGTGTTATCAATCTAAAAGATGCCACTTCCAAG<br>GTAGCGTTGGTATATGGACAGATGAACGAACCA<br>CCTGGCGCTCGAGCCCGGGTAGCTCTGACTGGT<br>TTGACCGTTGCTGAATACTTCAGAGACCAGGAG<br>GGCCAAGATGTCCTGCTGTTTATTGACAACATCT<br>TCCGCTTTACCCAGGCTGGCTCAGAGGTGTCTGC<br>CTTATTGGGCAGAATCCCTTCTGCTGTAGGCTAC<br>CAGCCCACCCTAGCCACCGACATGGGCACAATG<br>CAGGAAAGGATCACCACCACCAAGAAGGGATC<br>GATCACCTCGGTGCAGGCTATCTATGTGCCTGCT<br>GATGACCTGACTGACCCTGCCCCTGCAACCACC<br>TTTGCCCATTTGGATGCTACCACTGTGTTGTCCC<br>GGGCTATTGCTGAGTTGGGCATCTATCCAGCTGT<br>GGATCCACTGGACTCCACCTCTCGAATTATGGAT<br>CCCAACATTGTTGGCAATGAGCATTATGACGTC<br>GCCCGAGGAGTGCAGAAAATCCTGCAGGACTAC<br>AAATCTCTCCAGGACATCATTGCCATCTTGGGTA<br>TGGATGAACTTTCTGAGGAAGATAAATTGACTG<br>TGTCCCGGGCAAGAAAGATACAGCGCTTCTTGT<br>CTCAGCCATTCCAAGTTGCTGAGGTCTTCACGGG<br>TCACATGGGGAAGCTGGTGCCCTTGAAGGAGAC<br>CATTAAAGGATTCCAGCAGATTTTAGCAGGTGA<br>ATATGACCATCTCCCAGAACAAGCCTTCTACAT<br>GGTGGGACCCATTGAAGAAGCCGTGGCAAAGGC<br>TGATAAGCTGGCAGAAGAGCACGGGTCGTGAGG<br>GACTCCAGCCAAAGGCAGCACTGCAACTGATCT<br>CTCCATATCAAGCGAGAGCTCAGGTTTCCTTCCA<br>TGCAGGCCACACAAGAGCCTTGATTGAAGATGT<br>GATGTTCTCTCTGAAGAGTATTTAAAGTTTTCAA<br>TAAAGTATATACCCCTCATTTATGTCTGTTTATG<br>TTGCTTCTGAAACAGCTTATAATTGAGCTCACAG<br>TGGTGGGCCACATTTAGGAAATTGTTAGAAACA<br>GGATTATTATTAAAAAGATCAATTTATTAATG | | |
| **pJet1.2-scg3** | TAATACGACTCACTATAGGGAGCATGCGCCTCC<br>TCCTCATTTCCTCTCCTGGAGCTAGTGGAGTAAA<br>GCTACGCCCAGGCCCGCGTCCGCTGGCGGCGCA<br>GGAACTTCAGCACCCGCGGGGCGGACAGCGCCT<br>ACCGCACCTGCTCACCTGCTCTGGGCGCCAGAA | taatacgactc<br>actataggga<br>gcatgcgc | tcaaggatatg<br>aagagacgtg<br>ggtttatttccg |

| | |
|---|---|
| | GAGCCTGCATCCTCCTTCCAGCCCGGAGCAACT<br>GCGCCGGAGGCGCCCAGACCCTCTCCCTTCCCG<br>CACCCAGGCTCCTGTCCCTTCCAGCTTCTTAACT<br>CCCCTTCTCATTCATAACAAAAGCTACAGCTCAG<br>GGGCCCAGCGCCAAGCTCTTTCCAGCAAAGCAC<br>AGAAGAGCAAGAAAGAATGGGGTTCCTTTGGAC<br>CGGCTCTTGGATACTGGTGTTGGTGCTCAACAGC<br>GGCCCAATTCAAGCTTTCCCCAAACCCGAAGGC<br>AGCCAAGACAAATCCCTGCATAATAGAGAATTA<br>AGTGCAGAAAGACCTTTGAATGAACAGATCGCT<br>GAGGCAGAGGCAGACAAGATTAAAAAGGCATT<br>CCCTTCAGAAAGCAAGCCGAGTGAAAGCAATTA<br>TTCTTCTGTCGATAACTTGAATCTGCTGAGGGCA<br>ATAACAGAAAAGGAAACCGTTGAGAAAGAGAG<br>ACAATCCATAAGAAGCCCCCCGTTTGATAACCA<br>ACTGAACGTGGAAGACGCTGATTCAACCAAAAA<br>TCGGAAACTGATCGATGAGTACGATTCCACCAA<br>GAGTGGACTGGACCACAAGTTTCAAGATGACCC<br>AGACGGCCTTCATCAACTGGATGGAACTCCTTT<br>AACTGCTGAAGACATCGTCCATAAGATTGCCAC<br>CAGGATTTATGAGGAGAACGACAGAGGAGTGTT<br>TGACAAAATTGTTTCTAAACTGCTGAATCTTGGC<br>CTGATCACTGAAAGCCAGGCACATACTCTGGAA<br>GATGAAGTAGCAGAAGCTTTACAAAAACTGATT<br>TCAAAAGAGGCCAACAATTATGAGGAGACCCTG<br>GATAAACCCACAAGCAGGACCGAGAATCAGGA<br>TGGGAAAATACCAGAGAAAGTGACTCCGGTGGC<br>AGCAGTCCAAGATGGCTTCACTAACCGTGAAAA<br>CGATGAGACGGTGTCTAACACCTTGACCTTGTCC<br>AATGGCTTGGAAAGGAGAACTAACCCCCACAGG<br>GAAGACGACTTTGAGGAACTCCAGTATTTCCCC<br>AACTTCTATGCACTACTGACAAGCATCGACTCA<br>GAAAAAGAAGCAAAAGAGAAAGAAACCCTGAT<br>CACCATCATGAAGACATTGATTGACTTCGTGAA<br>AATGATGGTGAAATACGGTACGATATCTCCAGA<br>GGAAGGCGTGTCCTACCTTGAAAACTTGGATGA<br>AACAATTGCTCTGCAGACCAAGAACAAGCTAGA<br>AAAAAATACTACTGATAGCAAAACTCCACCAGA<br>GAAGAGTCAGGAAGAAACAGACAGTACCAAGG<br>AAGAAGCCGCCAAGATGGAAAAGGAATACGGA<br>AGCCTAAAAGACTCTACAAAAGATGATAACTCC<br>AACCTAGGAGGAAAGACAGATGAAGCCACAGG<br>GAAGACAGAAGCCTACTTGGAAGCCATTAGAAA<br>AAACATCGAATGGCTGAAGAAACATAACAAGA<br>AGGGCAACAAAGAAGATTACGACCTTTCAAAGA<br>TGAGGGACTTTATCAACCAACAAGCTGACGCTT<br>ATGTGGAGAAGGGCATCCTCGACAAGGAAGAA<br>GCCAACGCCATCAAACGCATCTACAGCAGCCTG<br>TGAAAATGGCGGGCAGCTTGAGCCTTCCTGTTG<br>TTCCAGCAAAAACAATATAGCTTACAAACTAAT<br>TCGGCGGTTAAAGGGTTACCAGCCCAGAAGTAT<br>TAGGATGTGCTGAATTTATAGTAGTTAATCCCTT<br>AGAAATGAGTAAAATAGAGCTCTCTTGCCATAA<br>ATACCTTATGAAAAGCAAAGCTGTAGAGAAGCC<br>GAGGTTTTTCTATATAGAATCCTTATTTCCTCTT |

| | | | |
|---|---|---|---|
| | GAATTTACATTTTGTAATCAGAGATGTGCTGCTC<br>TGGAAAAGACTCTAATGGGTTGAACATAAGTCT<br>GAACCTACTCCCCACTGTCCTCAGCCCCCTGAAG<br>CTCTGAGAGGCCCTGTCTCGGCATGCTAGACAC<br>CTGAGCACCTCACTGGATGTTTGTCATAGGATGT<br>CGTTTCCACTAGTCGATCTCTGTTGGGCACGGAA<br>ATAAACCCACGTCTCTTCATATCCTTGA | | |
| **pJet1.2-prnp** | TAATACGACTCACTATAGCCCCTTTCCACTCCCG<br>GCTCCCCCGCGTTGTCGGATCAGCAGACCGATT<br>CTGGGCGCTGCGTCGCATCGGTGGCAGGACTCC<br>TGAGTATATTTCAGAACTGAACCATTTCAACCG<br>AGCTGAAGCATTCTGCCTTCCTAGTGGTACCAGT<br>CCAATTTAGGAGAGCCAAGCAGACTATCAGTCA<br>TCATGGCGAACCTTGGCTACTGGCTGCTGGCCCT<br>CTTTGTGACTATGTGGACTGATGTCGGCCTCTGC<br>AAAAAGCGGCCAAAGCCTGGAGGGTGGAACAC<br>CGGTGGAAGCCGGTATCCCGGGCAGGGAAGCCC<br>TGGAGGCAACCGTTACCCACCTCAGGGTGGCAC<br>CTGGGGGCAGCCCCACGGTGGTGGCTGGGGACA<br>ACCCCATGGGGGCAGCTGGGGACAACCTCATGG<br>TGGTAGTTGGGGTCAGCCCCATGGCGGTGGATG<br>GGGCCAAGGAGGGGGTACCCATAATCAGTGGA<br>ACAAGCCCAGCAAACCAAAAACCAACCTCAAGC<br>ATGTGGCAGGGGCTGCGGCAGCTGGGGCAGTAG<br>TGGGGGGCCTTGGTGGCTACATGCTGGGGAGCG<br>CCATGAGCAGGCCCATGATCCATTTTGGCAACG<br>ACTGGGAGGACCGCTACTACCGTGAAAACATGT<br>ACCGCTACCCTAACCAAGTGTACTACAGGCCAG<br>TGGATCAGTACAGCAACCAGAACAACTTCGTGC<br>ACGACTGCGTCAATATCACCATCAAGCAGCACA<br>CGGTCACCACCACCACCAAGGGGGAGAACTTCA<br>CCGAGACCGATGTGAAGATGATGGAGCGCGTGG<br>TGGAGCAGATGTGCGTCACCCAGTACCAGAAGG<br>AGTCCCAGGCCTATTACGACGGGAGAAGATCCA<br>GCAGCACCGTGCTTTTCTCCTCCCCTCCTGTCAT<br>CCTCCTCATCTCCTTCCTCATCTTCCTGATCGTGG<br>GATGAGGGAGGCCTTCCTGCTTGTTCCTTCGCAT<br>TCTCGTGGTCTAGGCTGGGGGAGGGGTTATCCA<br>CCTGTAGCTCTTTCAATTGAGGTGGTTCTCATTC<br>TTGCTTCTCTGTGTCCCCCATAGGCTAATACCCC<br>TGGCACTGATGGGCCCTGGGAAATGTACAGTAG<br>ACCAGTTGCTCTTTGCTTCAGGTCCCTTTGATGG<br>AGTCTGTCATCAGCCAGTGCTAACACCGGGCCA<br>ATAAGAATATAACACCAAATAACTGCTGGCTAG<br>TTGGGGCTTTGTTTTGGTCTAGTGAATAAATACT<br>GGTGTATCCCCTGACTTGTACCCAGAGTACAAG<br>GTGACAGTGACACATGTAACTTAGCATAGGCAA<br>AGGGTTCTACAACCAAAGAAGCCACTGTTTGGG<br>GATGGCGCCCTGGAAAACAGCCTCCCACCTGGG<br>ATAGCTAGAGCATCCACACGTGGAATTCTTTCTT<br>TACTAACAAACGATAGCTGATTGAAGGCAACAG<br>GAAAAAAAAAATCAAATTGTCCTACTGACGTTG<br>AAAGCAAACCTTTGTTCATTCCCAGGGCACTAG<br>AATGATCTTTAGCCTTGCTTGGATTGAACTAGGA<br>GATCTTGACTCTGAGGAGAGCCAGCCCTGTAAA | taatacgactc<br>actatagcccc<br>tttccactcc | ccaggtgttca<br>tcatagcttcta<br>tttaatgtcagt<br>ctg |

| | | | |
|---|---|---|---|
| | AAGCTTGGTCCTCCTGTGACGGGAGGGATGGTT AAGGTACAAAGGCTAGAAACTTGAGTTTCTTCA TTTCTGTCTCACAATTATCAAAAGCTAGAATTAG CTTCTGCCCTATGTTTCTGTACTTCTATTTGAACT GGATAACAGAGAGACAATCTAAACATTCTCTTA GGCTGCAGATAAGAGAAGTAGGCTCCATTCCAA AGTGGGAAAGAAATTCTGCTAGCATTGTTTAAA TCAGGCAAAATTTGTTCCTGAAGTTGCTTTTTAC CCCAGCAGACATAAACTGCGATAGCTTCAGCTT GCACTGTGGATTTTCTGTATAGAATATATAAAAC ATAACTTCAAGCTTATGTCTTCTTTTTAAAACAT CTGAAGTATGGGACGCCCTGGCCGTTCCATCCA GTACTAAATGCTTACCGTGTGACCCTTGGGCTTT CAGCGTGCACTCAGTTCCGTAGGATTCCAAAGC AGACCCCTAGCTGGTCTTTGAATCTGCATGTACT TCACGTTTTCTATATTTGTAACTTTGCATGTATTT TGTTTTGTCATATAAAAAGTTTATAAATGTTTGC TATCAGACTGACATTAAATAGAAGCTATGATGA ACACCTGG | | |
| **pJet1.2-atp5b::nluc** | TAATACGACTCACTATAAGGCCACCATGTTGAG TCTTGTGGGGCGTGTGGCCTCGGCCTCGGCCTCC GGGGCCTTGCGGGGACTCAGCCCTTCGGCGGCT CTGCCACAGGCGCAGCTTCTACTGCGAGCTGCT CCCGCCGGGGTTCATCCTGCCAGAGACTATGCG GCGCAGGCGTCTGCGGCCCCGAAGGCAGGCACT GCCACCGGGCGAATCGTGGCAGTCATCGGCGCT GTGGTGGACGTCCAGTTCGATGAGGGATTACCA CCCATCCTAAATGCCCTGGAAGTGCAAGGCAGG GACAGCAGACTGGTTTTGGAGGTGGCCCAGCAT TTGGGGGAGAGCACGGTCAGAACTATTGCTATG GATGGCACTGAAGGCTTGGTTAGAGGCCAGAAA GTACTGGATTCAGGGGCACCAATCAAAATTCCT GTTGGTCCTGAGACCTTGGGCAGAATCATGAAT GTCATTGGAGAACCTATTGATGAGAGAGGTCCT ATCAAAACCAAACAATTTGCTCCTATTCATGCTG AGGCTCCTGAGTTCATAGAGATGAGTGTTGAGC AGGAGATTCTGGTGACTGGGATAAAGGTTGTGG ATCTGCTGGCCCCATACGCCAAGGGTGGGAAAA TCGGACTCTTTGGAGGTGCTGGCGTTGGAAAGA CAGTACTGATCATGGAGCTAATCAACAATGTCG CCAAAGCCCATGGTGGTTACTCTGTATTTGCTGG TGTTGGTGAGAGGACCCGTGAGGGCAATGATTT ATACCATGAAATGATTGAATCTGGTGTTATCAAT CTAAAAGATGCCACTTCCAAGGTAGCGTTGGTA TATGGACAGATGAACGAACCACCTGGCGCTCGA GCCCGGGTAGCTCTGACTGGTTTGACCGTTGCTG AATACTTCAGAGACCAGGAGGGCCAAGATGTCC TGCTGTTTATTGACAACATCTTCCGCTTTACCCA GGCTGGCTCAGAGGTGTCTGCCTTATTGGGCAG AATCCCTTCTGCTGTAGGCTACCAGCCCACCCTA GCCACCGACATGGGCACAATGCAGGAAAGGATC ACCACCACCAAGAAGGGATCGATCACCTCGGTG CAGGCTATCTATGTGCCTGCTGATGACCTGACTG ACCCTGCCCCTGCAACCACCTTTGCCCATTTGGA TGCTACCACTGTGTTGTCCCGGGCTATTGCTGAG | TAATACGACTCACTATAAGGCCACCATGTTGAGTCTTGTG | ttatttgtacaatgatcccccacccgc |

| | | | |
|---|---|---|---|
| | TTGGGCATCTATCCAGCTGTGGATCCACTGGACT<br>CCACCTCTCGAATTATGGATCCCAACATTGTTGG<br>CAATGAGCATTATGACGTCGCCCGAGGAGTGCA<br>GAAAATCCTGCAGGACTACAAATCTCTCCAGGA<br>CATCATTGCCATCTTGGGTATGGATGAACTTTCT<br>GAGGAAGATAAATTGACTGTGTCCCGGGCAAGA<br>AAGATACAGCGCTTCTTGTCTCAGCCATTCCAAG<br>TTGCTGAGGTCTTCACGGGTCACATGGGGAAGC<br>TGGTGCCCTTGAAGGAGACCATTAAAGGATTCC<br>AGCAGATTTTAGCAGGTGAATATGACCATCTCC<br>CAGAACAAGCCTTCTACATGGTGGGACCCATTG<br>AAGAAGCCGTGGCAAAGGCTGATAAGCTGGCA<br>GAAGAGCACGGGTCGATGGTCTTCACACTCGAA<br>GATTTCGTTGGGGACTGGCGACAGACAGCCGGC<br>TACAACCTGGACCAAGTCCTTGAACAGGGAGGT<br>GTGTCCAGTTTGTTTCAGAATCTCGGGGTGTCCG<br>TAACTCCGATCCAAAGGATTGTCCTGAGCGGTG<br>AAAATGGGCTGAAGATCGACATCCATGTCATCA<br>TCCCGTATGAAGGTCTGAGCGGCGACCAAATGG<br>GCCAGATCGAAAAAATTTTTAAGGTGGTGTACC<br>CTGTGGATGATCATCACTTTAAGGTGATCCTGCA<br>CTATGGCACACTGGTAATCGACGGGGTTACGCC<br>GAACATGATCGACTATTTCGGACGGCCGTATGA<br>AGGCATCGCCGTGTTCGACGGCAAAAAGATCAC<br>TGTAACAGGGACCCTGTGGAACGGCAACAAAAT<br>TATCGACGAGCGCCTGATCAACCCCGACGGCTC<br>CCTGCTGTTCCGAGTAACCATCAACGGAGTGAC<br>CGGCTGGCGGCTGTGCGAACGCATTCTGGCGGG<br>TGGGGGATCATTGTACAAATAA | | |
| **pJet1.2-atp5b::moon** | TAATACGACTCACTATAAGGCCACCATGTTGAG<br>TCTTGTGGGGCGTGTGGCCTCGGCCTCGGCCTCC<br>GGGGCCTTGCGGGGACTCAGCCCTTCGGCGGCT<br>CTGCCACAGGCGCAGCTTCTACTGCGAGCTGCT<br>CCCGCCGGGGTTCATCCTGCCAGAGACTATGCG<br>GCGCAGGCGTCTGCGGCCCCGAAGGCAGGCACT<br>GCCACCGGGCGAATCGTGGCAGTCATCGGCGCT<br>GTGGTGGACGTCCAGTTCGATGAGGGATTACCA<br>CCCATCCTAAATGCCCTGGAAGTGCAAGGCAGG<br>GACAGCAGACTGGTTTTGGAGGTGGCCCAGCAT<br>TTGGGGGAGAGCACGGTCAGAACTATTGCTATG<br>GATGGCACTGAAGGCTTGGTTAGAGGCCAGAAA<br>GTACTGGATTCAGGGGCACCAATCAAAATTCCT<br>GTTGGTCCTGAGACCTTGGGCAGAATCATGAAT<br>GTCATTGGAGAACCTATTGATGAGAGAGGTCCT<br>ATCAAAACCAAACAATTTGCTCCTATTCATGCTG<br>AGGCTCCTGAGTTCATAGAGATGAGTGTTGAGC<br>AGGAGATTCTGGTGACTGGGATAAAGGTTGTGG<br>ATCTGCTGGCCCCATACGCCAAGGGTGGGAAAA<br>TCGGACTCTTTGGAGGTGCTGGCGTTGGAAAGA<br>CAGTACTGATCATGGAGCTAATCAACAATGTCG<br>CCAAAGCCCATGGTGGTTACTCTGTATTTGCTGG<br>TGTTGGTGAGAGGACCCGTGAGGGCAATGATTT<br>ATACCATGAAATGATTGAATCTGGTGTTATCAAT<br>CTAAAAGATGCCACTTCCAAGGTAGCGTTGGTA<br>TATGGACAGATGAACGAACCACCTGGCGCTCGA | cDNA template was digested with HindIII to linearize prior to IVT | cDNA template was digested with HindIII to linearize prior to IVT |

| | |
|---|---|
| | GCCCGGGTAGCTCTGACTGGTTTGACCGTTGCTG<br>AATACTTCAGAGACCAGGAGGGCCAAGATGTCC<br>TGCTGTTTATTGACAACATCTTCCGCTTTACCCA<br>GGCTGGCTCAGAGGTGTCTGCCTTATTGGGCAG<br>AATCCCTTCTGCTGTAGGCTACCAGCCCACCCTA<br>GCCACCGACATGGGCACAATGCAGGAAAGGATC<br>ACCACCACCAAGAAGGGATCGATCACCTCGGTG<br>CAGGCTATCTATGTGCCTGCTGATGACCTGACTG<br>ACCCTGCCCCTGCAACCACCTTTGCCCATTTGGA<br>TGCTACCACTGTGTTGTCCCGGGCTATTGCTGAG<br>TTGGGCATCTATCCAGCTGTGGATCCACTGGACT<br>CCACCTCTCGAATTATGGATCCCAACATTGTTGG<br>CAATGAGCATTATGACGTCGCCCGAGGAGTGCA<br>GAAAATCCTGCAGGACTACAAATCTCTCCAGGA<br>CATCATTGCCATCTTGGGTATGGATGAACTTTCT<br>GAGGAAGATAAATTGACTGTGTCCCGGGCAAGA<br>AAGATACAGCGCTTCTTGTCTCAGCCATTCCAAG<br>TTGCTGAGGTCTTCACGGGTCACATGGGGAAGC<br>TGGTGCCCTTGAAGGAGACCATTAAAGGATTCC<br>AGCAGATTTTAGCAGGTGAATATGACCATCTCC<br>CAGAACAAGCCTTCTACATGGTGGGACCCATTG<br>AAGAAGCCGTGGCAAAGGCTGATAAGCTGGCA<br>GAAGAGCACGGGTCGATGCGTACGGGCAAGAA<br>TGAGCAGGAACTGCTGGAACTGGACAAATGGGC<br>AAGCCTGGGGAGCGGCTCCGGAAAGAACGAGC<br>AGGAACTGCTGGAACTGGACAAGTGGGCTTCTC<br>TGGGCTCCGGGTCCGGGAAAAACGAACAGGAG<br>CTGCTGGAGCTGGATAAGTGGGCCAGCCTGGGC<br>TCCGGGTCCGGGAAGAATGAACAGGAACTGCTG<br>GAACTGGATAAATGGGCCTCTCTGGGCAGCGGG<br>TCCGGAAAGAATGAACAGGAGCTGCTGGAACTG<br>GACAAGTGGGCGTCCCTGGGGAGCGGGTCCGGA<br>AAGAATGAACAGGAACTGCTGGAACTGGATAA<br>GTGGGCTTCCCTGGGAAGCGGCTCCGGAAAAAA<br>CGAGCAGGAGCTGCTGGAGCTGGACAAGTGGGC<br>CTCTCTGGGCTCCGGCTCCGGGAAGAATGAACA<br>GGAGCTGCTGGAACTGGATAAGTGGGCCTCTCT<br>GGGGTCCGGAAGCGGTAAAAACGAGCAGGAAC<br>TGCTGGAACTGGACAAGTGGGCAAGCCTGGGGT<br>CTGGCTCCGGCAAAAACGAACAGGAACTGCTGG<br>AACTGGACAAATGGGCCAGCCTGGGCTCCGGGT<br>CCGGGAAAAATGAGCAGGAACTGCTGGAACTG<br>GATAAATGGGCGAGCCTGGGCAGCGGGTCCGGA<br>AAAAACGAACAGGAGCTGCTGGAGCTGGACAA<br>ATGGGCGAGTCTGGGCTCCGGAAGCGGCAAGAA<br>TGAACAGGAACTGCTGGAGCTGGACAAATGGGC<br>CAGCCTGGGCAGCGGCTCCGGAAAGAATGAGCA<br>GGAACTGCTGGAGCTGGACAAATGGGCGTCTCT<br>GGGCTCCGGCTCCGGGAAAAACGAGCAGGAACT<br>GCTGGAGCTGGACAAGTGGGCTTCCCTGGGCAG<br>CGGCTCCGGAAAAAATGAACAGGAACTGCTGGA<br>ACTGGACAAGTGGGCCAGCCTGGGGTCCGGGTC<br>TGGGAAGAACGAGCAGGAACTGCTGGAGCTGG<br>ACAAGTGGGCTTCTCTGGGGTCTGGCTCTGGCA<br>AGAACGAGCAGGAACTGCTGGAACTGGACAAG |

| | |
|---|---|
| | TGGGCGAGCCTGGGCAGCGGGTCTGGCAAGAAT GAACAGGAGCTGCTGGAACTGGACAAATGGGCC AGCCTGGGGTCTGGGTCTGGAAAGAATGAGCAG GAACTGCTGGAGCTGGACAAGTGGGCCTCTCTG GGCTCTGGGTCTGGCAAGAACGAGCAGGAGCTG CTGGAGCTGGATAAGTGGGCGTCCCTGGGGTCC GGGTCCGGCAAGAATGAACAGGAGCTGCTGGA GCTGGATAAATGGGCTTCCCTGGGCTCTGGGTCT GGGAAGAATGAGCAGGAGCTGCTGGAACTGGA TAAGTGGGCGAGCCTGGGCAGCGGGTCTGGCAA GAATGAGCAGGAACTGCTGGAGCTGGACAAGTG GGCTTCTCTGTAA |

**Supplementary Table S3: Sequences of synaptic and *CLN3* mRNAs used for in vitro reconstitutions.**

| Table S3 Sequences of synaptic and *CLN3* mRNAs used for in vitro reconstitutions | | | | |
|---|---|---|---|---|
| **Gene name** | **Transcript ID** | **Species** | **Transcript sequence (5->3'))** | **DNA seq (T7 promoter [bold] added at 5' end) (5->3')** |
| ***Atp5b*** | NM_016774.3 | *Mus musculus* | GTCTCCACCCGGATTCCGCCATGTT<br>GAGTCTTGTGGGGCGTGTGGCCTC<br>GGCCTCGGCCTCCGGGGCCTTGCG<br>GGGACTCAGCCCTTCGGCGGCTCT<br>GCCACAGGCGCAGCTTCTACTGCG<br>AGCTGCTCCCGCCGGGGTTCATCCT<br>GCCAGAGACTATGCGGCGCAGGCG<br>TCTGCGGCCCCGAAGGCAGGCACT<br>GCCACCGGGCGAATCGTGGCAGTC<br>ATCGGCGCTGTGGTGGACGTCCAG<br>TTCGATGAGGGATTACCACCCATCC<br>TAAATGCCCTGGAAGTGCAAGGCA<br>GGGACAGCAGACTGGTTTTGGAGG<br>TGGCCCAGCATTTGGGGGAGAGCA<br>CGGTCAGAACTATTGCTATGGATG<br>GCACTGAAGGCTTGGTTAGAGGCC<br>AGAAAGTACTGGATTCAGGGGCAC<br>CAATCAAAATTCCTGTTGGTCCTGA<br>GACCTTGGGCAGAATCATGAATGT<br>CATTGGAGAACCTATTGATGAGAG<br>AGGTCCTATCAAAACCAAACAATT<br>TGCTCCTATTCATGCTGAGGCTCCT<br>GAGTTCATAGAGATGAGTGTTGAG<br>CAGGAGATTCTGGTGACTGGGATA<br>AAGGTTGTGGATCTGCTGGCCCCAT<br>ACGCCAAGGGTGGAAAATCGGAC<br>TCTTTGGAGGTGCTGGCGTTGGAA<br>AGACAGTACTGATCATGGAGCTAA<br>TCAACAATGTCGCCAAAGCCCATG<br>GTGGTTACTCTGTATTTGCTGGTGT<br>TGGTGAGAGGACCCGTGAGGGCAA<br>TGATTTATACCATGAAATGATTGAA<br>TCTGGTGTTATCAATCTAAAAGATG<br>CCACTTCCAAGGTAGCGTTGGTATA<br>TGGACAGATGAACGAACCACCTGG<br>CGCTCGAGCCCGGGTAGCTCTGAC<br>TGGTTTGACCGTTGCTGAATACTTC<br>AGAGACCAGGAGGGCCAAGATGTC<br>CTGCTGTTTATTGACAACATCTTCC<br>GCTTTACCCAGGCTGGCTCAGAGG | **TAATACGACTCACTATAG**GTCTCC<br>ACCCGGATTCCGCCATGTTGAGTCT<br>TGTGGGGCGTGTGGCCTCGGCCTC<br>GGCCTCCGGGGCCTTGCGGGGACT<br>CAGCCCTTCGGCGGCTCTGCCACA<br>GGCGCAGCTTCTACTGCGAGCTGCT<br>CCCGCCGGGGTTCATCCTGCCAGA<br>GACTATGCGGCGCAGGCGTCTGCG<br>GCCCCGAAGGCAGGCACTGCCACC<br>GGGCGAATCGTGGCAGTCATCGGC<br>GCTGTGGTGGACGTCCAGTTCGAT<br>GAGGGATTACCACCCATCCTAAAT<br>GCCCTGGAAGTGCAAGGCAGGGAC<br>AGCAGACTGGTTTTGGAGGTGGCC<br>CAGCATTTGGGGGAGAGCACGGTC<br>AGAACTATTGCTATGGATGGCACT<br>GAAGGCTTGGTTAGAGGCCAGAAA<br>GTACTGGATTCAGGGGCACCAATC<br>AAAATTCCTGTTGGTCCTGAGACCT<br>TGGGCAGAATCATGAATGTCATTG<br>GAGAACCTATTGATGAGAGAGGTC<br>CTATCAAAACCAAACAATTTGCTCC<br>TATTCATGCTGAGGCTCCTGAGTTC<br>ATAGAGATGAGTGTTGAGCAGGAG<br>ATTCTGGTGACTGGGATAAAGGTT<br>GTGGATCTGCTGGCCCCATACGCC<br>AAGGGTGGGAAAATCGGACTCTTT<br>GGAGGTGCTGGCGTTGGAAAGACA<br>GTACTGATCATGGAGCTAATCAAC<br>AATGTCGCCAAAGCCCATGGTGGT<br>TACTCTGTATTTGCTGGTGTTGGTG<br>AGAGGACCCGTGAGGGCAATGATT<br>TATACCATGAAATGATTGAATCTG<br>GTGTTATCAATCTAAAAGATGCCA<br>CTTCCAAGGTAGCGTTGGTATATGG<br>ACAGATGAACGAACCACCTGGCGC<br>TCGAGCCCGGGTAGCTCTGACTGG<br>TTTGACCGTTGCTGAATACTTCAGA<br>GACCAGGAGGGCCAAGATGTCCTG<br>CTGTTTATTGACAACATCTTCCGCT |

| | | | | |
|---|---|---|---|---|
| | | | TGTCTGCCTTATTGGGCAGAATCCC<br>TTCTGCTGTAGGCTACCAGCCCACC<br>CTAGCCACCGACATGGGCACAATG<br>CAGGAAAGGATCACCACCACCAAG<br>AAGGGATCGATCACCTCGGTGCAG<br>GCTATCTATGTGCCTGCTGATGACC<br>TGACTGACCCTGCCCCTGCAACCAC<br>CTTTGCCCATTTGGATGCTACCACT<br>GTGTTGTCCCGGGCTATTGCTGAGT<br>TGGGCATCTATCCAGCTGTGGATCC<br>ACTGGACTCCACCTCTCGAATTATG<br>GATCCCAACATTGTTGGCAATGAG<br>CATTATGACGTCGCCCGAGGAGTG<br>CAGAAAATCCTGCAGGACTACAAA<br>TCTCTCCAGGACATCATTGCCATCT<br>TGGGTATGGATGAACTTTCTGAGG<br>AAGATAAATTGACTGTGTCCCGGG<br>CAAGAAAGATACAGCGCTTCTTGT<br>CTCAGCCATTCCAAGTTGCTGAGGT<br>CTTCACGGGTCACATGGGGAAGCT<br>GGTGCCCTTGAAGGAGACCATTAA<br>AGGATTCCAGCAGATTTTAGCAGG<br>TGAATATGACCATCTCCCAGAACA<br>AGCCTTCTACATGGTGGGACCCATT<br>GAAGAAGCCGTGGCAAAGGCTGAT<br>AAGCTGGCAGAAGAGCACGGGTCG<br>TGAGGGACTCCAGCCAAAGGCAGC<br>ACTGCAACTGATCTCTCCATATCAA<br>GCGAGAGCTCAGGTTTCCTTCCATG<br>CAGGCCACACAAGAGCCTTGATTG<br>AAGATGTGATGTTCTCTCTGAAGA<br>GTATTTAAAGTTTCAATAAAGTAT<br>ATACCCCTCATTTATGTCTGTTTAT<br>GTTGCTTCTGAAACAGCTTATAATT<br>GAGCTCACAGTGGTGGGCCACATT<br>TAGGAAATTGTTAGAAACAGGATT<br>ATTATTAAAAAGATCAATTTATTAA<br>TG | TTACCCAGGCTGGCTCAGAGGTGT<br>CTGCCTTATTGGGCAGAATCCCTTC<br>TGCTGTAGGCTACCAGCCCACCCTA<br>GCCACCGACATGGGCACAATGCAG<br>GAAAGGATCACCACCACCAAGAAG<br>GGATCGATCACCTCGGTGCAGGCT<br>ATCTATGTGCCTGCTGATGACCTGA<br>CTGACCCTGCCCCTGCAACCACCTT<br>TGCCCATTTGGATGCTACCACTGTG<br>TTGTCCCGGGCTATTGCTGAGTTGG<br>GCATCTATCCAGCTGTGGATCCACT<br>GGACTCCACCTCTCGAATTATGGAT<br>CCCAACATTGTTGGCAATGAGCATT<br>ATGACGTCGCCCGAGGAGTGCAGA<br>AAATCCTGCAGGACTACAAATCTC<br>TCCAGGACATCATTGCCATCTTGGG<br>TATGGATGAACTTTCTGAGGAAGA<br>TAAATTGACTGTGTCCCGGGCAAG<br>AAAGATACAGCGCTTCTTGTCTCAG<br>CCATTCCAAGTTGCTGAGGTCTTCA<br>CGGGTCACATGGGGAAGCTGGTGC<br>CCTTGAAGGAGACCATTAAAGGAT<br>TCCAGCAGATTTTAGCAGGTGAAT<br>ATGACCATCTCCCAGAACAAGCCT<br>TCTACATGGTGGGACCCATTGAAG<br>AAGCCGTGGCAAAGGCTGATAAGC<br>TGGCAGAAGAGCACGGGTCGTGAG<br>GGACTCCAGCCAAAGGCAGCACTG<br>CAACTGATCTCTCCATATCAAGCGA<br>GAGCTCAGGTTTCCTTCCATGCAGG<br>CCACACAAGAGCCTTGATTGAAGA<br>TGTGATGTTCTCTCTGAAGAGTATT<br>TAAAGTTTCAATAAAGTATATACC<br>CCTCATTTATGTCTGTTTATGTTGCT<br>TCTGAAACAGCTTATAATTGAGCTC<br>ACAGTGGTGGGCCACATTTAGGAA<br>ATTGTTAGAAACAGGATTATTATTA<br>AAAAGATCAATTTATTAATG |
| ***Prnp*** | NM_001164790.1 | *Mus musculus* | CCCCTTTCCACTCCCGGCTCCCCCG<br>CGTTGTCGGATCAGCAGACCGATT<br>CTGGGCGCTGCGTCGCATCGGTGG<br>CAGGACTCCTGAGTATATTTCAGA<br>ACTGAACCATTTCAACCGAGCTGA<br>AGCATTCTGCCTTCCTAGTGGTACC<br>AGTCCAATTTAGGAGAGCCAAGCA<br>GACTATCAGTCATCATGGCGAACC<br>TTGGCTACTGGCTGCTGGCCCTCTT<br>TGTGACTATGTGGACTGATGTCGGC<br>CTCTGCAAAAAGCGGCCAAAGCCT<br>GGAGGGTGGAACACCGGTGGAAGC<br>CGGTATCCCGGGCAGGGAAGCCCT<br>GGAGGCAACCGTTACCCACCTCAG<br>GGTGGCACCTGGGGGCAGCCCCAC<br>GGTGGTGGCTGGGGACAACCCCAT<br>GGGGGCAGCTGGGGACAACCTCAT | **TAATACGACTCACTATAG**CCCCTT<br>TCCACTCCCGGCTCCCCCGCGTTGT<br>CGGATCAGCAGACCGATTCTGGGC<br>GCTGCGTCGCATCGGTGGCAGGAC<br>TCCTGAGTATATTTCAGAACTGAAC<br>CATTTCAACCGAGCTGAAGCATTCT<br>GCCTTCCTAGTGGTACCAGTCCAAT<br>TTAGGAGAGCCAAGCAGACTATCA<br>GTCATCATGGCGAACCTTGGCTACT<br>GGCTGCTGGCCCTCTTTGTGACTAT<br>GTGGACTGATGTCGGCCTCTGCAA<br>AAAGCGGCCAAAGCCTGGAGGGTG<br>GAACACCGGTGGAAGCCGGTATCC<br>CGGGCAGGGAAGCCCTGGAGGCAA<br>CCGTTACCCACCTCAGGGTGGCAC<br>CTGGGGGCAGCCCCACGGTGGTGG<br>CTGGGGACAACCCCATGGGGGCAG |

| | | | | |
|---|---|---|---|---|
| | | | GGTGGTAGTTGGGGTCAGCCCCAT<br>GGCGGTGGATGGGGCCAAGGAGGG<br>GGTACCCATAATCAGTGGAACAAG<br>CCCAGCAAACCAAAAACCAACCTC<br>AAGCATGTGGCAGGGGCTGCGGCA<br>GCTGGGGCAGTAGTGGGGGGCCTT<br>GGTGGCTACATGCTGGGGAGCGCC<br>ATGAGCAGGCCCATGATCCATTTTG<br>GCAACGACTGGGAGGACCGCTACT<br>ACCGTGAAAACATGTACCGCTACC<br>CTAACCAAGTGTACTACAGGCCAG<br>TGGATCAGTACAGCAACCAGAACA<br>ACTTCGTGCACGACTGCGTCAATAT<br>CACCATCAAGCAGCACACGGTCAC<br>CACCACCACCAAGGGGGAGAACTT<br>CACCGAGACCGATGTGAAGATGAT<br>GGAGCGCGTGGTGGAGCAGATGTG<br>CGTCACCCAGTACCAGAAGGAGTC<br>CCAGGCCTATTACGACGGGAGAAG<br>ATCCAGCAGCACCGTGCTTTTCTCC<br>TCCCCTCCTGTCATCCTCCTCATCT<br>CCTTCCTCATCTTCCTGATCGTGGG<br>ATGAGGGAGGCCTTCCTGCTTGTTC<br>CTTCGCATTCTCGTGGTCTAGGCTG<br>GGGGAGGGGTTATCCACCTGTAGC<br>TCTTTCAATTGAGGTGGTTCTCATT<br>CTTGCTTCTCTGTGTCCCCCATAGG<br>CTAATACCCCTGGCACTGATGGGC<br>CCTGGGAAATGTACAGTAGACCAG<br>TTGCTCTTTGCTTCAGGTCCCTTTG<br>ATGGAGTCTGTCATCAGCCAGTGCT<br>AACACCGGGCCAATAAGAATATAA<br>CACCAAATAACTGCTGGCTAGTTG<br>GGGCTTTGTTTTGGTCTAGTGAATA<br>AATACTGGTGTATCCCCTGACTTGT<br>ACCCAGAGTACAAGGTGACAGTGA<br>CACATGTAACTTAGCATAGGCAAA<br>GGGTTCTACAACCAAAGAAGCCAC<br>TGTTTGGGGATGGCGCCCTGGAAA<br>ACAGCCTCCCACCTGGGATAGCTA<br>GAGCATCCACACGTGGAATTCTTTC<br>TTTACTAACAAACGATAGCTGATTG<br>AAGGCAACAGGAAAAAAAAAAATC<br>AAATTGTCCTACTGACGTTGAAAG<br>CAAACCTTTGTTCATTCCCAGGGCA<br>CTAGAATGATCTTTAGCCTTGCTTG<br>GATTGAACTAGGAGATCTTGACTCT<br>GAGGAGAGCCAGCCCTGTAAAAAG<br>CTTGGTCCTCCTGTGACGGGAGGG<br>ATGGTTAAGGTACAAAGGCTAGAA<br>ACTTGAGTTTCTTCATTTCTGTCTC<br>ACAATTATCAAAAGCTAGAATTAG<br>CTTCTGCCCTATGTTTCTGTACTTCT<br>ATTTGAACTGGATAACAGAGAGAC<br>AATCTAAACATTCTCTTAGGCTGCA | CTGGGGACAACCTCATGGTGGTAG<br>TTGGGGTCAGCCCCATGGCGGTGG<br>ATGGGGCCAAGGAGGGGGTACCCA<br>TAATCAGTGGAACAAGCCCAGCAA<br>ACCAAAAACCAACCTCAAGCATGT<br>GGCAGGGGCTGCGGCAGCTGGGGC<br>AGTAGTGGGGGGCCTTGGTGGCTA<br>CATGCTGGGGAGCGCCATGAGCAG<br>GCCCATGATCCATTTTGGCAACGAC<br>TGGGAGGACCGCTACTACCGTGAA<br>AACATGTACCGCTACCCTAACCAA<br>GTGTACTACAGGCCAGTGGATCAG<br>TACAGCAACCAGAACAACTTCGTG<br>CACGACTGCGTCAATATCACCATC<br>AAGCAGCACACGGTCACCACCACC<br>ACCAAGGGGGAGAACTTCACCGAG<br>ACCGATGTGAAGATGATGGAGCGC<br>GTGGTGGAGCAGATGTGCGTCACC<br>CAGTACCAGAAGGAGTCCCAGGCC<br>TATTACGACGGGAGAAGATCCAGC<br>AGCACCGTGCTTTTCTCCTCCCCTC<br>CTGTCATCCTCCTCATCTCCTTCCTC<br>ATCTTCCTGATCGTGGGATGAGGG<br>AGGCCTTCCTGCTTGTTCCTTCGCA<br>TTCTCGTGGTCTAGGCTGGGGGAG<br>GGGTTATCCACCTGTAGCTCTTTCA<br>ATTGAGGTGGTTCTCATTCTTGCTT<br>CTCTGTGTCCCCCATAGGCTAATAC<br>CCCTGGCACTGATGGGCCCTGGGA<br>AATGTACAGTAGACCAGTTGCTCTT<br>TGCTTCAGGTCCCTTTGATGGAGTC<br>TGTCATCAGCCAGTGCTAACACCG<br>GGCCAATAAGAATATAACACCAAA<br>TAACTGCTGGCTAGTTGGGGCTTTG<br>TTTTGGTCTAGTGAATAAATACTGG<br>TGTATCCCCTGACTTGTACCCAGAG<br>TACAAGGTGACAGTGACACATGTA<br>ACTTAGCATAGGCAAAGGGTTCTA<br>CAACCAAAGAAGCCACTGTTTGGG<br>GATGGCGCCCTGGAAAACAGCCTC<br>CCACCTGGGATAGCTAGAGCATCC<br>ACACGTGGAATTCTTTCTTTACTAA<br>CAAACGATAGCTGATTGAAGGCAA<br>CAGGAAAAAAAAAAATCAAATTGTC<br>CTACTGACGTTGAAAGCAAACCTTT<br>GTTCATTCCCAGGGCACTAGAATG<br>ATCTTTAGCCTTGCTTGGATTGAAC<br>TAGGAGATCTTGACTCTGAGGAGA<br>GCCAGCCCTGTAAAAAGCTTGGTC<br>CTCCTGTGACGGGAGGGATGGTTA<br>AGGTACAAAGGCTAGAAACTTGAG<br>TTTCTTCATTTCTGTCTCACAATTAT<br>CAAAAGCTAGAATTAGCTTCTGCC<br>CTATGTTTCTGTACTTCTATTTGAA<br>CTGGATAACAGAGAGACAATCTAA |

| | | | | |
|---|---|---|---|---|
| | | | GATAAGAGAAGTAGGCTCCATTCC<br>AAAGTGGGAAAGAAATTCTGCTAG<br>CATTGTTTAAATCAGGCAAAATTTG<br>TTCCTGAAGTTGCTTTTTACCCCAG<br>CAGACATAAACTGCGATAGCTTCA<br>GCTTGCACTGTGGATTTTCTGTATA<br>GAATATATAAAACATAACTTCAAG<br>CTTATGTCTTCTTTTTAAAACATCT<br>GAAGTATGGGACGCCCTGGCCGTT<br>CCATCCAGTACTAAATGCTTACCGT<br>GTGACCCTTGGGCTTTCAGCGTGCA<br>CTCAGTTCCGTAGGATTCCAAAGC<br>AGACCCCTAGCTGGTCTTTGAATCT<br>GCATGTACTTCACGTTTTCTATATT<br>TGTAACTTTGCATGTATTTTGTTTT<br>GTCATATAAAAAGTTTATAAATGTT<br>TGCTATCAGACTGACATTAAATAG<br>AAGCTATGATGAACACCTGG | ACATTCTCTTAGGCTGCAGATAAG<br>AGAAGTAGGCTCCATTCCAAAGTG<br>GGAAAGAAATTCTGCTAGCATTGT<br>TTAAATCAGGCAAAATTTGTTCCTG<br>AAGTTGCTTTTTACCCCAGCAGACA<br>TAAACTGCGATAGCTTCAGCTTGCA<br>CTGTGGATTTTCTGTATAGAATATA<br>TAAAACATAACTTCAAGCTTATGTC<br>TTCTTTTTAAAACATCTGAAGTATG<br>GGACGCCCTGGCCGTTCCATCCAGT<br>ACTAAATGCTTACCGTGTGACCCTT<br>GGGCTTTCAGCGTGCACTCAGTTCC<br>GTAGGATTCCAAAGCAGACCCCTA<br>GCTGGTCTTTGAATCTGCATGTACT<br>TCACGTTTTCTATATTTGTAACTTT<br>GCATGTATTTTGTTTTGTCATATAA<br>AAAGTTTATAAATGTTTGCTATCAG<br>ACTGACATTAAATAGAAGCTATGA<br>TGAACACCTGG |
| ***Scg3*** | NM_011170.3 | *Mus musculus* | GGAGCATGCGCCTCCTCCTCATTTC<br>CTCTCCTGGAGCTAGTGGAGTAAA<br>GCTACGCCCAGGCCCGCGTCCGCT<br>GGCGGCGCAGGAACTTCAGCACCC<br>GCGGGGCGGACAGCGCCTACCGCA<br>CCTGCTCACCTGCTCTGGGCGCCAG<br>AAGAGCCTGCATCCTCCTTCCAGCC<br>CGGAGCAACTGCGCCGGAGGCGCC<br>CAGACCCTCTCCCTTCCCGCACCCA<br>GGCTCCTGTCCCTTCCAGCTTCTTA<br>ACTCCCCTTCTCATTCATAACAAAA<br>GCTACAGCTCAGGGGCCCAGCGCC<br>AAGCTCTTTCCAGCAAAGCACAGA<br>AGAGCAAGAAAGAATGGGGTTCCT<br>TTGGACCGGCTCTTGGATACTGGTG<br>TTGGTGCTCAACAGCGGCCCAATTC<br>AAGCTTTCCCCAAACCCGAAGGCA<br>GCCAAGACAAATCCCTGCATAATA<br>GAGAATTAAGTGCAGAAAGACCTT<br>TGAATGAACAGATCGCTGAGGCAG<br>AGGCAGACAAGATTAAAAAGGCAT<br>TCCCTTCAGAAAGCAAGCCGAGTG<br>AAAGCAATTATTCTTCTGTCGATAA<br>CTTGAATCTGCTGAGGGCAATAAC<br>AGAAAAGGAAACCGTTGAGAAAG<br>AGAGACAATCCATAAGAAGCCCCC<br>CGTTTGATAACCAACTGAACGTGG<br>AAGACGCTGATTCAACCAAAAATC<br>GGAAACTGATCGATGAGTACGATT<br>CCACCAAGAGTGGACTGGACCACA<br>AGTTTCAAGATGACCCAGACGGCC<br>TTCATCAACTGGATGGAACTCCTTT<br>AACTGCTGAAGACATCGTCCATAA<br>GATTGCCACCAGGATTTATGAGGA<br>GAACGACAGAGGAGTGTTTGACAA | **TAATACGACTCACTATAG**GGAGC<br>ATGCGCCTCCTCCTCATTTCCTCTC<br>CTGGAGCTAGTGGAGTAAAGCTAC<br>GCCCAGGCCCGCGTCCGCTGGCGG<br>CGCAGGAACTTCAGCACCCGCGGG<br>GCGGACAGCGCCTACCGCACCTGC<br>TCACCTGCTCTGGGCGCCAGAAGA<br>GCCTGCATCCTCCTTCCAGCCCGGA<br>GCAACTGCGCCGGAGGCGCCCAGA<br>CCCTCTCCCTTCCCGCACCCAGGCT<br>CCTGTCCCTTCCAGCTTCTTAACTC<br>CCCTTCTCATTCATAACAAAAGCTA<br>CAGCTCAGGGGCCCAGCGCCAAGC<br>TCTTTCCAGCAAAGCACAGAAGAG<br>CAAGAAAGAATGGGGTTCCTTTGG<br>ACCGGCTCTTGGATACTGGTGTTGG<br>TGCTCAACAGCGGCCCAATTCAAG<br>CTTTCCCCAAACCCGAAGGCAGCC<br>AAGACAAATCCCTGCATAATAGAG<br>AATTAAGTGCAGAAAGACCTTTGA<br>ATGAACAGATCGCTGAGGCAGAGG<br>CAGACAAGATTAAAAAGGCATTCC<br>CTTCAGAAAGCAAGCCGAGTGAAA<br>GCAATTATTCTTCTGTCGATAACTT<br>GAATCTGCTGAGGGCAATAACAGA<br>AAAGGAAACCGTTGAGAAAGAGA<br>GACAATCCATAAGAAGCCCCCCGT<br>TTGATAACCAACTGAACGTGGAAG<br>ACGCTGATTCAACCAAAAATCGGA<br>AACTGATCGATGAGTACGATTCCA<br>CCAAGAGTGGACTGGACCACAAGT<br>TTCAAGATGACCCAGACGGCCTTC<br>ATCAACTGGATGGAACTCCTTTAAC<br>TGCTGAAGACATCGTCCATAAGAT<br>TGCCACCAGGATTTATGAGGAGAA |

| | | | | |
|---|---|---|---|---|
| | | | AATTGTTTCTAAACTGCTGAATCTT<br>GGCCTGATCACTGAAAGCCAGGCA<br>CATACTCTGGAAGATGAAGTAGCA<br>GAAGCTTTACAAAAACTGATTTCA<br>AAAGAGGCCAACAATTATGAGGAG<br>ACCCTGGATAAACCCACAAGCAGG<br>ACCGAGAATCAGGATGGGAAAATA<br>CCAGAGAAAGTGACTCCGGTGGCA<br>GCAGTCCAAGATGGCTTCACTAAC<br>CGTGAAAACGATGAGACGGTGTCT<br>AACACCTTGACCTTGTCCAATGGCT<br>TGGAAAGGAGAACTAACCCCCACA<br>GGGAAGACGACTTTGAGGAACTCC<br>AGTATTTCCCCAACTTCTATGCACT<br>ACTGACAAGCATCGACTCAGAAAA<br>AGAAGCAAAAGAGAAAGAAACCC<br>TGATCACCATCATGAAGACATTGA<br>TTGACTTCGTGAAAATGATGGTGA<br>AATACGGTACGATATCTCCAGAGG<br>AAGGCGTGTCCTACCTTGAAAACTT<br>GGATGAAACAATTGCTCTGCAGAC<br>CAAGAACAAGCTAGAAAAAAATAC<br>TACTGATAGCAAAACTCCACCAGA<br>GAAGAGTCAGGAAGAAACAGACA<br>GTACCAAGGAAGAAGCCGCCAAGA<br>TGGAAAAGGAATACGGAAGCCTAA<br>AAGACTCTACAAAAGATGATAACT<br>CCAACCTAGGAGGAAAGACAGATG<br>AAGCCACAGGGAAGACAGAAGCCT<br>ACTTGGAAGCCATTAGAAAAAACA<br>TCGAATGGCTGAAGAAACATAACA<br>AGAAGGGCAACAAAGAAGATTACG<br>ACCTTTCAAAGATGAGGGACTTTAT<br>CAACCAACAAGCTGACGCTTATGT<br>GGAGAAGGGCATCCTCGACAAGGA<br>AGAAGCCAACGCCATCAAACGCAT<br>CTACAGCAGCCTGTGAAAATGGCG<br>GGCAGCTTGAGCCTTCCTGTTGTTC<br>CAGCAAAAACAATATAGCTTACAA<br>ACTAATTCGGCGGTTAAAGGGTTA<br>CCAGCCCAGAAGTATTAGGATGTG<br>CTGAATTTATAGTAGTTAATCCCTT<br>AGAAATGAGTAAAATAGAGCTCTC<br>TTGCCATAAATACCTTATGAAAAG<br>CAAAGCTGTAGAGAAGCCGAGGTT<br>TTTCTATATAGAATCCTTATTTCCT<br>CTTGAATTTACATTTTGTAATCAGA<br>GATGTGCTGCTCTGGAAAAGACTC<br>TAATGGGTTGAACATAAGTCTGAA<br>CCTACTCCCCACTGTCCTCAGCCCC<br>CTGAAGCTCTGAGAGGCCCTGTCTC<br>GGCATGCTAGACACCTGAGCACCT<br>CACTGGATGTTTGTCATAGGATGTC<br>GTTTCCACTAGTCGATCTCTGTTGG | CGACAGAGGAGTGTTTGACAAAAT<br>TGTTTCTAAACTGCTGAATCTTGGC<br>CTGATCACTGAAAGCCAGGCACAT<br>ACTCTGGAAGATGAAGTAGCAGAA<br>GCTTTACAAAAACTGATTTCAAAA<br>GAGGCCAACAATTATGAGGAGACC<br>CTGGATAAACCCACAAGCAGGACC<br>GAGAATCAGGATGGGAAAATACCA<br>GAGAAAGTGACTCCGGTGGCAGCA<br>GTCCAAGATGGCTTCACTAACCGT<br>GAAAACGATGAGACGGTGTCTAAC<br>ACCTTGACCTTGTCCAATGGCTTGG<br>AAAGGAGAACTAACCCCCACAGGG<br>AAGACGACTTTGAGGAACTCCAGT<br>ATTTCCCCAACTTCTATGCACTACT<br>GACAAGCATCGACTCAGAAAAAGA<br>AGCAAAAGAGAAAGAAACCCTGAT<br>CACCATCATGAAGACATTGATTGA<br>CTTCGTGAAAATGATGGTGAAATA<br>CGGTACGATATCTCCAGAGGAAGG<br>CGTGTCCTACCTTGAAAACTTGGAT<br>GAAACAATTGCTCTGCAGACCAAG<br>AACAAGCTAGAAAAAAATACTACT<br>GATAGCAAAACTCCACCAGAGAAG<br>AGTCAGGAAGAAACAGACAGTACC<br>AAGGAAGAAGCCGCCAAGATGGA<br>AAAGGAATACGGAAGCCTAAAAGA<br>CTCTACAAAAGATGATAACTCCAA<br>CCTAGGAGGAAAGACAGATGAAGC<br>CACAGGGAAGACAGAAGCCTACTT<br>GGAAGCCATTAGAAAAAACATCGA<br>ATGGCTGAAGAAACATAACAAGAA<br>GGGCAACAAAGAAGATTACGACCT<br>TTCAAAGATGAGGGACTTTATCAA<br>CCAACAAGCTGACGCTTATGTGGA<br>GAAGGGCATCCTCGACAAGGAAGA<br>AGCCAACGCCATCAAACGCATCTA<br>CAGCAGCCTGTGAAAATGGCGGGC<br>AGCTTGAGCCTTCCTGTTGTTCCAG<br>CAAAAACAATATAGCTTACAAACT<br>AATTCGGCGGTTAAAGGGTTACCA<br>GCCCAGAAGTATTAGGATGTGCTG<br>AATTTATAGTAGTTAATCCCTTAGA<br>AATGAGTAAAATAGAGCTCTCTTG<br>CCATAAATACCTTATGAAAAGCAA<br>AGCTGTAGAGAAGCCGAGGTTTTT<br>CTATATAGAATCCTTATTTCCTCTT<br>GAATTTACATTTTGTAATCAGAGAT<br>GTGCTGCTCTGGAAAAGACTCTAA<br>TGGGTTGAACATAAGTCTGAACCT<br>ACTCCCCACTGTCCTCAGCCCCCTG<br>AAGCTCTGAGAGGCCCTGTCTCGG<br>CATGCTAGACACCTGAGCACCTCA<br>CTGGATGTTTGTCATAGGATGTCGT<br>TTCCACTAGTCGATCTCTGTTGGGC |

| | | | | |
|---|---|---|---|---|
| | | | GCACGGAAATAAACCCACGTCTCT<br>TCATATCCTTGA | ACGGAAATAAACCCACGTCTCTTC<br>ATATCCTTGA |
| ***CLN33362*** | **AAS52304** | ***Ashbya gossypii*** | GGAGAGTCTGCATACCAAAGATCA<br>GCCGCTTGCATTGAAAGGGACGAA<br>CGGTGGCTCATGTCTTTGCCATCCT<br>TTCTTCTCCAATCCCCGAAGCACTT<br>CAGACACACACGTAACCTTGCATG<br>AAATATAGTATTTATTGTCATCGCC<br>GGCTGGACTGTTACTATCTCAAGCC<br>CAAAGACCATTGCATATCACACAC<br>AAATTCGTCACAAACACTTAAAGAT<br>TGTCCTGCGGTCAGGCTTCTCTGCT<br>ACACCGGGCCTCTGTCAGTATGTCT<br>GTTTGGGAACGTAAGTGTTCGCATC<br>ACCGGCTAATAAAAGTGGACGTGT<br>TTGCGTAACAAACTGACTGCTCGGA<br>GAACCGCATTTTAACCCTTACGCTT<br>CTGATTGATCTCGTGTATCACACCC<br>GGTAATCATGTACTTGTTTCCAGTC<br>CCGACCCTTAGTTACCGTGGCGATT<br>CGCAAATTGATTGATGATTCTTGCC<br>CTAGTTTCCCGCCATCCTTGTGCTTC<br>CCCTCTTCGACGCTCTTTATAGGCA<br>GCGCCAAATTTTATCGGTAGCGCGC<br>TATGTTCATTGTTATGTCACCCACA<br>TATCTTCTTCTTGGGCTTGACACTAT<br>TTGGCTGACGTCCAAGTTCGCAGAT<br>CTAATGCCCCGACTATCGAGCTTCC<br>AAAGCCTCCGCCGCCACGAGCAAA<br>AGAAACCGCATTTCACGACATCAG<br>AGCCGACAATCCCCAACACCCTTCA<br>CTGGTCAGTATGTACTACGGCGACT<br>CTCTATTCATAGGCGGAGATCAACC<br>GGCAAACAAAGATAGGCAGTCTCC<br>AGTTCTGGCCCATCTCTATGAGCAG<br>TTTAAGGTCTCGAGCACCGAATCTT<br>TGCAAGATATTCCCGTATGACCCAG<br>ACCCGAAATACAATACCAAGTTCA<br>GCGCCGTTCACTTCGCAAGCGTAAG<br>AGTTCTAAAGACCGTACGCAGGCTC<br>GCTGTGCTCTCCTAGGTCATGGTTT<br>GCTGACGATGTACAACTGACAAAA<br>ATCCCATTCCCATCCGCACCCAGGT<br>TCTAGAGGTAGTCGCCAACCCGTCC<br>AACGCAAACTCCAGATCTCCACCCA<br>ATTGCCTTTCAGGTGGTGCCTCGAT<br>CCTTGCCGTCTAAATTATAGCCAGC<br>CTGTGCCCATTTAACCGACGGTTAA<br>GTAATGAGAAAAAGCGTTCCTGGC<br>GAGATGCAGGCACGGGGGTGGGAA<br>TCGAATGGCCAGGCGCCGCAGTGC<br>TCCGTGCTACCTGCCTCTCCCCCGA | **TAATACGACTCACTATAG**GGAGA<br>GTCTGCATACCAAAGATCAGCCGCT<br>TGCATTGAAAGGGACGAACGGTGG<br>CTCATGTCTTTGCCATCCTTTCTTCT<br>CCAATCCCCGAAGCACTTCAGACAC<br>ACACGTAACCTTGCATGAAATATAG<br>TATTTATTGTCATCGCCGGCTGGAC<br>TGTTACTATCTCAAGCCCAAAGACC<br>ATTGCATATCACACACAAATTCGTC<br>ACAAACACTTAAAGATTGTCCTGCG<br>GTCAGGCTTCTCTGCTACACCGGGC<br>CTCTGTCAGTATGTCTGTTTGGGAA<br>CGTAAGTGTTCGCATCACCGGCTAA<br>TAAAAGTGGACGTGTTTGCGTAACA<br>AACTGACTGCTCGGAGAACCGCATT<br>TTAACCCTTACGCTTCTGATTGATC<br>TCGTGTATCACACCCGGTAATCATG<br>TACTTGTTTCCAGTCCCGACCCTTA<br>GTTACCGTGGCGATTCGCAAATTGA<br>TTGATGATTCTTGCCCTAGTTTCCC<br>GCCATCCTTGTGCTTCCCCTCTTCG<br>ACGCTCTTTATAGGCAGCGCCAAAT<br>TTTATCGGTAGCGCGCTATGTTCAT<br>TGTTATGTCACCCACATATCTTCTTC<br>TTGGGCTTGACACTATTTGGCTGAC<br>GTCCAAGTTCGCAGATCTAATGCCC<br>CGACTATCGAGCTTCCAAAGCCTCC<br>GCCGCCACGAGCAAAAGAAACCGC<br>ATTTCACGACATCAGAGCCGACAAT<br>CCCCAACACCCTTCACTGGTCAGTA<br>TGTACTACGGCGACTCTCTATTCAT<br>AGGCGGAGATCAACCGGCAAACAA<br>AGATAGGCAGTCTCCAGTTCTGGCC<br>CATCTCTATGAGCAGTTTAAGGTCT<br>CGAGCACCGAATCTTTGCAAGATAT<br>TCCCGTATGACCCAGACCCGAAATA<br>CAATACCAAGTTCAGCGCCGTTCAC<br>TTCGCAAGCGTAAGAGTTCTAAAG<br>ACCGTACGCAGGCTCGCTGTGCTCT<br>CCTAGGTCATGGTTTGCTGACGATG<br>TACAACTGACAAAAATCCCATTCCC<br>ATCCGCACCCAGGTTCTAGAGGTAG<br>TCGCCAACCCGTCCAACGCAAACTC<br>CAGATCTCCACCCAATTGCCTTTCA<br>GGTGGTGCCTCGATCCTTGCCGTCT<br>AAATTATAGCCAGCCTGTGCCCATT<br>TAACCGACGGTTAAGTAATGAGAA<br>AAAGCGTTCCTGGCGAGATGCAGG<br>CACGGGGGTGGGAATCGAATGGCC<br>AGGCGCCGCAG**T**GCTCCGTGCTACC |

| | | | | |
|---|---|---|---|---|
| | | | TATCAGCAGTGACGGCGACTCTAGC<br>TCGGGCTCTGTTCCACCATTTACTC<br>TGGACCGGCCCGCCCCGCACAAGA<br>CGACACTCAAGACAGCTGTGGCCC<br>CCCTAACATCTACTCCGAACGACTC<br>ACCGCGCACTCATCTCACGTATCGT<br>TCGATGCCCCAGACGCCACCCACTA<br>TTATGTCCGTTGCAGCTCAGGCGCG<br>TGCACAATGCCTCCCAGACCTACAC<br>TCCACCACGACTAAGAACCCCCTTC<br>ACTTGCTGCATTCTGAAGAATCTAA<br>GGACTCCGCCACGGGTATCGACCAT<br>GAGCTATGCGACATTGTTCAGGACT<br>CGGTCAAGAAACTACGCTAGCATA<br>C | TGCCTCTCCCCCGATATCAGCAGTG<br>ACGGCGACTCTAGCTCGGGCTCTGT<br>TCCACCATTTACTCTGGACCGGCCC<br>GCCCCGCACAAGACGACACTCAAG<br>ACAGCTGTGGCCCCCCTAACATCTA<br>CTCCGAACGACTCACCGCGCACTCA<br>TCTCACGTATCGTTCGATGCCCCAG<br>ACGCCACCCACTATTATGTCCGTTG<br>CAGCTCAGGCGCGTGCACAATGCCT<br>CCCAGACCTACACTCCACCACGACT<br>AAGAACCCCCTTCACTTGCTGCATT<br>CTGAAGAATCTAAGGACTCCGCCA<br>CGGGTATCGACCATGAGCTATGCG<br>ACATTGTTCAGGACTCGGTCAAGAA<br>ACTACGCTAGCATAC |
| ***CLN3657*** | AAS52304 | *Ashbya gossypii* | GGAGAGTCTGCATACCAAAGATCA<br>GCCGCTTGCATCACATGAGACCCTG<br>CGGCACTCTCTTATTTTGCGTAGTTT<br>CTTGGCCAAACTTAGGTAACCCACA<br>CTAAGAGGACACTGCCTGCATAAA<br>TACTGAAATTCTCTGTTCTCATCAT<br>CCGAGCAGTTAATAACTCCTAACGG<br>ACCCCATTTGCATCTTATTCGCACA<br>TCTGCGAAAGACTCTAGGAAACAC<br>CCACGCTCTAAAGCATCAATACTCC<br>TATTGTCCGTAAATGGTACATCCGT<br>GTGGGAAAGTCACGCGCTCCATCG<br>CTGGTTCCAGACCAGAACGATGGA<br>AGCGCATCAATATGCTTGCCAGCAT<br>GCCAGTGTTGCACTCCGCAAGCATC<br>TCAGTGATCTGGAGAACCAGAAGC<br>GGGCGTCCGGGACTGGCTTCCCGTA<br>CCTGCCGCAACTAGGTATCGGGATG<br>TTCAACTTGAGTTTGCGCCACTTCA<br>TGTCCTTCCAAACTCGCTTGCGACT<br>GTCCTAATCCACTCGCCTATCATGG<br>CAAGACATCATGGGGCTTTTCAGCT<br>CCAAGATGGTTTTTACTTCTACCCC<br>CTTTCACCTACATACGCTTACGGCG<br>GTAAGCCTGTAGTGATAGTTCTTAC<br>ATAAAAACACCCGCACCCCGAGCC<br>CGCAGCGCAATTGCTGCGACAACC<br>GCTCACATGGGCGGTACCATGACCC<br>GGAGCTGACCAACTTCTAGACGCTT<br>CCCTGACCTGCGTATTCAGCTCCGA<br>GTCATGTTTGCGCGCTTCATAACCT<br>ACAGATAACCGAGACCCCCAAGCC<br>GTGCTCCATGCTCCTTAATCGAAAC<br>AGTTGTGAGCATGGAGCAGGCATA<br>ATTGGGTCAGTATGCAGCTCTTACC<br>CAGGAGTTTACTATCATAACTGCTC<br>CAACGCCATTCATATGGCGAGCGTT<br>ACCCCTATAACAATCGCATCCCGAT<br>ATGCTGACCCCAAAGGGTCAGCAT | **TAATACGACTCACTATAG**GGAGA<br>GTCTGCATACCAAAGATCAGCCGCT<br>TGCATCACATGAGACCCTGCGGCAC<br>TCTCTTATTTTGCGTAGTTTCTTGGC<br>CAAACTTAGGTAACCCACACTAAG<br>AGGACACTGCCTGCATAAATACTG<br>AAATTCTCTGTTCTCATCATCCGAG<br>CAGTTAATAACTCCTAACGGACCCC<br>ATTTGCATCTTATTCGCACATCTGC<br>GAAAGACTCTAGGAAACACCCACG<br>CTCTAAAGCATCAATACTCCTATTG<br>TCCGTAAATGGTACATCCGTGTGGG<br>AAAGTCACGCGCTCCATCGCTGGTT<br>CCAGACCAGAACGATGGAAGCGCA<br>TCAATATGCTTGCCAGCATGCCAGT<br>GTTGCACTCCGCAAGCATCTCAGTG<br>ATCTGGAGAACCAGAAGCGGGCGT<br>CCGGGACTGGCTTCCCGTACCTGCC<br>GCAACTAGGTATCGGGATGTTCAAC<br>TTGAGTTTGCGCCACTTCATGTCCT<br>TCCAAACTCGCTTGCGACTGTCCTA<br>ATCCACTCGCCTATCATGGCAAGAC<br>ATCATGGGGCTTTTCAGCTCCAAGA<br>TGGTTTTTACTTCTACCCCCTTTCAC<br>CTACATACGCTTACGGCGGTAAGCC<br>TGTAGTGATAGTTCTTACATAAAAA<br>CACCCGCACCCCGAGCCCGCAGCG<br>CAATTGCTGCGACAACCGCTCACAT<br>GGGCGGTACCATGACCCGGAGCTG<br>ACCAACTTCTAGACGCTTCCCTGAC<br>CTGCGTATTCAGCTCCGAGTCATGT<br>TTGCGCGCTTCATAACCTACAGATA<br>ACCGAGACCCCCAAGCCGTGCTCC<br>ATGCTCCTTAATCGAAACAGTTGTG<br>AGCATGGAGCAGGCATAATTGGGT<br>CAGTATGCAGCTCTTACCCAGGAGT<br>TTACTATCATAACTGCTCCAACGCC<br>ATTCATATGGCGAGCGTTACCCCTA<br>TAACAATCGCATCCCGATATGCTGA |

| | | | | |
|---|---|---|---|---|
| | | | GCTGCAGGCCGTGTTTAACAGATGG<br>AAATCTCATGGCGGTCTGCAGCCCC<br>CATCTTTGCCTACAGGCCTGCGATG<br>TCAAATTCCGGGCCTGTCTTTCCCT<br>CTAGTCGGGATCCTGGAGCAGCGT<br>GCGAATCTAAAGCACGACTATGTCT<br>CCTCGGTAACCTTAATACCGCCAGT<br>TAGCCGATGAGATCAATCTGTAGTA<br>TGGTTATGCAAGCGCTTCCGTGGGC<br>TCGCCAAGTGCGGGTCTAGCTCCGG<br>CCACTGCGGCCAGCGGCTCTCCCGC<br>TGTCCGCATGGCAGGCAGCTAGAC<br>CCCGGCCTATGTCCCGACATCCGCT<br>CTGGTGACCCTTACGCGGCATATGC<br>CGACGATCAAGGCTTTTATGCCGCT<br>CACTCCAACTACTTCTCGCAAGCCA<br>AAGCGCCTAGTGCTCTGGCTTAGTG<br>AAAAGTACCAGACGAGGCCCGCGC<br>CTGTGGAGGTTACATCATAGGCGCG<br>CGCAAAAGGCACTACCCATCAACA<br>TTCACCGCCTCTCCAGGAGATCGCG<br>CAGTTGCTGCATCAGTAGCAACATC<br>AGCGCGACCCCATGGCCGTGGACC<br>TCCAGTTCAATGCCCGGGAACCACT<br>CCTGGTCAAGAAACTACGCTAGCAT<br>AC | CCCCAAAGGGTCAGCATGCTGCAG<br>GCCGTGTTTAACAGATGGAAATCTC<br>ATGGCGGTCTGCAGCCCCCATCTTT<br>GCCTACAGGCCTGCGATGTCAAATT<br>CCGGGCCTGTCTTTCCCTCTAGTCG<br>GGATCCTGGAGCAGCGTGCGAATC<br>TAAAGCACGACTATGTCTCCTCGGT<br>AACCTTAATACCGCCAGTTAGCCGA<br>TGAGATCAATCTGTAGTATGGTTAT<br>GCAAGCGCTTCCGTGGGCTCGCCAA<br>GTGCGGGTCTAGCTCCGGCCACTGC<br>GGCCAGCGGCTCTCCCGCTGTCCGC<br>ATGGCAGGCAGCTAGACCCCGGCC<br>TATGTCCCGACATCCGCTCTGGTGA<br>CCCTTACGCGGCATATGCCGACGAT<br>CAAGGCTTTTATGCCGCTCACTCCA<br>ACTACTTCTCGCAAGCCAAAGCGCC<br>TAGTGCTCTGGCTTAGTGAAAAGTA<br>CCAGACGAGGCCCGCGCCTGTGGA<br>GGTTACATCATAGGCGCGCGCAAA<br>AGGCACTACCCATCAACATTCACCG<br>CCTCTCCAGGAGATCGCGCAGTTGC<br>TGCATCAGTAGCAACATCAGCGCG<br>ACCCCATGGCCGTGGACCTCCAGTT<br>CAATGCCCGGGAACCACTCCTGGTC<br>AAGAAACTACGCTAGCATAC |

**Supplementary Table S4: Atp5b_DNA oligos sequence for smFISH.**

| Table S4_Atp5b_DNA oligos sequence for smFISH | | |
|---|---|---|
| NAME_start | Sequence (5'->3') | position of the first nucleotide |
| Atp5b_p39 | GCCCCACAAGACTCAACA | 39 |
| Atp5b_p160 | GCATAGTCTCTGGCAGGA | 160 |
| Atp5b_p275 | GGCATTTAGGATGGGTGG | 275 |
| Atp5b_p354 | CAATAGTTCTGACCGTGCTC | 354 |
| Atp5b_p375 | AGCCTTCAGTGCCATCCA | 375 |
| Atp5b_p399 | CCAGTACTTTCTGGCCTC | 399 |
| Atp5b_p424 | GGAATTTTGATTGGTGCCCC | 424 |
| Atp5b_p490 | GGACCTCTCTCATCAATAGG | 490 |
| Atp5b_p545 | CTCTATGAACTCAGGAGCC | 545 |
| Atp5b_p577 | CCAGTCACCAGAATCTCC | 577 |
| Atp5b_p625 | ATTTTCCCACCCTTGGCG | 625 |
| Atp5b_p675 | GCTCCATGATCAGTACTGTC | 675 |
| Atp5b_p756 | AATCATTGCCCTCACGGG | 756 |
| Atp5b_p831 | GTCCATATACCAACGCTACC | 831 |
| Atp5b_p880 | AAACCAGTCAGAGCTACCC | 880 |
| Atp5b_p960 | CCTGGGTAAAGCGGAAGA | 960 |
| Atp5b_p988 | CCCAATAAGGCAGACACC | 988 |
| Atp5b_p1011 | AGCCTACAGCAGAAGGGA | 1011 |
| Atp5b_p1066 | GTGGTGGTGATCCTTTCC | 1066 |
| Atp5b_p1111 | GCAGGCACATAGATAGCC | 1111 |
| Atp5b_p1162 | GCATCCAAATGGGCAAAGG | 1162 |
| Atp5b_p1210 | GCTGGATAGATGCCCAAC | 1210 |
| Atp5b_p1326 | CCTGGAGAGATTTGTAGTCC | 1326 |
| Atp5b_p1453 | ATGTGACCCGTGAAGACC | 1453 |
| Atp5b_p1537 | GCTTGTTCTGGGAGATGG | 1537 |
| Atp5b_p1561 | TCAATGGGTCCCACCATG | 1561 |
| Atp5b_p1648 | GAGAGATCAGTTGCAGTGC | 1648 |
| Atp5b_p1678 | GGAAGGAAACCTGAGCTC | 1678 |
| Atp5b_p1701 | CAAGGCTCTTGTGTGGCC | 1701 |
| Atp5b_p1831 | CTAAATGTGGCCCACCAC | 1831 |

**Supplementary Table S5: Antibody used in this study.**

| **Table S5_Antibodies in use** | | | | | |
|---|---|---|---|---|---|
| **Name** | **Type** | **Concentration in use in this study** | **Application in this study** | **Source** | **Identifier** |
| **anti-vGLUT1** | polyclonal rabbit | 1/500 | ICC in WT neurons | Takamori et al., 2001 | Shigeo3 |
| **anti-MAP2** | polyclonal chicken | 1/1000 | ICC in WT neurons | kind gift from Suzanne Wegmann's lab | n/a |
| **anti-EIF4E** | monoclonal rabbit | 1/250 | ICC in HEK293 | Abcam | ab33766 |
| **anti-rabbit STAR 488** | goat anti-rabbit | 1/500 | ICC in HEK293 | Aberior | ab464 |
| **anti-chicken AF488** | anti-chicken | 1/500 | ICC in WT neurons | kind gift from Suzanne Wegmann's lab | n/a |
| **anti-rabbit AF568** | goat anti-rabbit | 1/500 | ICC in WT neurons | Alexa Fluor | A-11001 |
| **anti-betaActin** | monoclonal mouse | 1/2000 | WB | Sigma Aldrich | A5441-0.2ML |
| **anti-mCherry** | monoclonal mouse | 1/2000 | WB | Abcam | Ab125096 |
| **anti-mouse HRP** | anti-mouse IgG | 1/10000 | WB | Sigma Aldrich | A4416-1ML |
